# Redirecting DNA repair for efficient CRISPR-Cas-based gene targeting in tomato

**DOI:** 10.1101/2024.03.12.584635

**Authors:** Tien Van Vu, Ngan Thi Nguyen, Jihae Kim, Minh Huy Vu, Young Jong Song, Mil Thi Tran, Yeon Woo Sung, Jae-Yean Kim

## Abstract

The CRISPR-Cas-based gene targeting (GT) method has enabled precise modifications of genomic DNA ranging from single base to several kilobase scales through homologous recombination (HR). In plant somatic cells, canonical nonhomologous end-joining (cNHEJ) is the predominant mechanism for repairing double-stranded breaks (DSBs), thus limiting the HR-mediated GT. In this study, we implemented various approaches to shift the repair pathway preference toward HR by using a dominant-negative KU80 mutant protein (KUDN) to disrupt the initiation of cNHEJ and enhance DSB end resection through nucleases. Our results show from 1.71- to 3.55-fold improvement of the GT efficiency at the callus stage and a more remarkable, up to 9.84-fold, increase in GT efficiency at two specific tomato loci, *SlHKT1;2* and *SlEPSPS1*, when we screened transformants obtained from the KUDN-mediated cNHEJ suppression approach. With practical levels of efficiency, this enhanced KUDN-based GT tool successfully facilitated GT at an additional locus, *SlCAB13*. These findings provide a promising method for more efficient and precise plant breeding in the future.

## INTRODUCTION

Gene targeting (GT) was initially designed to replace a genomic DNA sequence with exogenous DNA donors through the homologous recombination (HR) mechanism (Paszkowski et al., 1988). GT is one of the few methods capable of precisely editing genes of interest across scales ranging from single base pairs to kilo bases while avoiding any undesirable alterations to the genome (Puchta, 2005; Vu et al., 2019). While the prime editing technique shows promise for similar precision, it remains confined to small-scale DNA modifications without any scar (Chen and Liu, 2023; Vu et al., 2024). GT holds the potential to meticulously replace entire genes or alleles or enable complex edit installation, offering a valuable strategy for gene/allele pyramiding in precision plant breeding (Vu et al., 2019). In plants, the GT efficiency has been notably low for practical applications (Puchta, 2005; Vu et al., 2020).

Two major strategies that could enhance the GT efficiency in plants are (1) leveraging artificial DNA double-stranded breaks (DSBs) and (2) introducing a substantial quantity of DNA donors (Baltes et al., 2014; Cermak et al., 2015; Puchta et al., 1993). The efficiency of HR has been boosted from 10 to 100 times by employing the meganuclease I-Sce I to induce site-specific DSBs (Puchta et al., 1996). Recently, various approaches have been presented to improve GT rates. Firstly, molecular scissors such as the Clustered Regularly Interspaced Short Palindromic Repeats (CRISPR)-CRISPR-associated protein (Cas) are utilized to induce DSBs at the intended target sequences (Belhaj et al., 2013; Cermak et al., 2015; Voytas, 2013; Vu et al., 2020). Secondly, the frequency of HR is heightened by incorporating a viral replicon vehicle for donor templates (Baltes et al., 2014; Cermak et al., 2015; Hummel et al., 2018; Vu et al., 2020). Likewise, several novel approaches have been reported in supporting elevated GT efficiency in mammals and plants, including the facilitating the HR pathway choice via suppression of the canonical nonhomologous end-joining (cNHEJ) by chemical (Chu et al., 2015; Vu et al., 2021a) or biological agents (Endo et al., 2016; Movahedi et al., 2022; Nishizawa-Yokoi et al., 2012; Qi et al., 2013; Wu et al., 2022) and facilitation of DSB end resection by introducing DSB end processing enzymes (Charpentier et al., 2018; Park et al., 2021).

Previously, we established a GT system based on a geminiviral replicon and the CRISPR-LbCas12a nuclease (Vu et al., 2020) that was improved by using the temperature-tolerant (ttLbCas12a) variant (Huang et al., 2021; Merker et al., 2020b; Schindele et al., 2023; Vu et al., 2021a) and further enhanced by chemical treatments for cNHEJ suppression (Vu et al., 2021a). However, it is still challenging to conduct GT experiments without allele-associated markers for selecting GT events, which limits the applications of GT in tomato and other plants. We hypothesize that our GT system can be further improved by updating with the recent advancements in favoring the HR choice and enhancing DSB resection in HR reaction. Our data indicates that the GT efficiency was enhanced up to 3.55 folds and 9.84 folds at the callus and plant stages, respectively, using the improved KUDN-based GT tool and was applicable to other loci at practical levels, though its efficiency is still genomic context-dependent.

## RESULTS

### Employing end resection-related enzymes led to an altered repaired pathway profile

The end resection of the DSBs is prerequisite for generating 3’ ssDNA overhangs that can be commonly used for a homology-directed repair (HDR) process, such as microhomology-mediated end joining (MMEJ), single-strand annealing (SSA), or HR, depending on the levels of end resection(Symington and Gautier, 2011; Truong et al., 2013). Therefore, adding DSB end processing enzymes to the cleavage sites is expected to help enhance GT efficiency. Previously reported end resection enzymes and approaches such as hExo1(Hackley, 2021), hHE, CtIP (Charpentier et al., 2018) and T5 exonuclease (T5exo) (Zhang et al., 2020) together with the CRISPR-Cas9 or Cas12a system significantly enhanced homologous recombination-mediated gene editing. We designed and tested GT tools using the end resection enzyme by fusing them to the ttLbCas12a protein (Figure S1) for carrying them to the target sites. The targeted deep sequencing data obtained from analyzing 10-day post-transformation (dpt) explants showed slight improvement of GT with T5exo-ttLbCas12a and ttLbCas12a-CtIP fusions at both *SlHKT1;2* and *SlEPSPS1* sites (Figure S2a). On the other hand, the hExo1-ttLbCas12a and ttLbCas12a-hHE fusions led to a slight reduction of GT efficiency. However, none of the end resection enzyme fusions resulted in a significant enhancement of GT efficiency. Assessing the indel mutation efficiency among the fusions revealed significant reduction by the hExo1-ttLbCas12a and ttLbCas12a-CtIP compared to the ttLbCas12a control at the *SlEPSPS1* site (Figure S2b) that could be partially explained by the low protein accumulation levels of the fusions (Figure S3). By contrast, adding T5exo to the N-terminal of ttLbCas12a slightly enhanced the indel mutation efficiency (Figure S2b) that might be supported by the higher protein level of the T5exo-ttLbCas12a fusion (Figure S3). Further analysis revealed the enhancement of intramolecular MMEJ frequency using 2-or 4-nt microhomology (MH) at the *SlHKT1;2* but not *SlEPSPS1* site with T5exo, hHE and CtIP compared to the control (Figure S2c and Table S1). Interestingly, the CtIP dramatically enhanced 4-nt MH MMEJ frequency (Figure S2c) indicating a shift in end resection length with CtIP.

### NKUDN enhanced GT efficiency

The Ku70/80 complex was shown to play roles in early sensing DSBs, binding and activating the cNHEJ pathway (Symington and Gautier, 2011). Knocking out or blocking KU70/80 complex by chemicals led to the enhancement of the alternative pathways, including the HR (Chu et al., 2015; Nishizawa-Yokoi et al., 2012; Schmidt et al., 2019; West et al., 2002; Wu et al., 2022). A truncated Ku80(449–732) fragment showed a dominant-negative activity that interferes with the NHEJ pathway in hamster (Marangoni et al., 2000). We identified the tomato *KU80* homolog (Solyc01g091350) and a putative SlKu80DN (427-709) fragment (named as KUDN) by NCBI BLAST (Figure S4). We hypothesized that this KUDN could increase the HR-based GT of the plant. By fusing KUDN to ttLbCas12a at the N-terminal (termed NKUDN) and C-terminal (CKUDN), the KUDN peptide is directly located to the targeted sites for effective actions.

We observed the improvement of HR-based GT efficiency at the callus stage revealed by targeted NGS using NKUDN but not CKUDN configuration (Figure 1b). More importantly, NKUDN significantly enhanced the GT efficiency at the *SlEPSPS1* locus compared to the construct without KUDN (0.270±0.103% vs. 0.076±0.024%, a 3.55-fold increase). NKUDN also helped to improve GT at the *SlHKT1;2* site from 0.075±0.017% to 0.128±0.037%, a 1.71-fold enhancement (Figure 1b). The GT efficiency reached up to 0.199% and 0.471% at the *SlHKT1;2*, and *SlEPSPS1* sites, respectively, with the involvement of the KUDN-ttLbCas12a module (Figure 1b). This data indicates that NKUDN positively impacted GT, albeit the GT efficiency differed among different targeted sites. Subsequently, we found that NKUDN caused higher indel mutation efficiency to both the *SlHKT1;2* and *SlEPSPS1* targets (Figure 1c).

**Figure 1.**
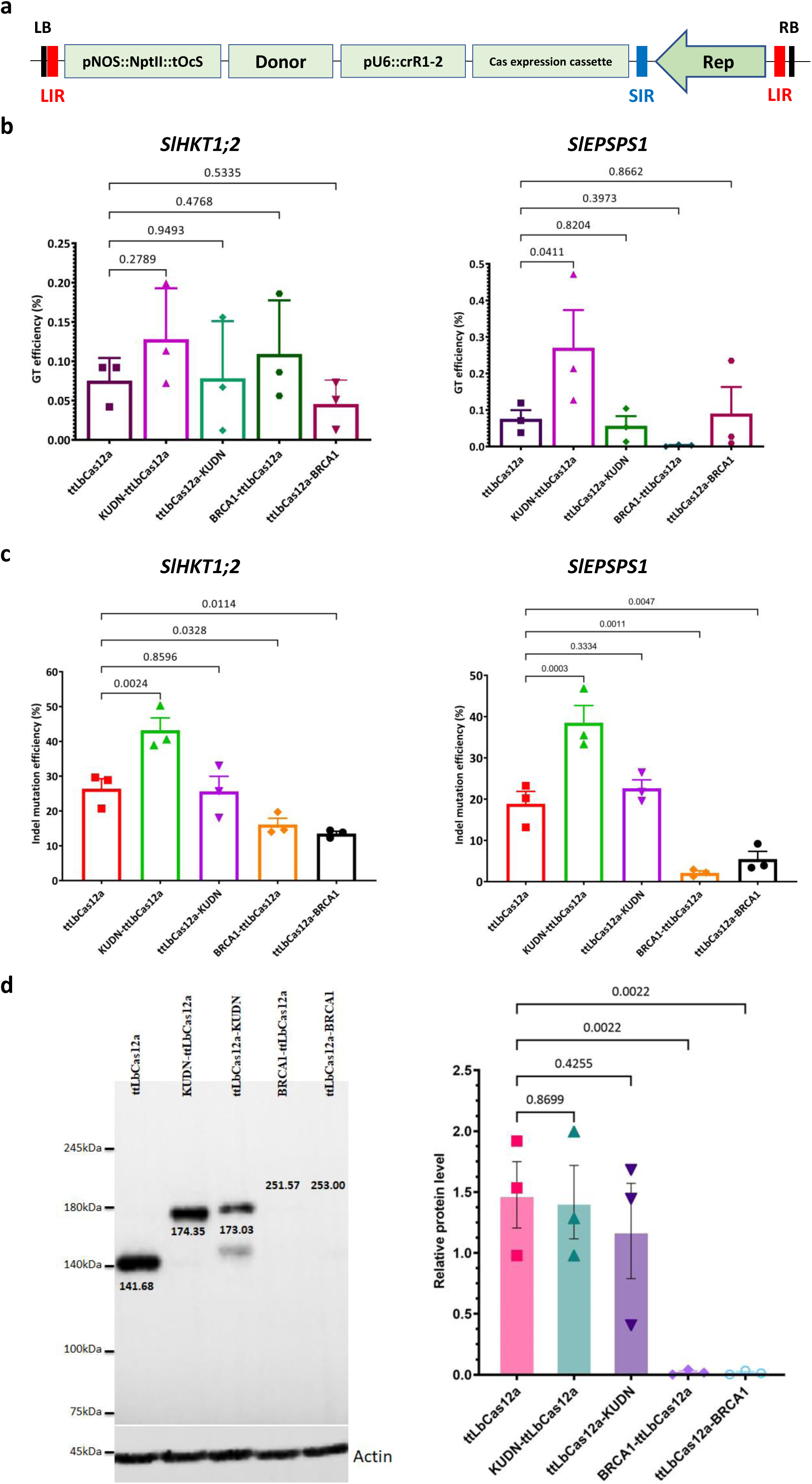
Impacts of the factors that affect DSB repair pathway choice on the editing efficiency in tomato. (**a**) Plasmid map of the GT tool used in the study. The GT construct was cloned into a geminiviral replicon vector with the boundaries of two long intergenic sequences (LIRs) and one short intergenic sequence (SIR). The replicon is autonomously replicated with the support of the Rep protein/expression cassette. (**b-c**) The impacts of KUDN and BRCA1 on GT (**b**) and indel mutation (**c**) efficiency at the callus stage. The efficiencies were assessed by targeted NGS using 21-dpt cotyledon/callus samples. (**d**) The protein expression levels of ttLbCas12a and its KUDN and BRCA1 fusions. Western blot membranes showing bands of the Cas proteins with expected sizes and expression cassettes. Actin levels were used as loading controls. P-values of the t-test for pair-wise comparison between the relative protein levels of ttLbCas12a and the fusions are indicated on the top of the bars. The data points are shown on the plots. The expression cassettes and Cas configurations are at the bottom of the bars.

We reasoned that the fusions of KUDN to ttLbCas12a could alter the protein’s expression levels thereby affecting the GT efficiency. Using Western blot analysis, we found that the relative protein levels of the fusions were not significantly changed between the ttLbCas12a (1.478±0.273) and KUDN-ttLbCas12a (1.418±0.301) (Figure 1d). This indicates that the enhanced GT and indel mutation of the NKUDN could be attributed to the KUDN activity.

### The KUDN involvement led to a notable increase MMEJ repair

A DSB in DNA can be repaired through two main pathways: cNHEJ and HDR. Among the sub-pathways of HDR, MMEJ, also known as alternative NHEJ, is an error-prone repair mechanism that requires short microhomologies (MHs) flanking the DSB ends (Truong et al., 2013). These sequences anneal to seal the break, causing a deletion mutation that loses a MH and the sequence between the MHs (Vu et al., 2021b). The MMEJ repair mechanism shares a common initiation step with HR, which requires the redirection of the DSB repair pathway choice from cNHEJ to HDR by initiating DSB end resection (Sfeir and Symington, 2015; Truong et al., 2013; Vu et al., 2021a). To determine if the addition of KUDN could impair cNHEJ and enhance MMEJ, we analyzed the MMEJ traces from the targeted NGS data.

The NKUDN alone improved MMEJ frequency at 1.31-2.00 folds at the *SlHKT1;2* and *SlEPSPS1* sites. Also, the usage of 2-nt MHs was increased with NKUDN at both the targeted sites (Figure 2b). Importantly, it was observed that the MMEJ frequency was highest in the construct containing KUDN-ttLbCas12a (as shown in Figure 2a, b). This indicates that the 2-nt MH-mediated DSB repair was enhanced, resulting in higher indel mutations, and shifted to the HDR choice.

**Figure 2.**
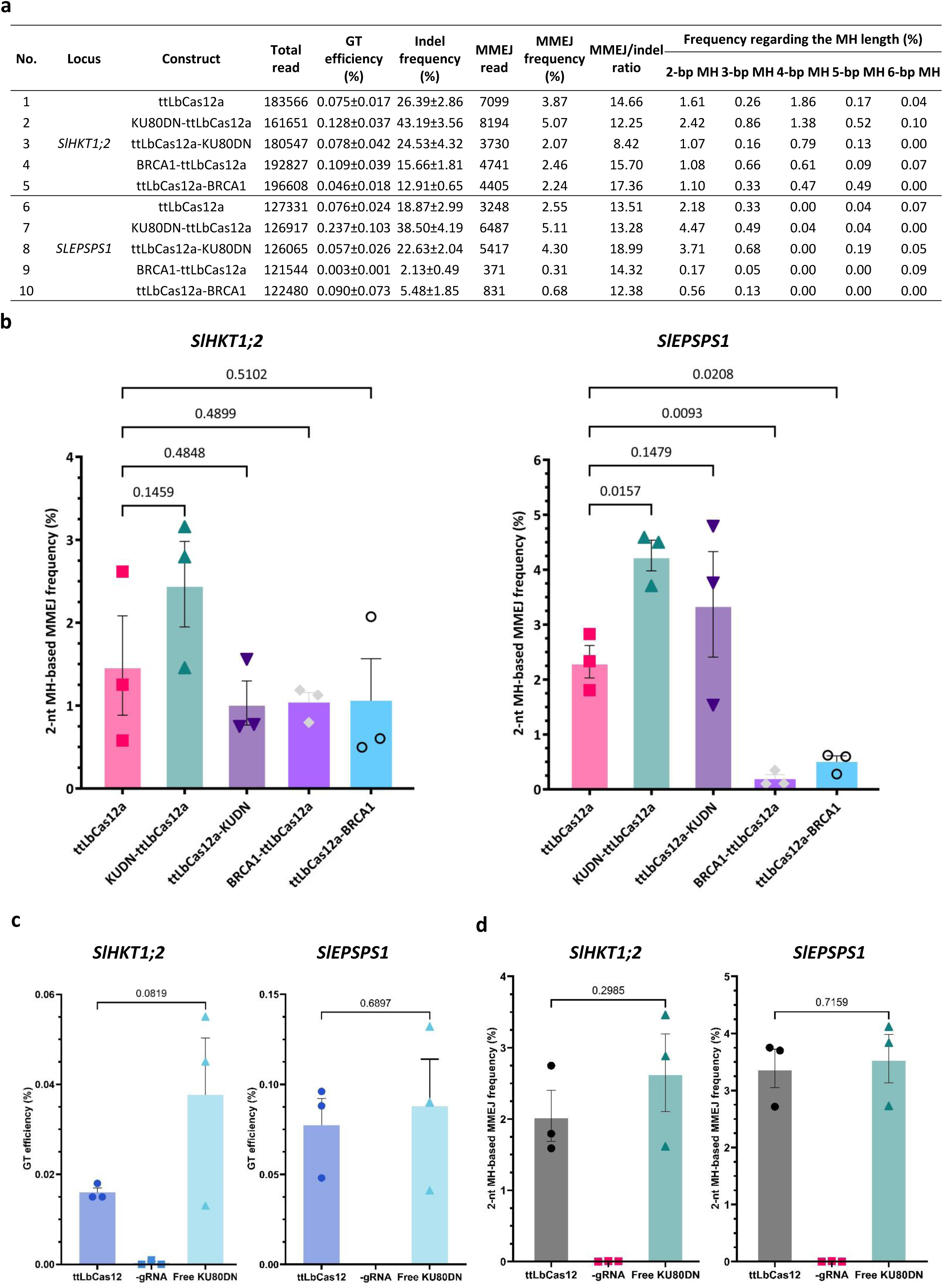
The impacts of KUDN and BRCA1 on the intramolecular MMEJ patterns at the *SlHKT1;2* and *SlEPSPS1* sites. (**a**) Table summarizing the impacts of microhomology lengths and MMEJ frequencies. (**b**) The 2-nt microhomology-based MMEJ frequency. The indel and MMEJ frequencies were measured by targeted NGS. (**c-d**) Impacts of free KUDN on GT (**c**) and 2-nt MH MMEJ (**d**) frequency. The data points are shown on the plots.

The NKUDN fusion alone improved the MMEJ frequency by 1.31-2.00 folds. Furthermore, the use of 2-nt MHs was increased with NKUDN at both the targeted sites (Figure 2b). Importantly, we found that, as observed with the GT outcomes (Figure 1b), the MMEJ frequency was highest in the construct containing KUDN-ttLbCas12a (as shown in Figure 2a, b).

### Free forms of KUDN showed comparable or reduction of GT performance

We have observed positive effects of KUDN fusions on DSB repair choice towards HDR (Figure 1, 2). We then explored whether similar or more pronounced effects could be achieved using free form KUDN. Our experiments showed a slight improvement in the GT efficiency at the *SlHKT1;2* site and a comparable performance at the *SlEPSPS1* when KUDN was overexpressed alongside the GT tools (Figure 2c). Additionally, we found that the GT performance was correlated with the efficiency of indel mutation and MMEJ (Figure 2d, Figure S5a, b, and Table S2).

We also used the Suntag system(Tanenbaum et al., 2014) to recruit KUDN to the targeted site. We added 10x Suntag peptide epitopes (GCN4_v4) to the C-terminal of ttLbCas12a, and expressed a fusion of scFv-KUDN in parallel (Supplemental sequences). However, the recruitment system unexpectedly led to a reduction of GT efficiency at both targeted sites (Figure S5c and Table S3), possibly due to a correlated reduction of indel mutation efficiency (Figure S5d).

### BRCA1 fusions to ttLbCas12a reduced the editing activity of the GT tools

BRCA1 plays a crucial role in the HR pathway by interfering with 53BP1 accumulation at the DSB site, enabling end resection and redirecting the repair pathway choice towards HR (Prakash et al., 2015). Although putative BRCA1 homologs exist in plants, their functional roles are not fully understood yet(Trapp et al., 2011). In this study, we identified a tomato homolog (Solyc08g023280) of the AtBRCA1 protein and employed it to engineer GT tools to carry BRCA1 to targeted sites from the N-(NBRCA1) or C-terminal (CBRCA1) of ttLbCas12a (Figure S1b). We observed reduced GT and indel mutation efficiencies of the BRCA1-based GT tools at both targeted sites (Figure 1a, b), along with weaker MMEJ patterns (Figure 2a, b). Moreover, the GT, indel, and MMEJ frequency was dramatically reduced at the *SlEPSPS1* site (Figure 1b, c and 2a, b). However, due to the inability to achieve stable expression of BRCA1 fusions, the role of BRCA1 in GT regulation could not be conclusively determined (Figure 1c).

### Cell cycle synchronization approaches were not effective in tomato

During the S/G2 phases of the mitotic cell cycle, HR is more favorable (Heyer et al., 2010; Lieber, 2010). Therefore, it is reasonable to provide the GT tool to these favorable phases for enhancing its efficiency. Strong enhancements of CRISPR-Cas9-based gene editing via HDR were observed when the cell cycle was synchronized using chemicals to reversibly halt at the S or G2 phases (Lin et al., 2014). In mammals, biological approaches for cell cycle synchronization toward S/G2 phases were conducted successfully using a fusion of the N-terminal fragment of hGemini (1-110) (hGem) to the C-terminal of SpCas9 (Gutschner et al., 2016). Similarly, in plants, synchronization of protein expression in S-G2 phases was achieved by fusing the C-terminal fragment (3C) of the Cdt1a protein (CDT1a(3C)) to the N-terminal of marker proteins (Yin et al., 2014). When the S phase-specific promoter of the *HISTONE THREE RELATED2* (pHTR2) gene drove the expression of Cdt1a(3C), the synchronization of protein expression within the S-G2 phases was more robust (Yin et al., 2014). In this study, we aimed to improve GT efficiency by using biological approaches for S/G2 cell cycle synchronization with hGem, AtCDT1a(3C), and pHTR2 (Figure S1b). However, our data did not show significant alterations of the GT efficiency among the treatments (Figure S6a, b). Interestingly, the pHTR2 drove the enhancements of the indel mutation (Figure S6c, d) and 2-nt MH MMEJ (Figure S6e, f and Table S4) efficiencies at both the targeted sites. This suggests a shift of the repair pathway toward MMEJ but not HR. The enhancement of the indel mutation and MMEJ frequencies could be, in part, explained by the enhancement of ttLbCas12a protein accumulation under the control of pHTR2 (Figure S7).

### NKUDN enhanced the GT efficiency at the plant stage

An effective GT tool for plant breeding hinges on obtaining edited plants that carry GT alleles with a frequency that ensures their stable inheritance in the next generation. To accurately assess the performance of the KUDN addition to GT tools, we need to determine GT efficiency at the plant stage. We screened the plants obtained from cotyledon explants transformed with the GT constructs for the presence of GT alleles and GT efficiency using a series of methods (Figure S8). With the GT constructs for editing the *SlHKT1;2* locus, we obtained five GT0 events (F91.96, F92.31, F92.98, and F93.6 and F93.267) (Figure 3a and Figures S9-11) out of 535 analyzed transformants. Two of these events were from each construct with KUDN, which proved to be 1.43-1.62 times more efficient than the control construct without KUDN (as shown in Figure 3a, b). Additionally, NKUDN was found to be the best construct for GT at the *SlEPSPS1* locus, resulting in a 4.23% GT efficiency compared to only 0.43% in the control construct, which is a 9.84-fold enhancement (Figure 3a, b and Figures S12-13). One (F96.289) and three (F97.31, F97.32 and F97.35) *SlEPSPS1* events were obtained using the control and NKUDN-based GT tools, respectively. The data from plant stage analysis again indicate that NKUDN had a positive impact on GT in tomato.

**Figure 3.**
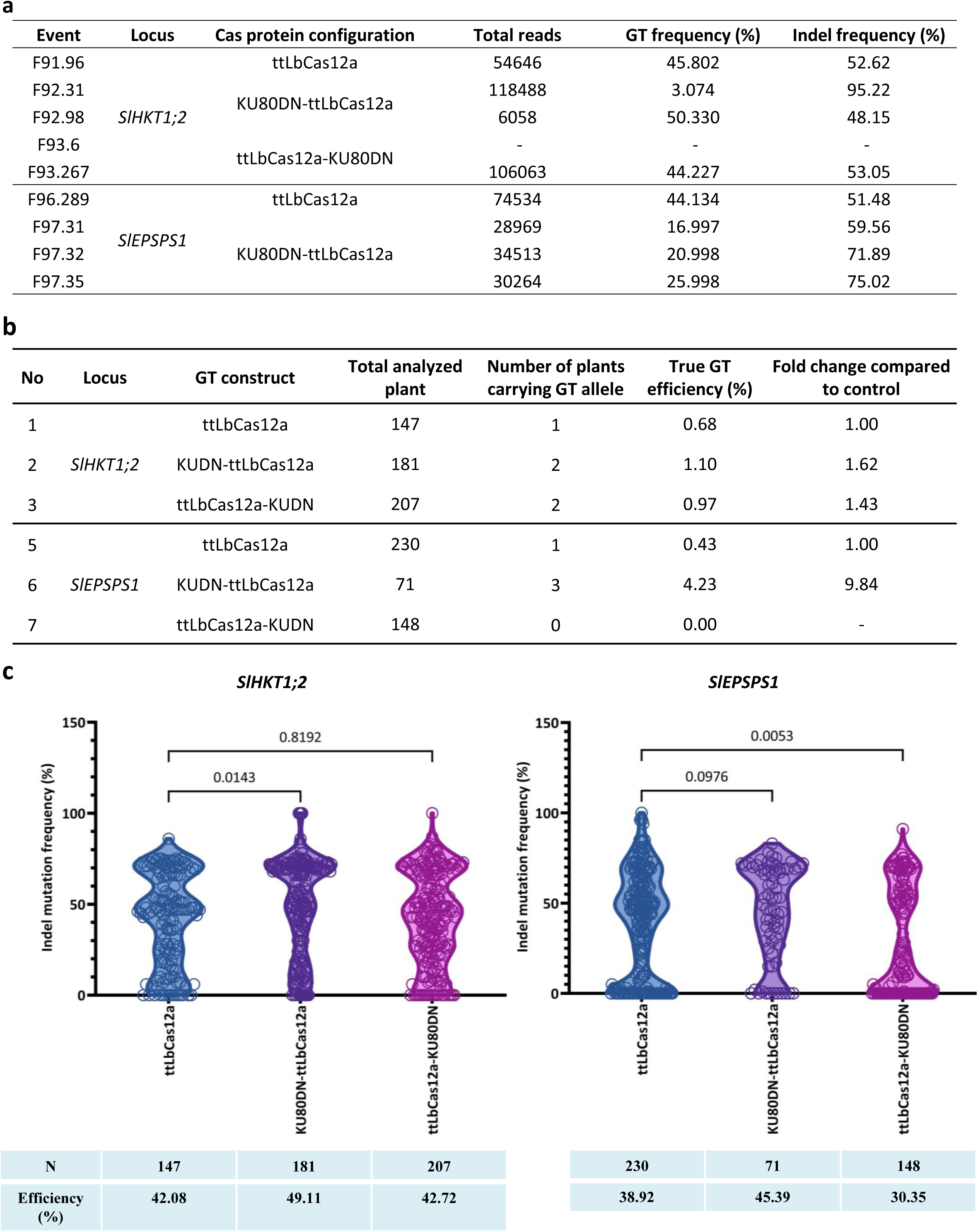
KUDN improved GT efficiency at the plant stage. (**a**) Validation of GT0 events by targeted deep sequencing. (**b**) Data showing the GT tools’ efficiency at the plant stage using KUDN and the control. (**c**) The indel mutation frequency and efficiency of the constructs without and with KUDN fusions. The indel mutation frequency of each transformant is shown as a data point in the violin plots. The statistical analysis was performed using GraphPad 9, and the p-values of the pair-wise comparisons are shown at the top. The bottom panel summarizes the number of analyzed transformants (N) and the average indel mutation frequency obtained by the constructs. The data points are shown on the plots.

After screening the transformants obtained from tissue culture of the explants transformed with the GT constructs, it was observed that the construct using NKUDN showed higher indel mutation efficiency and frequency of indel mutation within each plant, as depicted in Figure 3c. On the other hand, the CKUDN construct showed a comparable efficiency at the *SlHKT1;2* site while a decrease was observed at the *SlEPSPS1* site. This data is consistent with the results obtained through targeted NGS at the callus stage (Figure 1).

### KUDN-based GT tools show promise for broad gene targeting in tomato

We conducted an experiment to investigate whether the KUDN-based GT tools could be used to introduce specific DNA changes to other parts of the tomato genome. We tested the GT tools to insert a short DNA sequence to *SlCAB13* (Figure S14). At the plant stage, we screened twenty seven transformants and found one precisely edited CAB13 GT0 event (Figure S15). These results suggest that the KUDN-based GT tools could be used for other parts of the tomato genome, but their efficiency needs to be improved.

### The GT alleles were stably inherited in the next generations

The validity of the GT tools depends on the stable inheritance of the GT alleles from GT0 events to their progenies. To confirm this, we raised the next generations of the GT0 events and studied the inheritance of the GT alleles up to GT2. Our analysis showed that all the GT alleles carried by the GT0 events were stably inherited up to GT1 and GT2 generation (as shown in Figure 4 and Figure S16), thus confirming the reliability of the KUDN-based GT tools. However, the *SlEPSPS1* GT0 events were too weak to produce fruits and seeds for harvesting. This was likely due to the high levels of *SlEPSPS1* indel frequency (as seen in Figure S13) carried by the events, leading to insufficient activity of the *SlEPSPS1* enzyme that produces precursors of aromatic amino acids in the shikimate pathway (Maeda and Dudareva, 2012). The weakening of the shikimate pathway could result in reduced growth and fruiting of the edited tomato.

**Figure 4.**
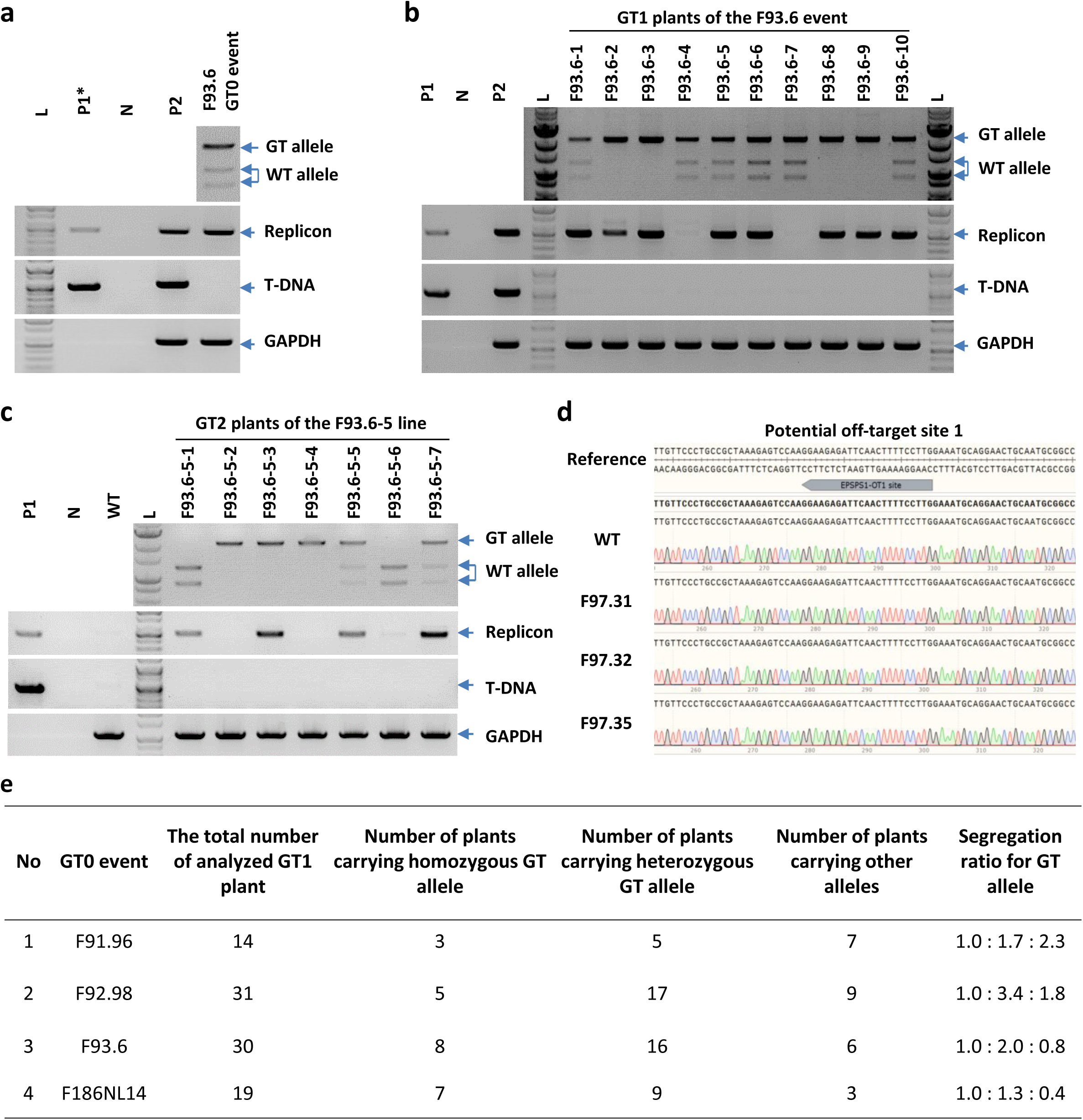
The inheritance of *SlHKT1;2* GT allele in GT1 generation. (**a-c**) CAPS assay revealed the GT allele and the presence of T-DNA and replicon in GT0 event F93.6 (**a**), its next-generation GT1, (**b**), and GT2 (**c**). (**d**) Chromatograms of the sequenced data show no editing traces at the potential off-target sites of *SlEPSPS1* gRNAs. (**e**) Table showing the segregation ratios of the GT alleles from four representative GT0 events. F91.96: *SlHKT1;2* GT0 event generated by the GT tool with ttLbCas12a; F92.98: *SlHKT1;2* GT0 event generated by the GT tool with KUDN-ttLbCas12a; F93.6: *SlHKT1;2* GT0 event generated by the GT tool with KUDN-ttLbCas12a; F97.31, F97.32, and F97.35: EPSPS1 GT0 events generated by the GT tool with KUDN-ttLbCas12a; F168NL14: *SlCAB13* GT0 event generated by the GT tool with ttLbCas12a-KUDN.

### Obtaining T-DNA and replicon-free GT events was possible via genetic segregation

Obtaining genome-edited plants without containing transgenes is the first demand to be accepted by public legislation before being considered for further steps in the commercialization of the edited plants (Chen et al., 2019; Gao, 2018; Voytas and Gao, 2014; Vu et al., 2022a). Our GT system employed a geminiviral replicon for delivering and amplifying GT tools to the targeted sites, thereby enhancing GT efficiency (Vu et al., 2020). Vu and co-workers showed the possibility of obtaining T-DNA and replicon-free GT plants in tomato’s next generation of GT0 events (Vu et al., 2020). Here, we also assess the presence of T-DNA and replicon in the GT0 events and their progenies.

In the GT0 generation, all events carried either T-DNA or replicon (Figure S17a). However, we were able to obtain T-DNA and replicon-free plants of the *SlHKT1;2* GT0 event F93.6 in GT1 (Figure 4b) and GT2 (Figure 4c) generations. This could be explained by the absence of T-DNA in the GT0 generation of the F93.6 event, which reduced the replicon load in the next generations of the plants. In the *SlHKT1;2* GT0 F91.96 event, we noticed that the first generation displayed a replicon band with a thin band of T-DNA (as shown in Figure S17a). In the subsequent generation, all the heterozygous and homozygous GT1 plants of the F91.96 event exhibited a very thin band of T-DNA. Two plants specifically, F91.96-6 and F91.96-10, did not show any replicon or T-DNA band (as indicated in Figure S18a). This suggests that homozygous plants that are free of T-DNA and replicon can be obtained in the GT2 plants, even if the first generation plant contained T-DNA and replicon.

For the *SlHKT1;2* GT0 F92.98 event, we observed thick bands of T-DNA and replicon (as depicted in Figure S17a). Thus, in its GT1 generation, almost all plants displayed replicon and T-DNA bands, although some had only thin bands of replicon and no T-DNA, such as F92.98-1 and F92.98-4 (as shown in Figure S18b). These plants were raised to evaluate T-DNA and replicon in the next generation. It may be more difficult to obtain T-DNA-and replicon-free plants from GT events that carried both T-DNA and replicon in the first generation, such as the CAB13 event F168NL14, since its GT1 plants mostly had replicons (Figure S17b). We found some plants like F168NL14-8 and 9 without T-DNA, while the other only showed weak T-DNA bands. It may also allow the obtaining of T-DNA and replicon-free plants in their GT2 generation, as obtained in the case of the F93.6 event. Regarding GT2 homozygous plants, we obtained one plant, F92.98-9-17, which was free of replicon and with a very thin T-DNA band (Figure S19a). Therefore, in the next generation, we could obtain homozygous plants that are free of both T-DNA and replicon.

Similarly, for the *SlCAB13* event, we observed that the GT1 line F168NL-14-9 had a very thin band of replicon and T-DNA (Figure S17b). However, in the GT2 plants from this line, we obtained all plants that were free of T-DNA and only some plants showed a band of replicon (Figure S19b). Consequently, we have successfully obtained many homozygous plants that are free of both T-DNA and replicon in GT2 plants such as F168NL14-9-(2,4,5,6,7,8,9,11,12,13,15,16,17) (Figure S19b).

### The KUDN-based GT tools maintained ttLbCas12a specificity without any detected off-target traces

The CRISPR-LbCas12a system uses a 20-to 24-nt gRNA to guide the process. However, this gRNA may bind to unwanted sites adjacent to a T-rich PAM, which can activate the cleavage by LbCas12a and result in off-target indel mutations (Murugan et al., 2020). Cas12a proteins are highly specific due to their sensitivity to mismatches within the gRNA binding sequence (Zetsche et al., 2015). In plants, LbCas12a and its orthologs have shown high accuracy in editing (Vu et al., 2020; Xu et al., 2017; Zhang et al., 2021). The ttLbCas12a used in this study was also highly specific in tomato (Vu et al., 2021a; Vu et al., 2020).

To test if the KUDN-based GT tool can alter the specificity of ttLbCas12a, we analyzed and identified potential off-target sites of the used gRNAs by Cas-offinder (Bae et al., 2014) with less than four mismatches. We found that only some gRNAs that targeted *SlEPSPS1* and *SlCAB13* loci contain several potential off-target sites (Figure S20a). We then amplified and sequenced the sequences flanking the identified potential off-target sites in the GT events (Figure S20b). However, no off-target trace was found at all the potential off-targets (Figure 4d and Figure S20c). This indicates that the KUDN-based GT tools are highly specific, and adding KUDN did not alter the specificity of ttLbCas12a.

## DISCUSSION

Among the CRISPR-Cas-based approaches, CRISPR-Cas-based GT is the only technique that allows for precise gene/allele replacement at large scales or complexed edits, which has not yet been achievable with other precise methods like prime editing (Chen and Liu, 2023; Vu et al., 2019). We have developed a geminiviral replicon and ttLbCas12a-based GT system that achieved efficient gene insertion using a double selection method (Vu et al., 2020). However, the GT efficiency remains low without using a target-associated selection marker (Vu et al., 2021a; Vu et al., 2020) and requires improvement for practical applications.

The low GT efficiency is widely conservative from animals to plants and has a common reason: the cNHEJ is dominant over its competitive HDR pathway for repairing DSBs. Multiple approaches have been successfully employed for enhancing HR in animals that could be borrowed to enhance GT in plants. In this work, we sought to study the approaches for GT improvement by (1) stimulation of DSB end resection with end processing enzymes, (2) biasing the DSB repair pathway choice toward HR, and (3) synchronizing the expression of GT tools at the HR-favorable S-G2 phases of the cell cycle. Despite we observed no enhancement of the GT performance with the first and the third approaches, directly disturbing the initiation of cNHEJ using KUDN led to significant enhancement of GT efficiency at the plant stage. We reasoned that the interference of cNHEJ by KUDN increases the activation of the HDR pathway, thereby enhancing GT efficiency.

The resection of DSBs is necessary for strand annealing and repair by HDR mechanisms. Addition of DSB end processing enzymes has been shown to facilitate HR-mediated repair in mammals (Charpentier et al., 2018; Hackley, 2021; Park et al., 2021; Zhang et al., 2020). However, in our experiments, adding nucleases to targeted sites only changed the profile of the edited products (Figure S2). One reason for this could be the difficulty in accumulating fused proteins (Figure S3). In the case of T5exo-ttLbCas12a fusion, the protein accumulation and cleaving profile was better than ttLbCas12a alone, but GT efficiency was only slightly improved. Interestingly, ttLbCas12a-CtIP fusion led to a reduction in cleavage due to low protein accumulation, but GT efficiency was slightly improved at both targeted sites (Figure S2a, b), indicating that CtIP activity might alter DSB repair towards HDR. Another possible reason for the observed data is the difference in expression systems between cell line-based experiments in animals and *Agrobacterium*-mediated delivery of GT tools. The former could deliver high doses of GT complexes and donor DNAs simultaneously, while the latter may not be able to mimic the same loads. If this is the case, there could be general difficulties in repeating the observation of GT data reported in animals.

The analysis of the targeted NGS data during the callus stage showed that the NKUDN enhanced GT efficiency from 1.71-3.55 folds the tested sites (Figure 1). The enhancing effects of NKUDN were also validated in the plants at both loci, where NKUDN had a 1.62-fold (*SlHKT1;2*) and 9.84-fold (*SlEPSPS1*) enhancement compared to the control (Figure 3), correlated with the enhancement observed in the *in vitro* samples (Figure 1b). This data suggests that NKUDN increased the number of cells carrying the GT alleles, leading to a higher chance of obtaining regenerated plants with the GT alleles. It is worth noting that the GT efficiency obtained during the plant stage is more reliable than that of the callus stage, thereby validating the positive impacts of NKUDN for GT in tomato. This observation possibly reflects a reduction in cNHEJ repair of DSBs due to the malfunctioning of KU complexes in tomato cells that is consistent with the pioneering work in CHO cells (Marangoni et al., 2000) or in *Arabidopsis* (Tamura et al., 2002; West et al., 2002). Moreover, the GT enhancement is similar to what was observed in plants with a *ku70*-knockout background (Nishizawa-Yokoi et al., 2012; Qi et al., 2013).

The efficiency of GT can be significantly increased by introducing a DSB at the targeted site (Puchta et al., 1993). The CRISPR-Cas system utilizes a gRNA-Cas protein complex to introduce DSBs to targeted sites, thereby enhancing GT efficiency (Vu et al., 2019; Wang et al., 2017; Xu et al., 2014). Therefore, the formation of DSBs correlated with the expression levels of the Cas protein, which may affect GT frequency. However, since we observed no significant improvement in protein accumulation between NKUDN and control (Figure 1d), the enhancement of GT efficiency could be due to the antagonist role of NKUDN toward the KU complex that redirected the repair choice from the cNHEJ route to MMEJ and HR.

When the process of cNHEJ malfunctions, it can redirect the repair of double-strand breaks to the HDR pathway, which includes MMEJ, SSA, and HR (Nishizawa-Yokoi et al., 2012; Qi et al., 2013; Vu et al., 2021b). These subpathways require different levels of end resections at varying speeds of repair kinetics (Symington and Gautier, 2011; Taleei and Nikjoo, 2013; Vu et al., 2022b). The deletion of short DNA sequences at targeted sites through MMEJ was observed to shift in the editing frequencies (Figure 2). The use of KUDN constructs led to an increase in MMEJ frequency at the tested loci, as previously recorded by Qi and colleagues in a *ku70*-deficient background(Qi et al., 2013). The impact on MMEJ was more pronounced with NKUDN, which was correlated with the enhancement of GT efficiency since the common end resection step initiated both subpathways. The MMEJ traces extracted from targeted NGS data were intrachromosomal repaired products, providing more direct evidence of the KUDN-induced suppression of cNHEJ and subsequent redirection of the DSB repair to the HDR pathway. As a result, the enhancement of MMEJ and other alternative DSB repair pathways that are more error-prone led to an overall increase in the indel mutation efficiency of the NKUDN-based GT tools at both the callus and plant stages (as shown in Figure 1b and Figure 3c, respectively). The enhancement of MMEJ frequency by the KUDN may be an interesting feature to improve the chromosome engineering approaches such as inversion (Schmidt et al., 2019) or translocation (Beying et al., 2020).

In this study, the cell cycle synchronization approaches were not as effective as reported in animals, possibly due to the differences in delivery of GT tools and the differences in cell types and kinetics of the repair mechanism. Interestingly, the pHTR2 drove enhanced expression of ttLbCas12a resulting in a higher indel mutation and 2-nt MH MMEJ, but not the GT efficiency. It is possible that the pHTR2 stimulated the ttLbCas12a accumulation at the early S phase when the sister chromatids are not available and thus the HR mechanism is not at the most favorable conditions. Moreover, the MMEJ pathway is active during G1 until early S phase (Taleei and Nikjoo, 2013) that might involve efficiently in repairing the DSBs formed by ttLbCas12a at the early S phase leading to the pHTR2-mediated enhancement of the indel mutation and MMEJ efficiency (Figure S6c, f).

During the plant stage, we obtained several GT0 events from different constructs (Figure 3 and Figures S9-13). The NKUDN-based construct returned a more significant number of GT0 events and also exhibited higher indel mutation efficiency (Figure 3). This was correlated with the target NGS data (Figure 1). We used these tools in other loci which resulted in a GT0 event from the *SlCAB13* locus (Figure S15). Notably, the GT alleles were inherited in the next generations (Figure 4 and Figure S16), indicating that the GT alleles were stably fixed into the tomato’s genome. This observation validates our KUDN-based GT tools and is consistent with other reports of GT in plants (Cermak et al., 2015; Dahan-Meir et al., 2018; Merker et al., 2020a; Merker et al., 2020b; Schindele et al., 2023; Wolter and Puchta, 2019). More importantly, the GT tools did not induce off-targeting activities in other genome sites (Figure S20). The T-rich PAM LbCas12a is more specific compared to SpCas9 due to its more extended PAM sequence and hyper-sensitivity toward mismatches within the binding sequence of gRNAs (Kim et al., 2016; Vu et al., 2020; Xu et al., 2017; Zetsche et al., 2015).

One of the most important considerations when assessing a gene editing technique is whether the editing tool can be removed from the edited plant or integrated into the plant’s genome in a stable way. This is the first criterion for determining whether the edited product is subject to strict or relaxed regulation (Buchholzer and Frommer, 2023; Metje-Sprink et al., 2018). In this study, we found that all GT0 events contained either T-DNA or replicon of both cargoes (Figure 4a and Figure S17a), but we were able to isolate T-DNA-free and replicon-free GT1 (Figure 4b) and GT2 (Figure 4c and Figure S19) lines of F92.98, F93.6 and F168NL14 GT0 events by genetically segregating the integrated T-DNAs and replicons (Figure 4A and Figure S17a). This could be due to the fact that the F92.98-1 GT1 plant, F93.6 GT0 event, and F168NL14-9 plants contained replicons but not T-DNA (Figures S17-18). The replicons existed in a circularized form that did not stably integrate into the genome and could potentially be segregated or diluted out during sexual reproduction (Figure 4b, c and Figure S19). It is well-known that the replication of viral genomes is largely suppressed in germline cells by systemic RNAi-based gene silencing (Fortes et al., 2023; Lei et al., 2021; Schwach et al., 2005). It is important to note that the efficient transmission of geminiviral genomes through seeds may be limited to specific strains and has a size constraint (Kil et al., 2016; Rojas et al., 2018). In our study, the reduced seed transmission rate was possibly due to the large size of the GT tool-carrying circularized replicons engineered from a BeYDV (Vu et al., 2020). In addition, the KUDN might have some negative effects on the integration of T-DNA to the genome of regenerated plants (Nishizawa-Yokoi et al., 2012). Therefore, we can expect to obtain T-DNA-free and replicon-free GT0 or GT1 lines that did not have T-DNA integration, as effectively shown with the G93.6-5 GT1 line and the GT2 generation of F92.98-1 and F168NL14-9 GT1 plants.

Taken together, our data support the hypothesis that addition of KUDN interferes with cNHEJ, shifting DSB repair to HDR subpathways and ultimately enhancing MMEJ and HR-mediated GT efficiency.

## CONCLUSIONS

Precision plant genome editing is crucial for fast-breeding crops to tackle current agriculture challenges such as climate changes and arable land shortages. CRISPR-Cas-based GT is an extremely precise genome editing technique that enables kilobase-scale gene/allele replacements, which can save time and labor in plant breeding. GT tools are also useful for precise gene pyramiding and *de novo* domestication of genes/alleles from wild relatives. Therefore, enhancing GT efficiency is a priority mission for crop breeding.

The plant GT tool has been significantly improved by adding a DSB generator, the CRISPR-Cas molecular scissors such as ttLbCas12a and SpCas9, and geminiviral replicon, the homologous DNA donor amplifier. In this work, we successfully added KUDN as a novel component to the CRISPR-LbCas12a-based GT tool, which dramatically improved GT efficiency at the plant stage (Figure S21). The novel NKUDN-GT tools do not generate off-target editing, and T-DNA-free and replicon-free lines could be reliably obtained in the progenies of GT0 events. Our novel GT tool can be employed in other crops and significantly contribute to the precision plant breeding field soon.

## EXPERIMENTAL PROCEDURES

### System design and plasmid cloning

Initially, the *tomato HKT1;2*, and *EPSPS1* loci were selected as targets for conducting HR-based allele replacement experiments since they were studied in our lab (Vu et al., 2021a; Vu et al., 2020). Subsequently, *SlCAB13* was used to show the applicability of the improved KUDN-carrying GT tool. *SlCAB13* is a gene encoding for a light-harvesting chlorophyll a/b binding protein (type III, homolog 13) that plays essential roles in photosynthesis. Missing AAATTGTGA proximal to the start codon of *SlCAB13* in domesticated tomato led to sensitivity to continuous lighting conditions (Velez-Ramirez et al., 2014). We designed the GT construct to restore the bases to the corresponding location of the *SlCAB13* gene (AAATTGTGA insertion) that may help tomato tolerate continuous lighting conditions, thereby increasing tomato yield. The homologous donor for targeting *SlCAB13* and the gRNA expression cassette were designed and cloned accordingly, as shown in Figure S14 and Supplemental sequence file.

A geminiviral replicon system was employed as the vector for the delivery of guide RNA and CRISPR-Cas expression cassettes, as well as GT donor templates (Figure 1a and Figure S1) of all the constructs. A dual gRNA construct was employed for each of the targeted sites, which were designed in tandem repeats of LbCas12a scaffolds and gRNA sequences (Figure S1a). The loci and gRNA sequences are listed in Table S5. The plasmids containing fusions (hExo1a, T5exo, hHE, CtIP, Ku80DN, BRCA1, AtCDT1a(3C), and Gem), promoter pHTR2, expression cassettes and the binary vectors (Figure S1) were cloned using Golden gate assembly method (Engler et al., 2014; Weber et al., 2011). The DNA sequences are listed in Supplemental sequence file. Vu and coworkers designed the homologous donors of the *SlHKT1;2* and *SlEPSPS1* earlier (Vu et al., 2021a).

The long CaMV35S long promoter (Addgene #50267) and CaMV 35S terminator (Addgene #50337) were used in all the Cas protein expression cassettes. An additional copy of the *AtTRP1* intron 1 was inserted into the coding sequence of the ttLbCas12a variant, a modification previously tested in GT experiments by Vu and colleagues (Vu et al., 2021a; Vu et al., 2020). The crRNA and sgRNA expression cassettes were transcribed using the core sequence of the AtU6 promoter (Nekrasov et al., 2013) and terminated by an oligo dT (7xT).

### *Agrobacterium*-mediated tomato transformation

For our study, we used a local tomato cultivar called Hongkwang and performed *Agrobacterium*-mediated tomato transformation following the established protocol of our laboratory (Vu et al., 2020). First, *A. tumefaciens* GV3101::pMP90 cells were cultured overnight in primary culture using LB medium with appropriate antibiotics in a shaking incubator at 30°C. The agrobacterium cells were then harvested from the culture by centrifugation and suspended in a liquid ABM-MS medium (pH 5.2) with AS (200 µM). The OD600nm of the suspension was adjusted to 0.8. Cotyledon fragments were cut from 7-day-old seedlings and pre-cultured on the PREMC medium (containing MS salts, B5 vitamins, 2 mg/L zeatin trans-isomer, 0.5 mg/L IAA, 1 mM putrescine, 0.5 mg/L MES, 30 g/L maltose, and 7.5 g/L agar) one day before the transformation. The transformation process involved mixing tomato cotyledon fragments with the bacterium suspension and keeping them at room temperature for 25 minutes. The explants were then transferred to the cocultivation medium, which contained all the elements from the ABM-MS medium and AS (200 µM) at pH 5.8. These cocultivation plates were placed in darkness at 25°C for two days, and the explants were moved to a non-selection medium (NSELN) for five days before being subcultured into the selection medium (SEL5).

The NSELN and SEL5 media contained all the components of the pre-culture medium, supplemented with 250 mg/L timentin and 60 mg/L kanamycin. NSELN also contained 2µM NU7441. The explants were subsequently sub-cultured on SEL5R (SEL5 with 1 mg/L zeatin trans-isomer), SEL4CA (SEL5 with 0.5 mg/L zeatin trans-isomer, 0.05 mf/L IAA, 0.1 g/L ascorbic acid, and 5 mg/L AgNO3), and SEL4C (SEL4CA without AgNO3) every 15 days to improve regeneration efficiency. Once the shoots grew to a length of around 2.0 cm, they were subcultured to the RIM medium for rooting. The RIM medium was similar to SEL4C, but without zeatin trans-isomer and IAA, and contained 0.1 mg/L NAA and 0.3 mg/L IBA. The shoots with roots were acclimated in a greenhouse under 16 hours of light and 8 hours of darkness at a temperature of 26 ± 2°C.

### Plant genomic isolation by CTAB method

To extract the genomic DNA (gDNA) from plant leaves or cotyledon explants/callus, we used the CTAB method with some minor modifications based on Vu and coworkers (Vu et al., 2021a). We started by grinding approximately 200 mg of leaf sample in liquid nitrogen and then adding 300 µL of Solution I, which contained 1M NaCl, 2% Sarkosyl, and 5 µL of RNase at 10 mg/mL/tube. The mixture was then incubated at 37°C for 30 minutes and centrifuged at 13,000 rpm at 4°C for 10 minutes. We then incubated 250 µL of the supernatant with 500 µL of extraction buffer (containing 100 mM Tris-Cl, 20 mM EDTA, 1.4 M NaCl, and 2% CTAB) at 60°C for 35 minutes. After that, we added 750 µL of Chloroform:Isoamylalcohol (24:1) and centrifuged the mixture at 13,000 rpm at 4°C for 15 minutes. We added 20 µL of 3M CH3COONa (pH 5.2) and 360 µL of isopropanol to the 600 µL supernatant liquid, followed by centrifugation at 13,000 rpm at 4°C for 5 minutes. The genomic DNA (pellet) was washed with 80% ethanol and centrifuged again at 13,000 rpm at 4°C for 5 minutes. Finally, we incubated the DNA at 37°C for 30 minutes to remove any residual ethanol and dissolved the gDNA pellet in 50 µL of DNA Elution buffer (EB). We then used a NanoDrop 1000 UV/Vis Spectrophotometer (NanoDrop Technologies Inc., Wilmington, DE, USA) to determine the quality and concentration of the DNA.

### Targeted deep sequencing

The method used for targeted NGS analysis was previously described by Vu and coworkers (Vu et al., 2021a) and was slightly modified for this study. PCR was used to amplify the targeted sites in the gDNA isolated from the cotyledon/callus samples, using the first PCR primer pair (Table S6) which was located outside the homologous sequences and flanking the editing sites. The second and third PCR reactions were carried out according to the guidelines provided by the miniseq sequencing service provider (KAIST Bio-Core Center, Daejeon, Korea). The raw data files were pre-processed by the NGS service provider and were subsequently analyzed using the RGEN Cas-analyzer tool (Park et al., 2017) and CRISPResso2 (Clement et al., 2019). The targeted deep sequencing of the first batch of experiment (end resection facilitation) was conducted with 10-dpt samples as reported earlier (Vu et al., 2021a). However, since the GT efficiency was low and highly variable, we decided to use 21-dpt samples for the analysis afterward.

At the callus stage, GT efficiency is calculated as follows: GT(%) = 100 x (number of NGS reads containing GT alleles / total NGS reads).

### Validation of GT events by molecular analyses

The CTAB method was used to extract genomic DNAs from tomato. PCR was used to verify the gene targeting (GT) sites and the donor junctions, by using the 1st primer pairs that covered the respective sites (Table S6). High-fidelity Taq DNA polymerase (Phusion Taq, Thermo Fisher Scientific, Massachusetts, USA) was employed in this process, followed by Sanger sequencing (Solgent, Daejeon, Korea) for analysis. To differentiate GT-edited events due to changes in the cutting site of enzymes during donor design and cloning, the cleaved amplified polymorphic sequence (CAPS) method was used. BpiI restriction enzyme was used to digest PCR products. The binding site of BpiI was modified in the GT alleles. BpiI did not digest potential GT products, and the PCR products containing the undigested band were sent for Sanger sequencing. Furthermore, the ICE-Synthego program was used to decompose the Sanger sequencing results, which helped in identifying indel mutations and GT efficiency within the plant. We then assessed the potential GT events that showed GT allele frequency > 10% in ICE Synthego data by targeted NGS to validate whether they were true GT events. Their PCR products were cloned into pJET1.2 blunt plasmids (Thermo Fisher Scientific, Massachusetts, USA) and sequenced.

GT efficiency at the plant stage was calculated as GT(%) = 100*(Total plants carrying GT alleles/Total analyzed plants).

In order to determine whether GT events contained T-DNA and replicon, PCR reactions were performed using the primers listed in Table S6. The protocol provided by the manufacturer of Diastar Taq DNA polymerase (Solgent, Daejeon, Korea) was followed, and the PCR reactions were run for 30 cycles. The resulting PCR products were then separated on a 0.8% agarose gel. The presence or absence of T-DNA or replicon was determined based on the presence or absence of DNA bands on the agarose gels.

### Western blot analysis

Tobacco leaves that were 35 days old were infiltrated with Agrobacterium clones containing the GT constructs. After 3 days, an equal amount of infiltrated tobacco leaf samples were collected and kept in liquid nitrogen for grinding. Total proteins were extracted using a previously reported protocol (Kumar et al., 2017). The extracted proteins (around 40 µg) were loaded onto 5% SDS-PAGE gels with a PM2610 protein ladder (SMOBIO, Hsinchu, TW) and electrophoresis was run at a fixed 100 V for 2 hours. The proteins were then transferred from the gel to a PVDF membrane (Immonilon-P transfer membrane, Merck, Darmstadt, Germany) using the wet transfer method, which was conducted for 2 hours with a transfer buffer containing Tris-base, glycine and 20% methanol. The membrane was blocked using a buffer containing 5% low-fat milk powder in 1x TBS-T (Tris, NaCl, Tween) under gentle shaking conditions (50 rpm) for 2 hours. The membrane was then incubated overnight in a buffer containing 5% low-fat milk powder in 1x TBS-T and HA antibody (1:5000) at 4°C. The β-actin-specific antibody (1:20000) was used as an internal control. The next day, the membrane was washed five times with 1x TBS-T and the second antibody (anti-rabbit, diluted at 1:10000 ratio in 1x TBS-T) was added to the membrane. The membrane was incubated at 4°C overnight. The following morning, the membrane was washed twice with the 1x TBS-T buffer and then incubated with the Clarity Western ECL Substrate solution (Bio-Rad, California, USA). Signal detection of the membranes was performed by the Chemi-Doc Imaging system (Bio-Rad, California, USA) after a 5-minute incubation. The images were taken every 5 seconds for 30 seconds and the protein expression of the GT constructs was quantified using ImageJ software (NIH, Maryland, USA).

### Off-target analysis

In order to identify any potential off-target locations in the tomato genome, we used the sequence of each gRNA as a query sequence in http://www.rgenome.net/cas-offinder. For this purpose, we selected the PAM type of LbCpf1 (5’TTTN3’) and the *Solanum lycopersicum* (SL2.4)-Tomato genome. We then designed specific primers for off-target sites using the NCBI PrimerBlast and used them for PCR amplification and Sanger sequencing (as outlined in Table S7) to verify the results.

### Assessment of the inheritance of GT alleles in the next generation of GT0 events

After the successful validation of GT0 events, they underwent self-pollination, and the resulting seeds were collected and stored for further investigations. These seeds were then planted in soil and the genomic DNAs of the offspring plants were individually extracted for examination through Sanger sequencing. The sequencing chromatogram files obtained were subsequently analyzed with the ICE Synthego tool, which provided useful insights into the GT and indels frequencies in each plant. Moreover, to assess the presence of replicon and T-DNA constructs, PCRs were performed using the primers specified in Table S6.

### Statistical analysis

All comparison experiments were conducted with at least three replicates, and the data was collected by counting purple spots, performing targeted deep sequencing, and screening plant events. Some of the experiments using targeted deep sequencing were conducted in two replicates. The editing data, statistical analysis, and plots were further processed using MS Excel and GraphPad Prism 9 programs and explained in detail in the legends of Figures and Tables. Pairwise comparison data were tested with Student’s t-test with unequal variance and two-tailed parameters. Fisher’s LSD test was applied for multiple comparisons using similar parameters. A difference was considered significant when the statistical tests returned a p-value of less than 0.05.

## ACKNOWLEDGMENTS

This work was supported by the National Research Foundation of Korea (Program 2020M3A9I4038352, 2020R1A6A1A03044344, 2021R1A5A8029490, 2022R1A2C3010331) and the Program for New Plant Breeding Techniques (NBT, Grant PJ01686702), Rural Development Administration (RDA), Korea.

## AUTHOR CONTRIBUTIONS

T.V.V. and J.Y.K. conceived and designed the research. T.V.V., N.T.N., J.K., M.H.V., Y.J.S., M.T.T., and Y.W.S. conducted experiments. T.V.V., N.T.N. and J.Y.K. analyzed data. T.V.V., N.T.N., and J.Y.K. wrote the manuscript. T.V.V. and J.Y.K. finalized the manuscript. All authors read and approved the manuscript.

