## Supplemental Materials for "Redirecting DNA repair for efficient CRISPR-Cas-based gene targeting in tomato"

**Running title: Enhanced CRISPR-Cas-based GT in tomato.**

#### Supplemental Figures

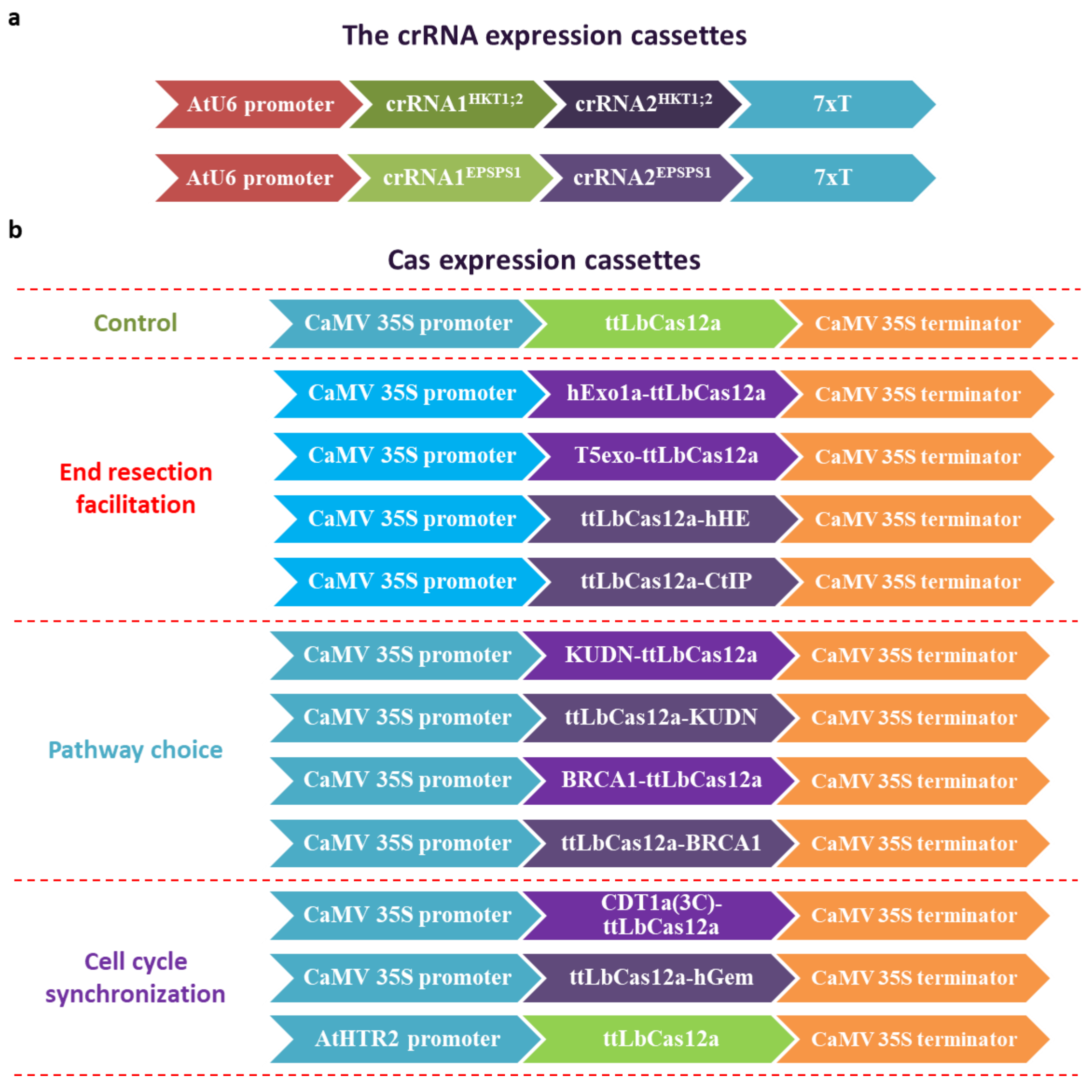


**Figure S1. The expression cassettes of gRNAs and Cas protein used in the study. a** Two crRNAs (LbCas12a scaffold and gRNA) were cloned in tandem repeats under the driving of the AtU6 promoter and terminated by seven T (7xT) for cleaving each targeted site. **b** The Cas protein expression cassettes were designed for the Control, End resection facilitation, Pathway choice, and Cell cycle synchronization using either the CaMV 35S promoter with or without the TRP1 intron 1 and the CaMV 35S terminator.

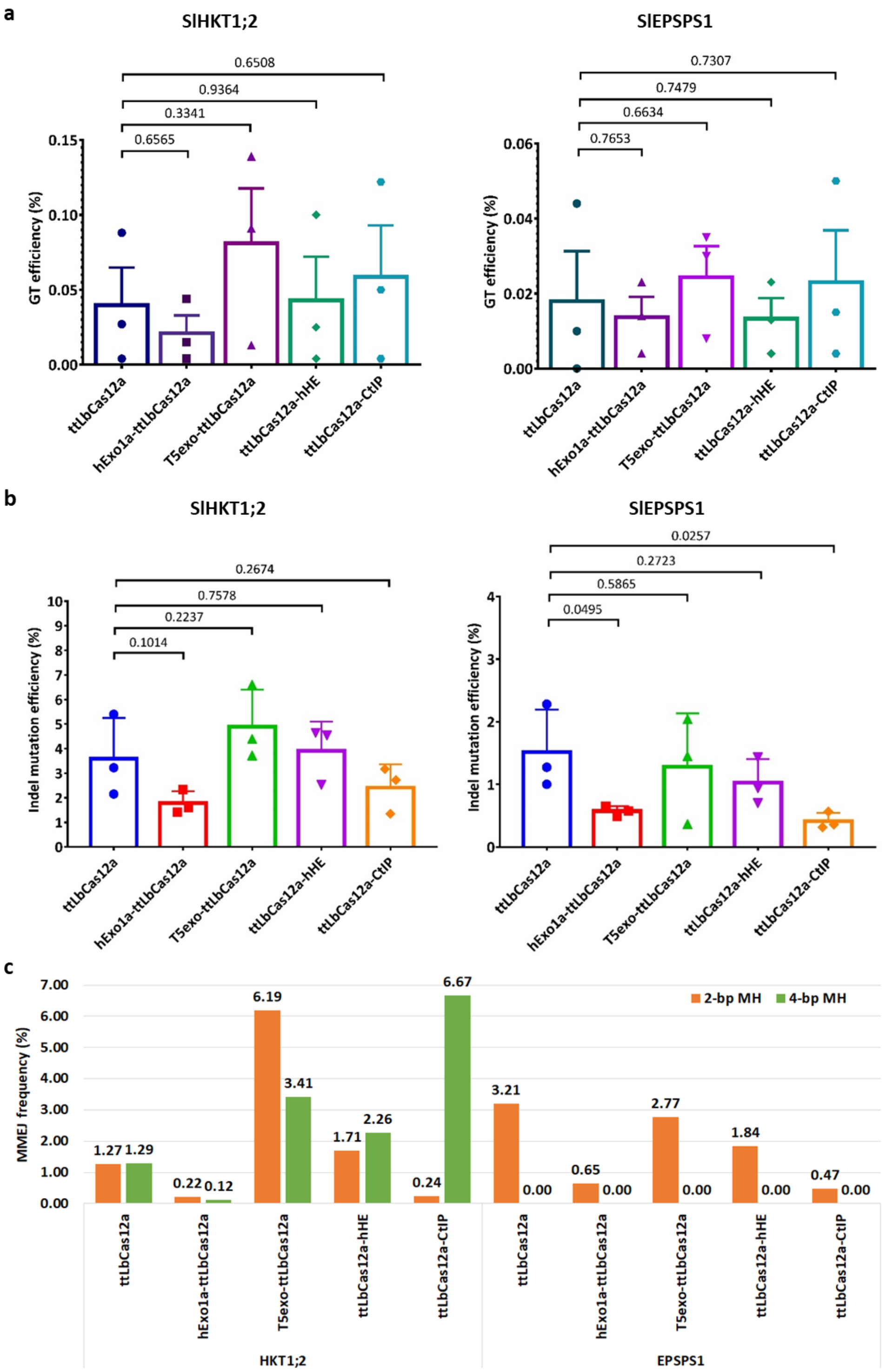


**Figure S2. The impact of end resection facilitation by nucleases on the editing efficiency in tomato. a-c.** The impacts of the nucleases on GT (**a**), indel mutation (**b**), and 2-nt and 4-nt MH MMEJ (**c**) efficiency at the callus stage. The efficiencies were assessed by targeted NGS using 21-dpt cotyledon/callus samples. The data points are shown as the dots on the plots.

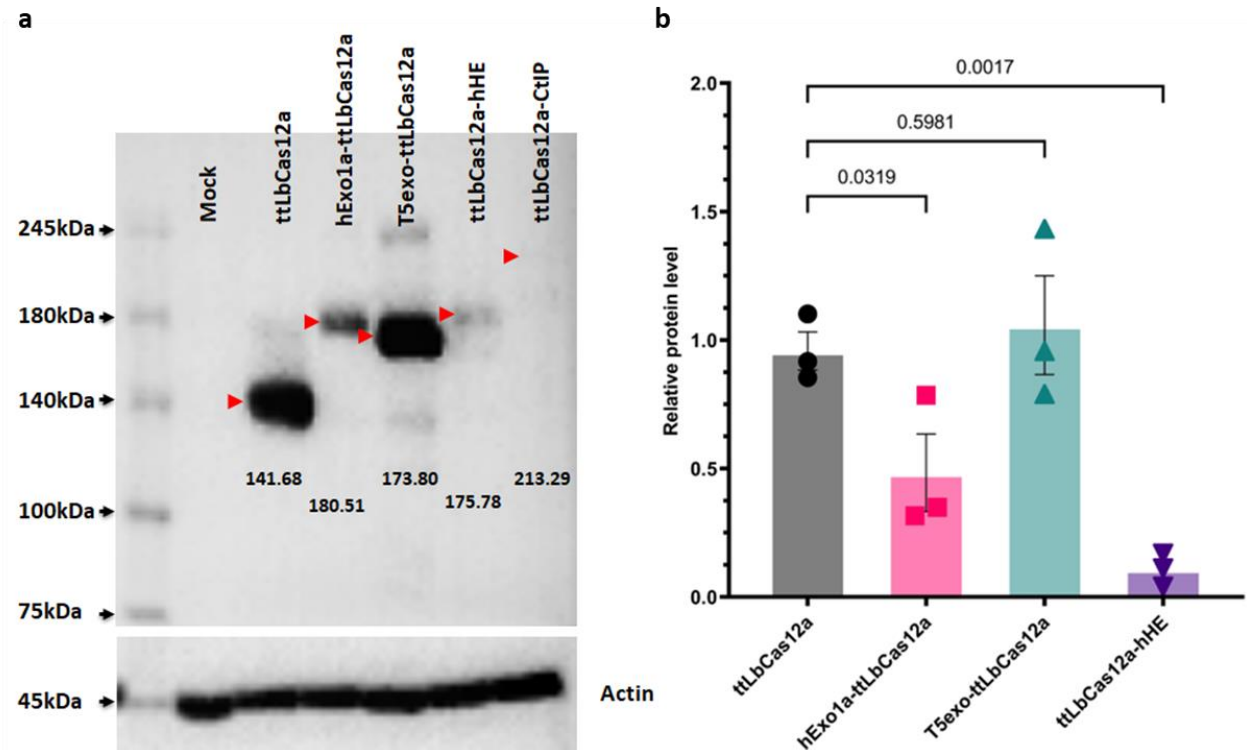


**Figure S3. The protein expression levels of ttLbCas12a and its fusions with the end resection nucleases.** **a** Western blot membranes showing bands of the Cas proteins with expected sizes and expression cassettes. Actin levels were used as loading controls. The red arrows denote the position of expected bands. **b** Relative protein levels of the Cas protein. P-values of the t-test for pair-wise comparison between the relative protein levels of ttLbCas12a and the fusions are indicated on the top of the bars. The expression cassettes and Cas configurations are at the bottom of the bars. The data points are shown as the dots on the plots.

```
Range 1: 4 to 704 GenPept   Graphics                                    ▼ Next Match ▲ Pre

Score          Expect Method                        Identities      Positives       Gaps
206 bits(524)  8e-57  Compositional matrix adjust.  205/761(27%)    357/761(46%)    95/761(12%)

Query  6    NKAAVVLCMDVGFTMSNSIPGIESPFEQAKKVITMFVQRQVFAENKDEIALVLFGTDGTD  65
            NK A+VL +DVG +M + +P IE       KV ++ +Q+++     DE+  VLFGT  T
Sbjct  4    NKEALVLVIDVGPSMHSVLPEIE-------KVCSLLIQKKLVFSRYDEVGFVLFGTADTK  56

Query  66   NPLS---GGDQYQNITVHRHLMLPDFDLLEDIESKIQPGSQQADFLDALIVSMDVIQHET  122
            N L    GG  Y+++TV R++ + D DL++ ++ K+  GS   DFLDA++V  D++
Sbjct  57   NELKEEIGG--YEHVTVLRNIKVVDEDLVDALQ-KLPRGSVPGDFLDAIVVGTDMLIKFG  113

Query  123  IGKKFEKRHIEIFTDLSSRFSKSQLDIIIHSLKKCDISLQFFLPFSLGKEDGSGDRGDGP  182
               K  KR + + T+  SR                     PF   KED
Sbjct  114  RTNK-AKRRLCLITNAVSRIKD---------------------PFEGTKEDQVNTIAT--  149

Query  183  FRLGGHGPSFPLKGITEQQKEGLEIVKMVMISLEGEDGLDEIYSFSESLRKLCV------  236
             ++   G    +  +  + K+  E  + +M   E  D L  ++S   S + + V
Sbjct  150  -QMTAQG--IKMDCVIVRMKQDRETNRSIM---EENDFLMSVFSNKSSSKVVYVESPTSL  203

Query  237  FKKIERHSIH----WPCRLTIGSNLSIRIAAYKSILQERVK--KTWTVVDAKTLK--KED  288
              +   +I      +    L I + L I++  YK    +E+     K ++       T K    D
Sbjct  204  LGALRTRNISPVTIYRGDLEISAQLKIKVWVYKKTSEEKFPTLKKYSERTPTTDKFGAHD  263

Query  289  IQKETVYCLNDDDETEVLKEDIIQGFRYGSDIVPFSKVDEEQMKYKSEGKCFSVLGFCKS  348
            I+ E     + +D    V  E  I+GF+YG  +VP S  + E +K+K E K  +LGF  S
Sbjct  264  IKVEYENKIIEDPNKVVPPEQRIKGFQYGPQVVPISSAELEAVKFKPE-KSVKLLGFTDS  322

Query  349  SQVQRRFFMGNQVLKVFAAR-DDEAAAVALSSLIHALDDLDMVAIVRYAYDK-RANPQVG  406
            S + R +++ +  + +F A   ++ A +ALS+L  A+ +++ VAIVR  + + + N  VG
Sbjct  323  SNIMRHYYLKD--VNIFIAEPGNKNAILALSALARAMKEMNKVAIVRCVWRQGQGNVVVG  380

Query  407  VAFPHI--KHNYECLVYVQ-LPFMEDLRQYMFSSLKN-SKKYAPTEAQLNAVDALIDSMS  462
            V  P++  K N     Y   LPF ED+R + F S  N       P E Q +A D L+ +
Sbjct  381  VLTPNVSDKDNTPDSFYFNILPFAEDVRDFQFPSFSNLPSSMQPNEKQQDAADKLVQMLD  440

Query  463  LAKKDEKTDTLEDLFPTTKIPNPRFQRLFQCLLHRALHPREPLPPIQQHIWNMLNPPAEV  522
            LA    K + L  F      PNP  +R ++ L  ++ HP   +PP+ + +  +  P  E+
Sbjct  441  LAPPG-KQEVLSPDF----TPNPVLERYYRYLNLKSKHPDAAVPPLDETLRKITEPDVEL  495

Query  523  TTKSQIPLSKIKTLFPLIEAKKKDQVTAQEIFQDNHEDGPTAKKLKTEQGGAHFSVSSL-  581
             ++++  + +++  F L +  K  + +A+ I     ++ P+    + E+      V ++
Sbjct  496  LSQNKSIIEELRRSFELKDNPKLKK-SARRI-----KERPSGSDEEIEEFNKDADVKAID  549

Query  582  -----AEGSVTSVGSVNPAENFRVLVKQK------KASFEEASNQLINHIEQFLDTNETP  630
              A+  V  VG VNP ++F  ++ ++           + ++  N++ + +E  D +
Sbjct  550  SMEYSAKTEVEKVGDVNPVKDFEDMLSRRDNPKWISKAIQDMKNRIFDLVENSCDGDT--  607

Query  631  YFMKSIDCIRAFREEAIKFSEEQRFNNFLKALQEKVEIKQLNHFWEIVVQDGITLITKEE  690
             F K++ C+ A R+  I   E ++FN+FL  L +  + K L  F   +     ITLITK E
Sbjct  608  -FHKALQCLVALRKGCILEQEPKQFNDFLCHLSKFCQEKDLRSFCLYLTSHEITLITKAE  666

Query  691  ASGSSVTAEEAKKFLAPKDKPSGDTAAVFEEGGDVDDLLDM  731
            A  S ++  EA+ F+    KP  D+  +  E   DD++ +
Sbjct  667  APDSEISEHEARSFMV---KPELDSQNMKSEAKAEDDIMSI  704
```

**Query = hamster KU80; Sbjct = tomato KU80**

**Figure S4. Protein alignment with NCBI blast between the hamster Ku80 and tomato Ku80 peptides.**
The red discontinuous box denotes the starting or KUDN peptide.

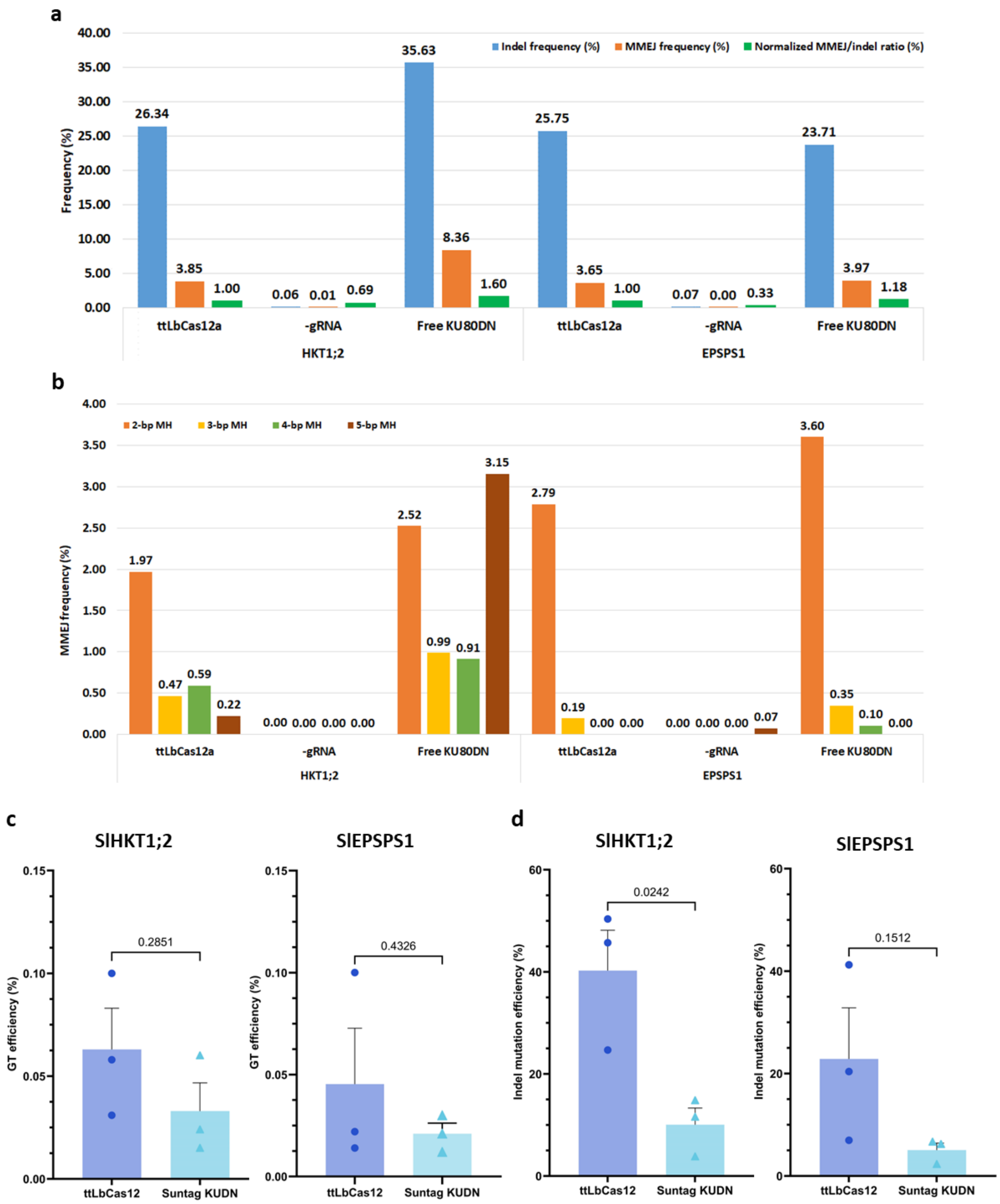


**Figure S5. The impacts of the overexpression of free and Suntag-recruited KUDN on the editing efficiency in tomato at the callus stage. a-b.** The impacts of free KUDN on the editing efficiency (**a**) and the MMEJ efficiency with different microhomology lengths (**b**). **c-d**. The impacts of Suntag-recruited KUDN to the targeted sites on the GT (**c**) and indel mutation (**d**) efficiency at the callus stage, assessed by targeted deep sequencing.

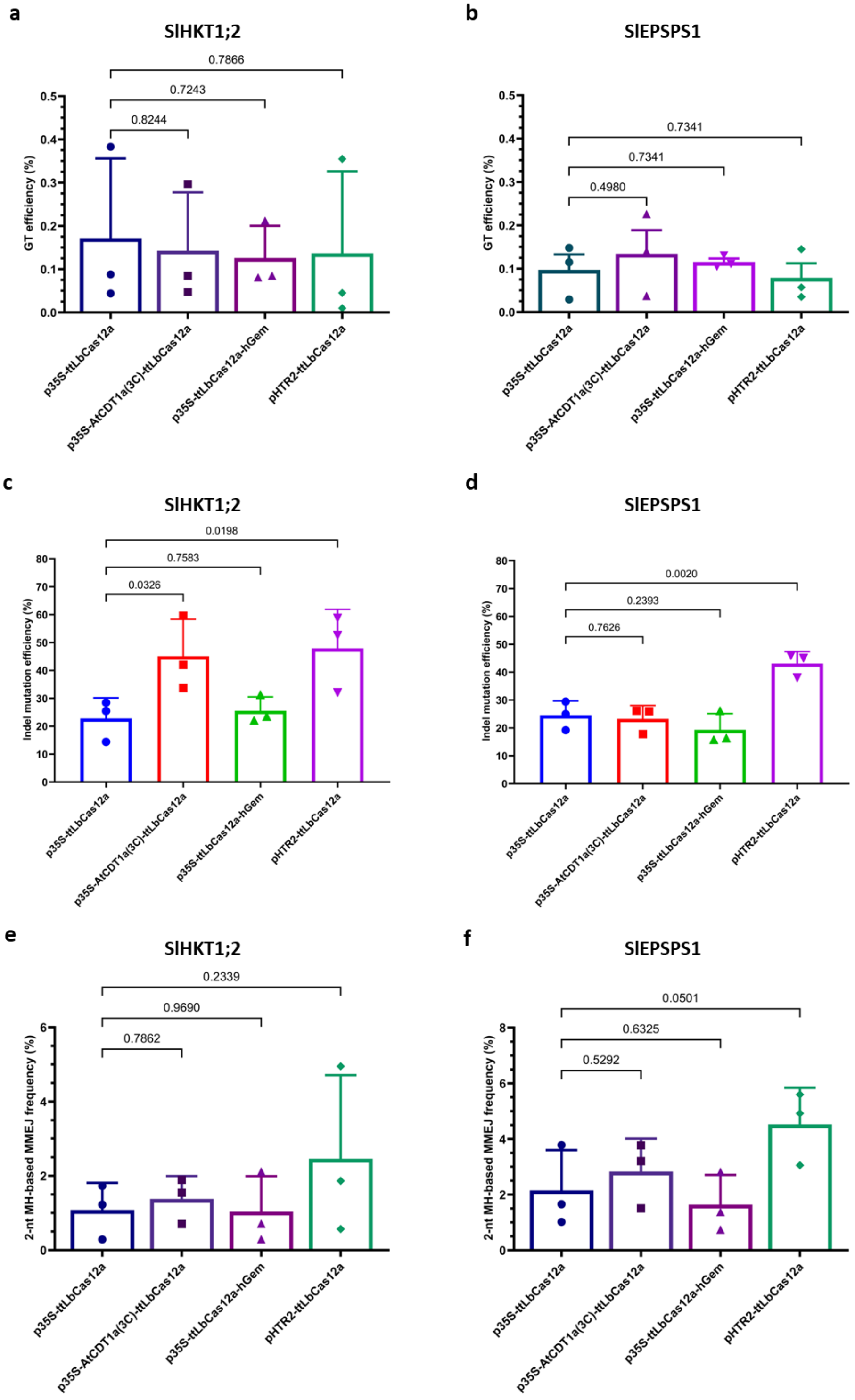


**Figure S6. The impact of cell cycle synchronization on the editing efficiency in tomato. a-c**. The impacts of cell cycle synchronization on GT (**a-b**), indel mutation (**c-d**), and 2-nt MH MMEJ (**e-f**) efficiency at the callus stage. The efficiencies were assessed by targeted NGS using 21-dpt cotyledon/callus samples. The data points are shown as the dots on the plots.

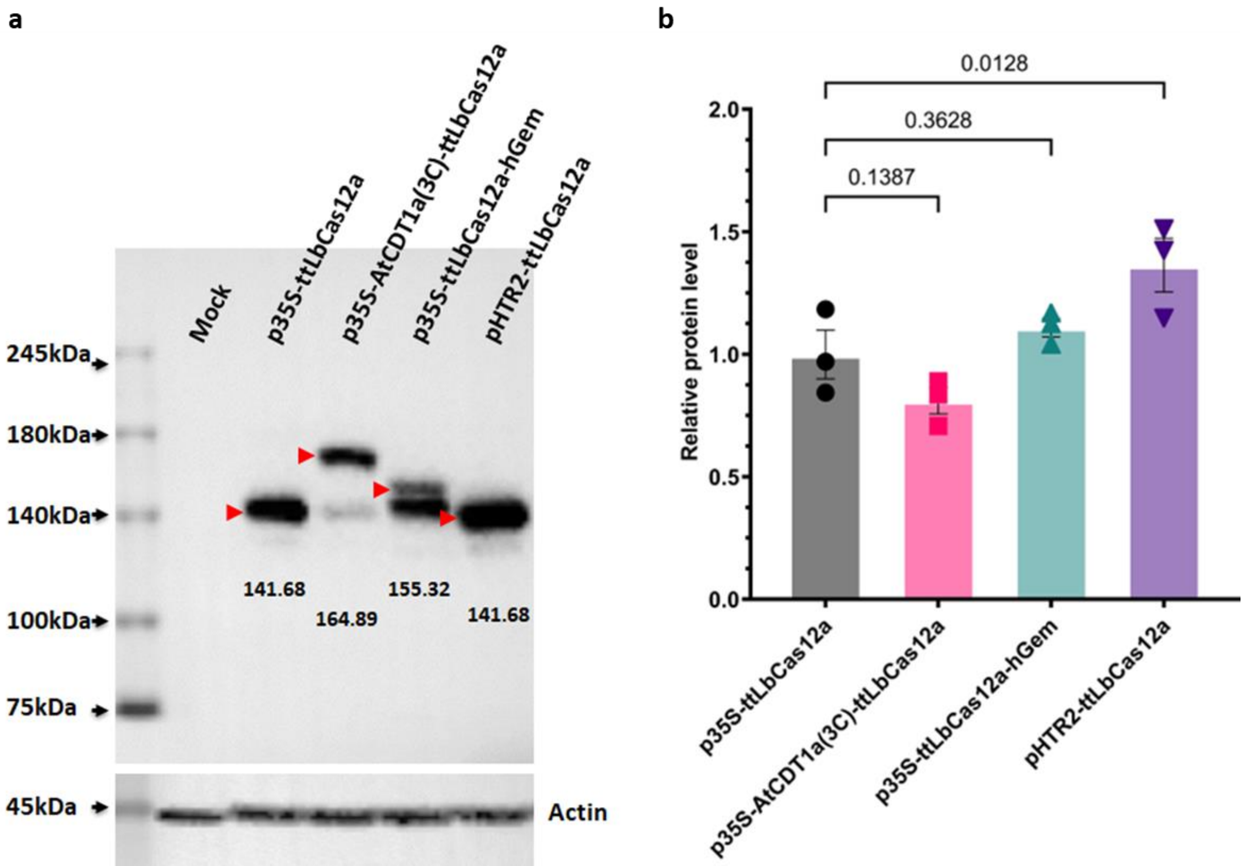


**Figure S7. The protein expression levels of ttLbCas12a and the constructs for cell cycle synchronization.** **a** Western blot membranes showing bands of the Cas proteins with expected sizes and expression cassettes. Actin levels were used as loading controls. The red arrows denote the position of expected bands. **b** Relative protein levels of the Cas protein. P-values of the t-test for pair-wise comparison between the relative protein levels are indicated on the top of the bars. The expression cassettes and Cas configurations are at the bottom of the bars. The data points are shown as the dots on the plots.

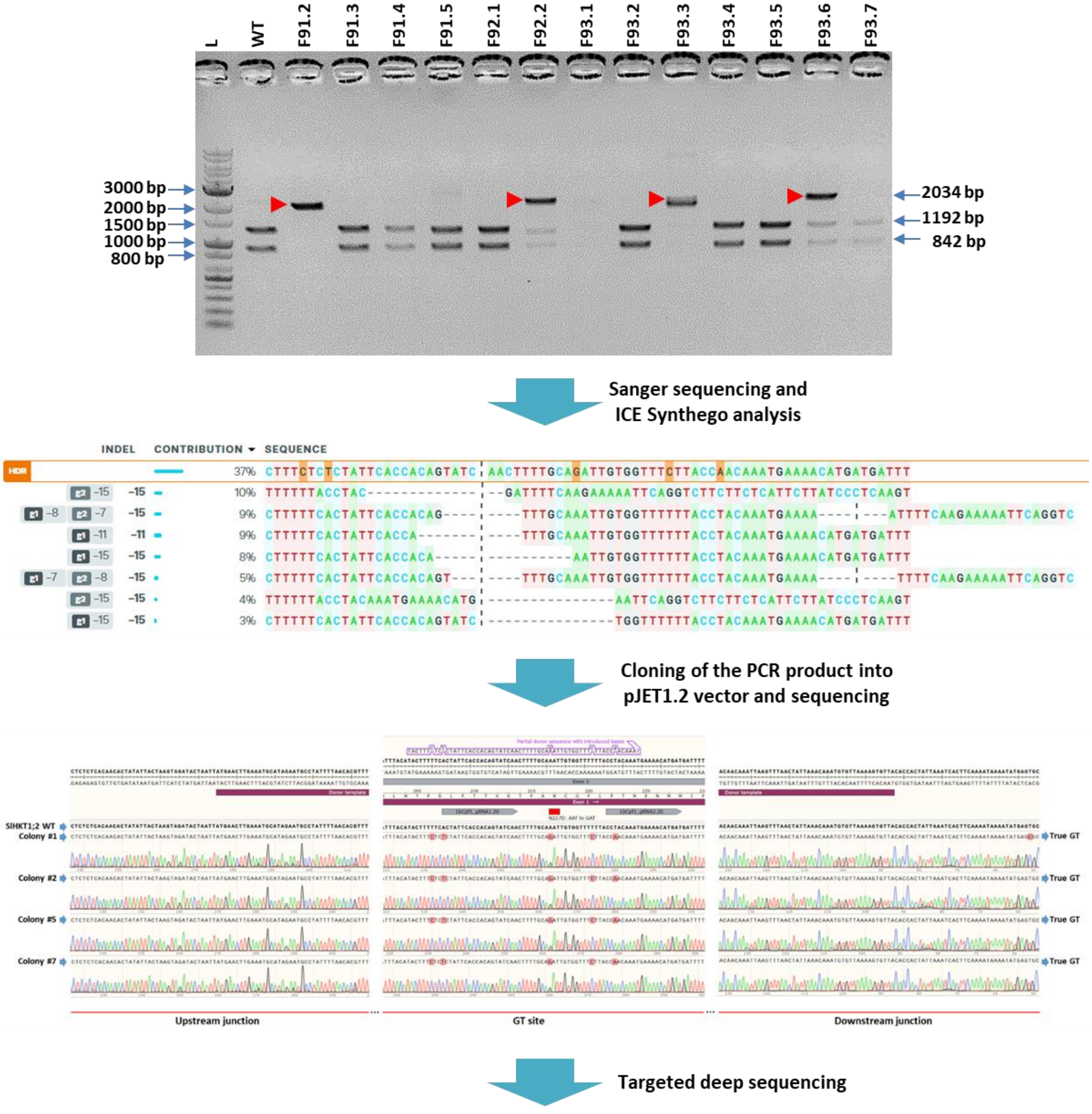


| Total reads | GT frequency (%) | Indel frequency (%) |
|---|---|---|
| 6058 | 50.330 | 48.151 |

**Figure S8. The step-by-step procedure to confirm a GT0 event.** The CAPS assay might first screen transformants. Alternatively, the PCR products amplified from the targeted sites of GT transformants were directly sequenced and analyzed by ICE Synthego. Potential GT events were then validated by (1) cloning the PCR products into pJET1.2 plasmid and sequencing and (2) targeted deep sequencing.

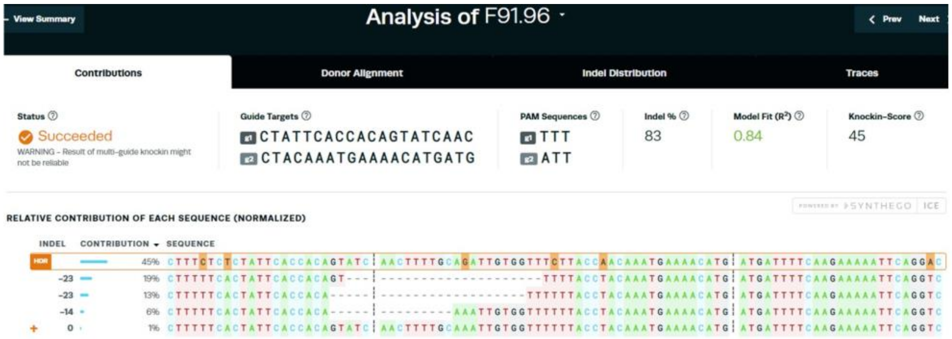


ICE Synthego analysis found the GT allele from the plant

- **pJET1.2 cloning and sequencing**

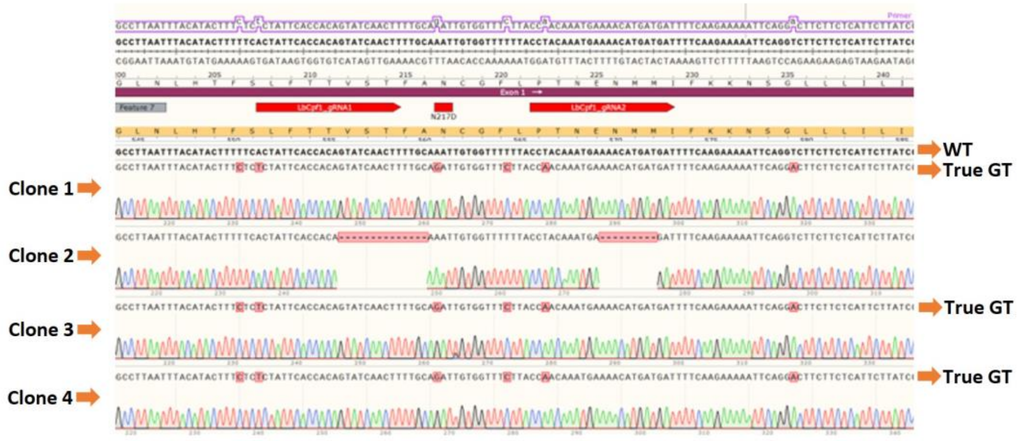


3/4 clones appear to carry perfect GT alleles. Junction sequencing is conducting for validating it.

- **Miniseq data**

| Total Sequences | With both indicator sequences | More than minimum frequency | Insertions | Deletions | Indel frequency | HDR frequency |
|---|---|---|---|---|---|---|
| 124307 | 54646 | 54646 | 0 | 28754 | 28754 (52.6%) | 25029 (45.8%) |

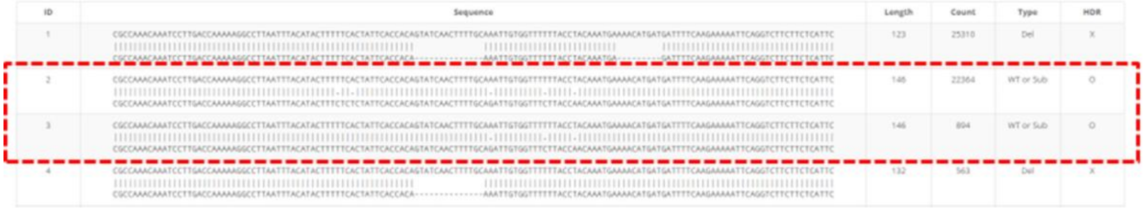


**Figure S9. Identification and validation of the *SlHKT1;2* GT0 event, F91.96.** Top panel: ICE-Synthego analysis identified F91.96 as a potential GT event confirmed by pJET1.2 cloning and sequencing of PCR product and targeted NGS. Sequencing data revealed that 3/4 of colonies contained desired base changes at the GT site and no additional change at the upstream and downstream junctions.

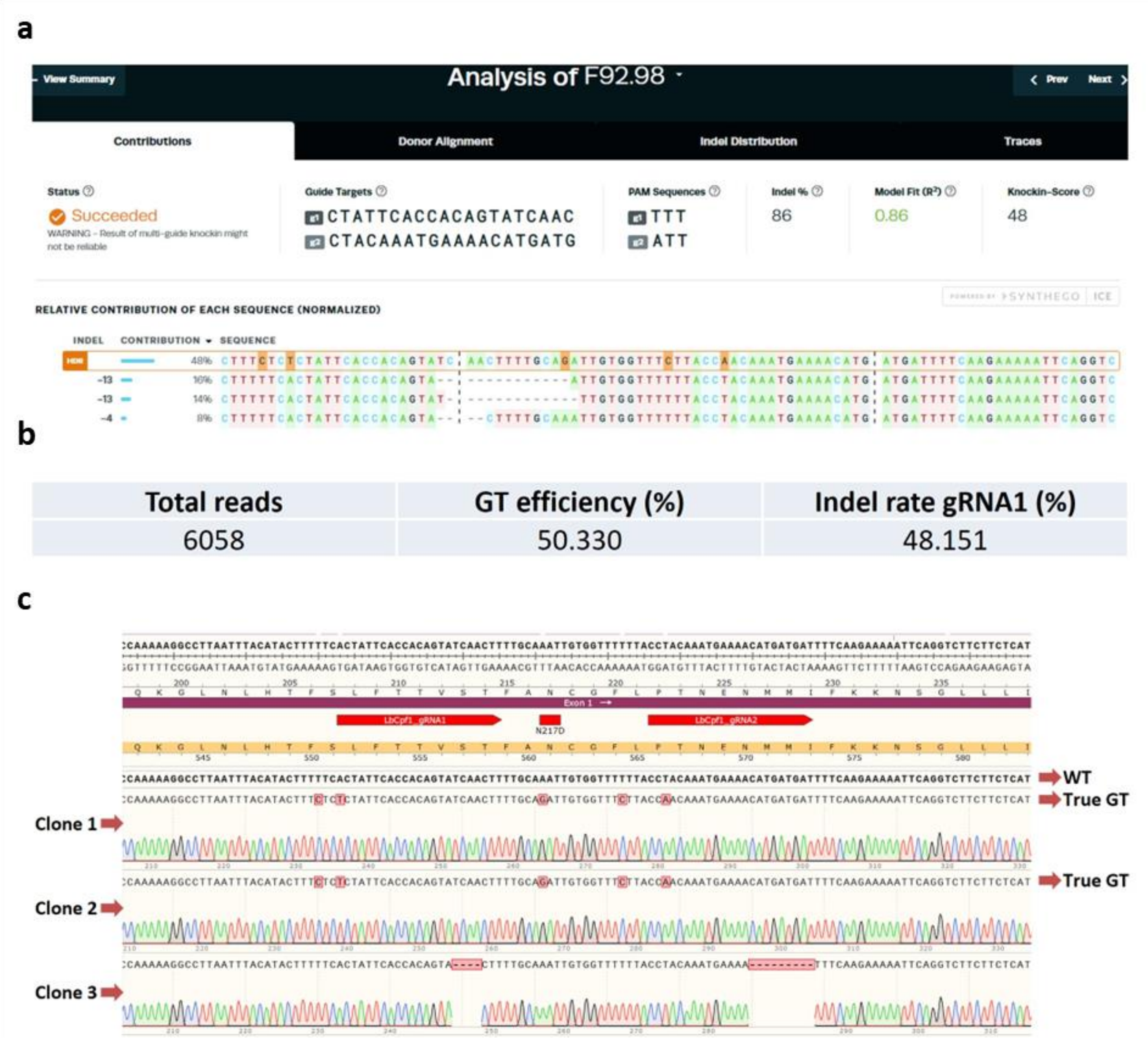


**Figure S10. Identification and validation of the *SlHKT1;2* GT0 event, F92.98, obtained by the NKUDN-GT construct. a** ICE-Synthego analysis identified F92.98 as a potential GT event confirmed by targeted NGS (**b**) and pJET1.2 cloning and sequencing of PCR product (**c**). Sequencing data revealed that 2/3 of colonies contained desired base changes at the GT site and no additional charge at the upstream and downstream junctions.

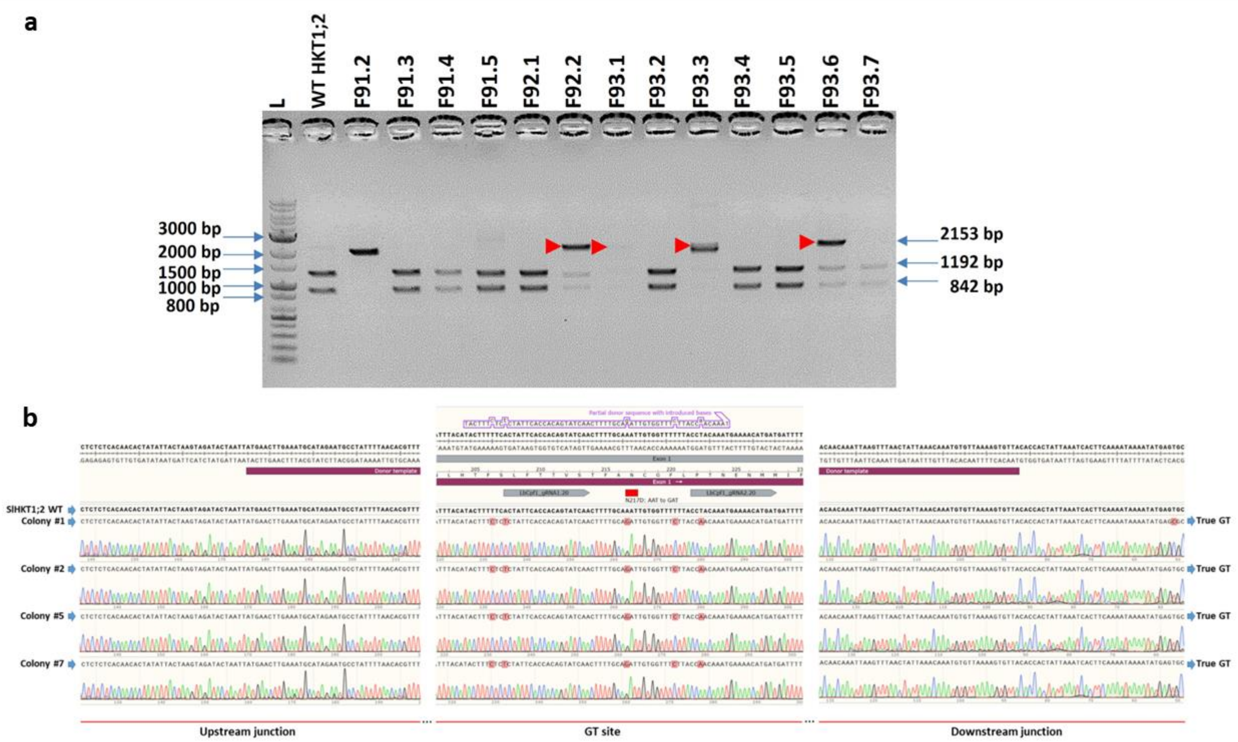


**Figure S11. Identification and validation of the *SlHKT1;2* GT0 event, F93.6, obtained by the CKUDN-GT construct. a** CAPS assay identified F93.6 as a potential GT event. The potential GT bands are denoted with red rectangles. **b** Cloning the F93.6 PCR product into pJET1.2 and sequencing revealed a true GT allele. Sequencing data revealed that 3/7 colonies contained desired base changes at the GT site and no additional change at the upstream and downstream junctions.

- **ICE Synthego analysis**

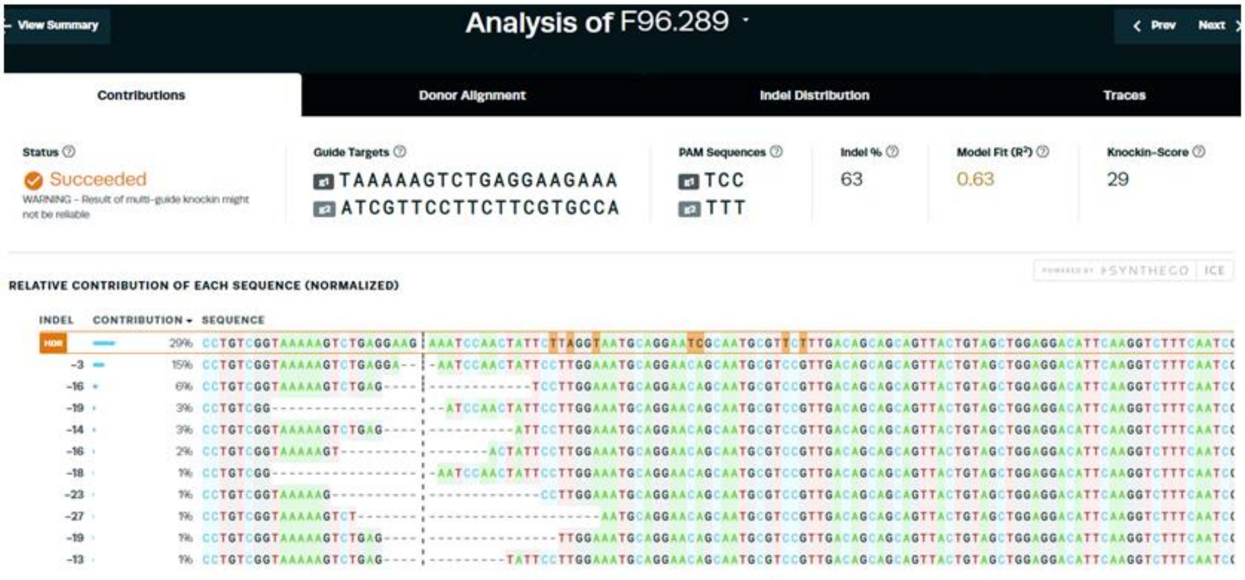


- **Miniseq analysis**

| Plant | Total reads | GT efficiency (%) | Indel rate (%) |
|---|---|---|---|
| F96.289 | 74534 | 44.134 | 51.483 |

**Figure S12. Identification and validation of *SlEPSPS1* GT0 event, F96.289.** Top panel is a ICE-Synthego analysis that identified F96.289 as a potential GT event and subsequent confirmation by targeted NGS in the bottom panel

**a**

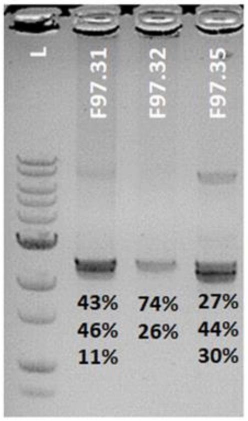


**Calculation of normalized editing efficiency**

| No. | Event | Total miniseq reads | GT efficiency by miniseq (%) | Indel mutation efficiency by miniseq (%) | GT efficiency by Sanger seq, decomposed by ICE Synthego (%) | Approx. GT efficiency normalized to PCR bands (%) | Approx. indel mutation efficiency normalized to PCR bands (%) |
|---|---|---|---|---|---|---|---|
| 1 | F97.31 | 28969 | 39.622 | 59.560 | 11 | 17.1 | 74.0 |
| 2 | F97.32 | 34513 | 28.059 | 71.892 | 6 | 20.6 | 79.2 |
| 3 | F97.35 | 30264 | 99.134 | 0.819 | 16 | 26.4 | 73.2 |

**b**

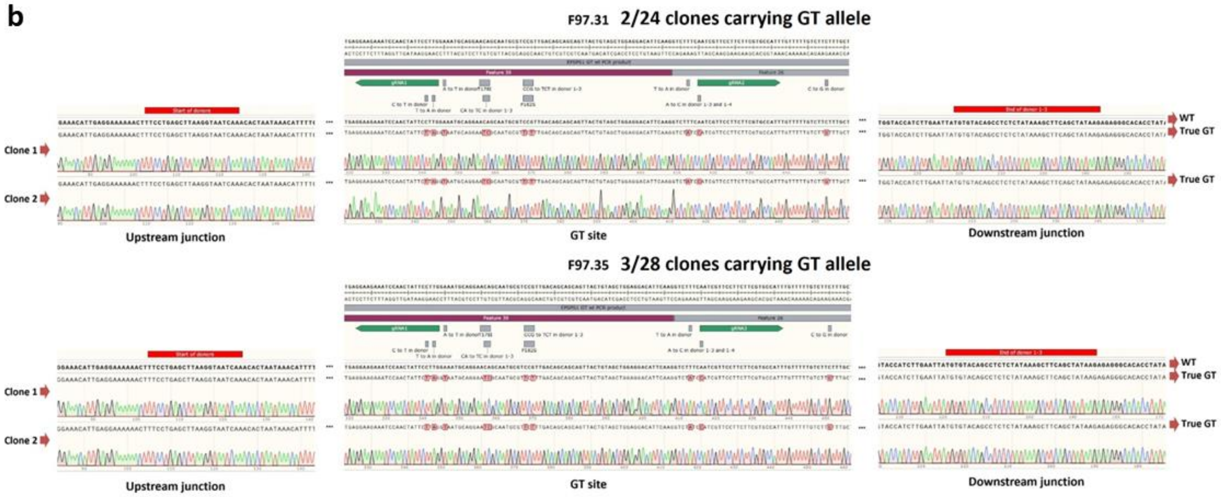


**Figure S13. Identification and validation of the *SlEPSPS1* GT events, F97.31, F97.32 and F97.35, obtained by the NKUDN-GT construct. a** Calculation of normalized GT frequency. Because the PCR product contains a mixture of WT-sized and indel bands, only the upper-most bands corresponding to WT-sized bands were considered to contain true GT alleles. The second PCR reaction for targeted NGS sample preparation only amplified the WT-sized bands so in this case, the miniseq GT frequency did not reflect the true GT frequency. The targeted NGS (miniseq) results and resolved bands on agarose gel were used to identify the true GT frequency. The PCR bands on the agarose gel were measured by ImageJ and normalized to the band size to calculate the band frequency.. **b** The potential *SlEPSPS1* GT events, F97.31 and F97.35, were confirmed by pJET1.2 cloning and sequencing that show true GT allele carried by 2/24 clones (F97.31) and 3/28 clones (F97.35). Sequencing data revealed that 3/7 colonies contained desired base changes at the GT site and no additional change at the upstream and downstream junctions.

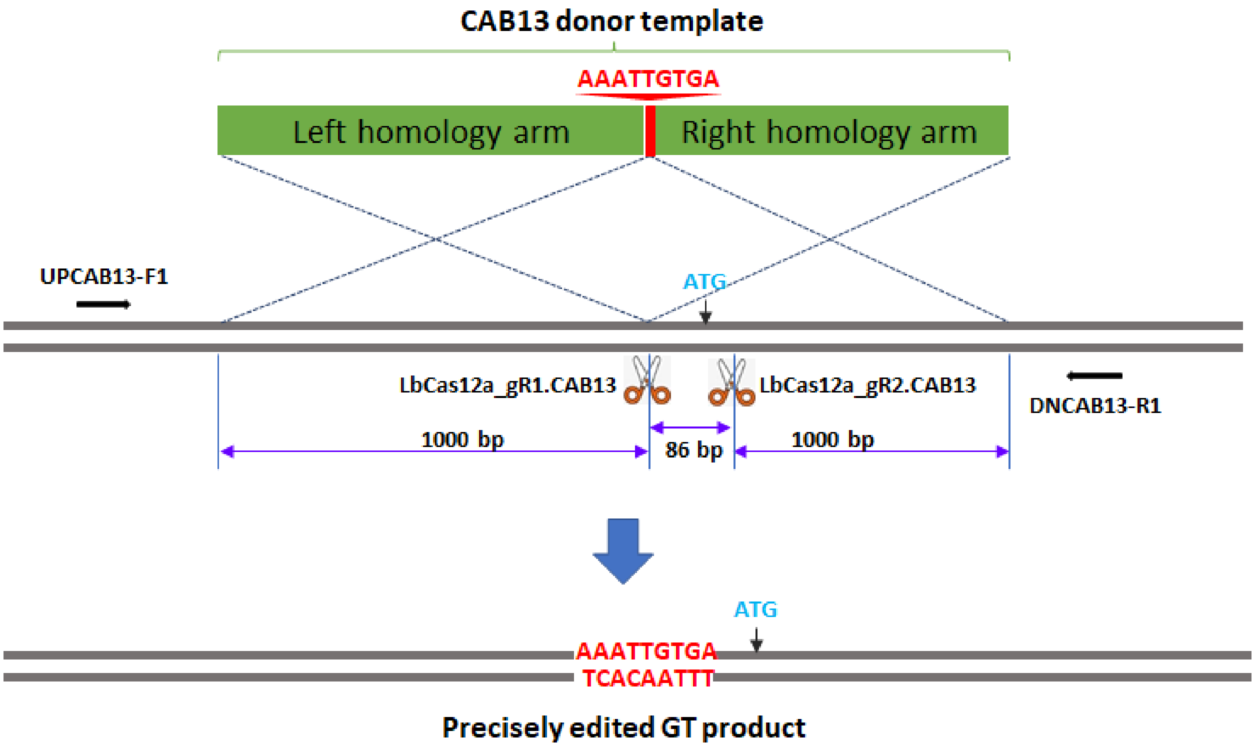


**Figure S14. The donor and mode of allele replacement of the *SlCAB13*.** Schematic diagrams describing the expected GT processes for exchanging the homologous DNA donor template with the genomic sequence at the *SlCAB13* locus. During the cloning of the *SlCAB13* donor, the AAATTGTGA sequence was introduced for integration into the genomic site. Two designated cutting sites were strategically planned to induce double-strand breaks at the targeted sites. The reverse and forward primers, represented by black arrows, are designed to amplify the targeted sites using a polymerase chain reaction. The diagrams were drawn not to their actual scales.

a

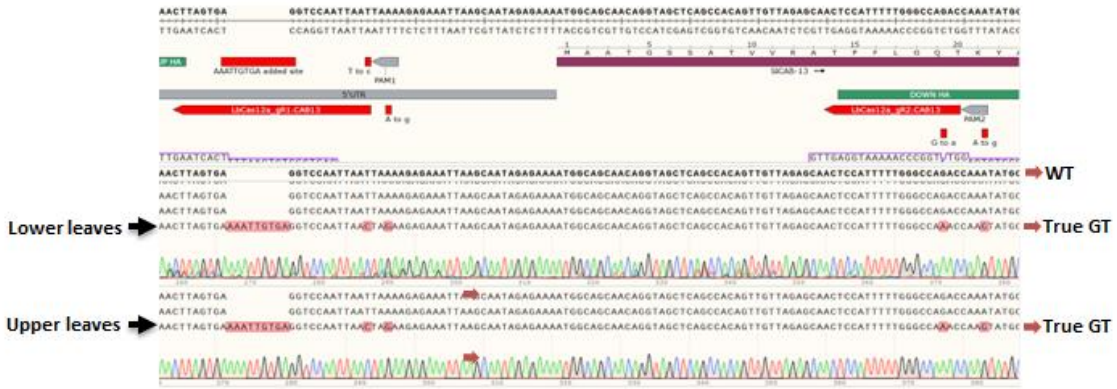


b

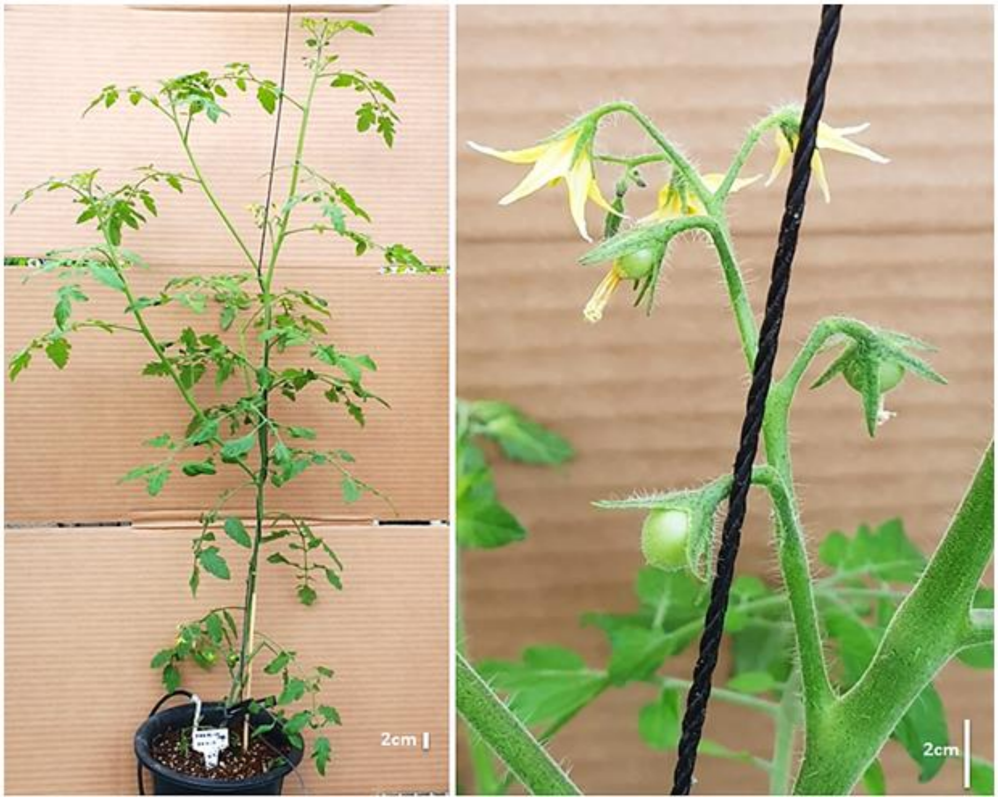


**Figure S15. Design and evaluation of KUDN-based GT tools for *SlCAB13* editing. a** Sequencing data revealed that 3/21 (~14.2%) colonies contained desired base changes at the GT site and no additional change at the upstream and downstream junctions. **b** The *SlCAB13* GT0 event planting in the greenhouse showed normal morphology during vegetative and reproductive stages.

a

b

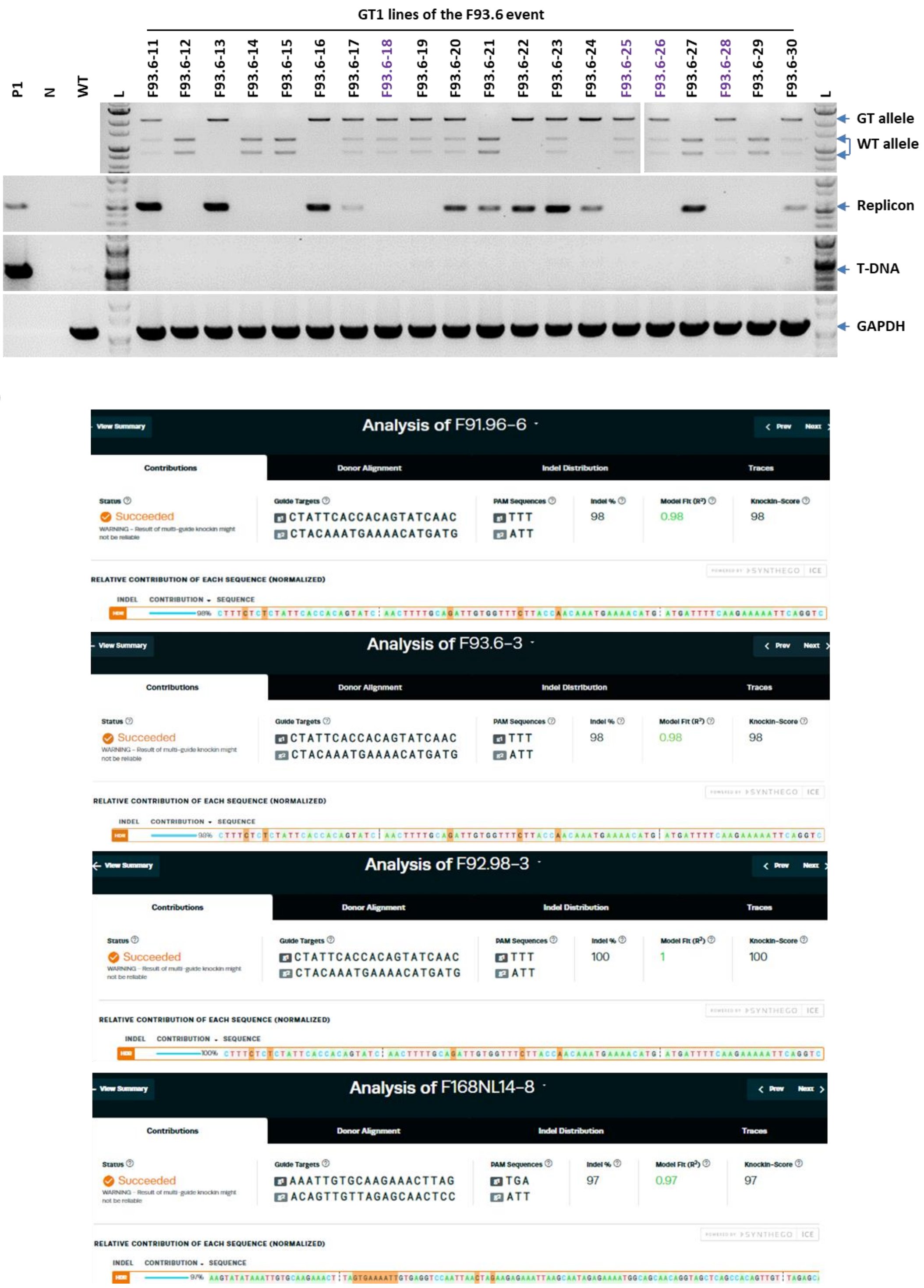


**Figure S16. The inheritance and segregation of GT alleles. a** The inheritance of *SlHKT1;2* GT allele in GT1 generation. CAPS assay was used to analyze the GT1 samples. **b** Representative plants carry homozygous GT alleles of *SlHKT1;2* (three above panels) and *SlCAB13* (bottom panel) revealed by sequencing and ICE Synthego analysis.

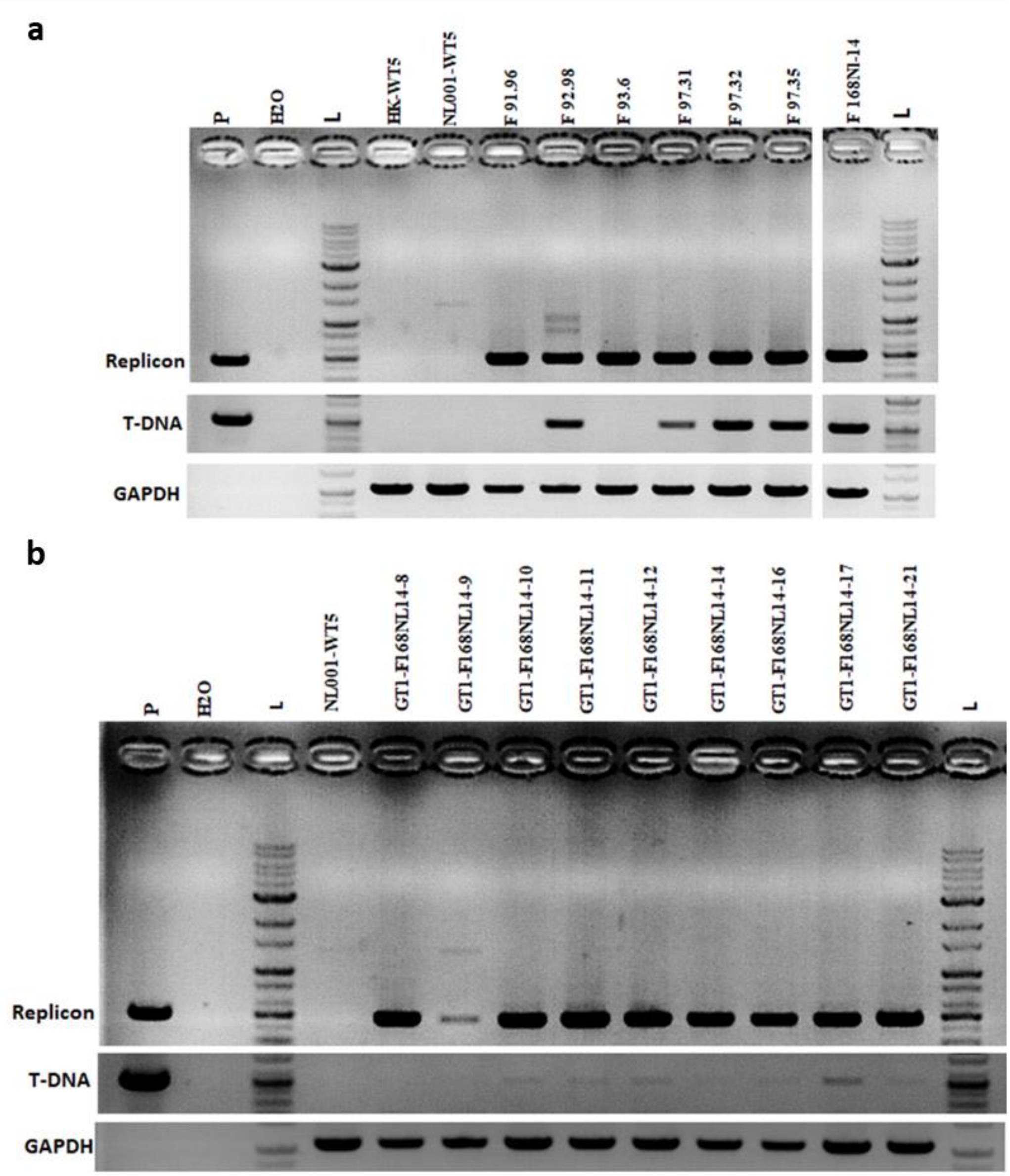


**Figure S17. PCRs assessed T-DNA and replicon in the GT plants. a** Representative GT0 events obtained in the study. **b** GT1 plants of the *SlCAB13* GT event F168NL14. The plasmids isolated from agro clones that carried the GT tool used for generating the events are referred to as positive control (P). H20 denotes the water control, while HK-WT5 and NL001-WT5 represent Hongkwang wild-type and NL001 wild-type, respectively. *SlHKT1;2* GT0 events were obtained using the GT construct without KUDN, with NKUDN, and with CKUDN, respectively, in Hongkwang's background and are referred to as F91.96, F92.98, and F93.6. Similarly, *SlEPSPS1* GT0 events were obtained using the GT construct with NKUDN in Hongkwang's background and are referred to as F97.31, F97.32, and F97.35. The *SlCAB13* GT0 event was obtained using the GT construct with CKUDN in the NL001 background and is referred to as F168NL14. GT1-F168NL14-8, 9, 10, 11, 12, 14, 16, 17, and 21 are GT1 plants carrying the homozygous *SlCAB13* GT allele.

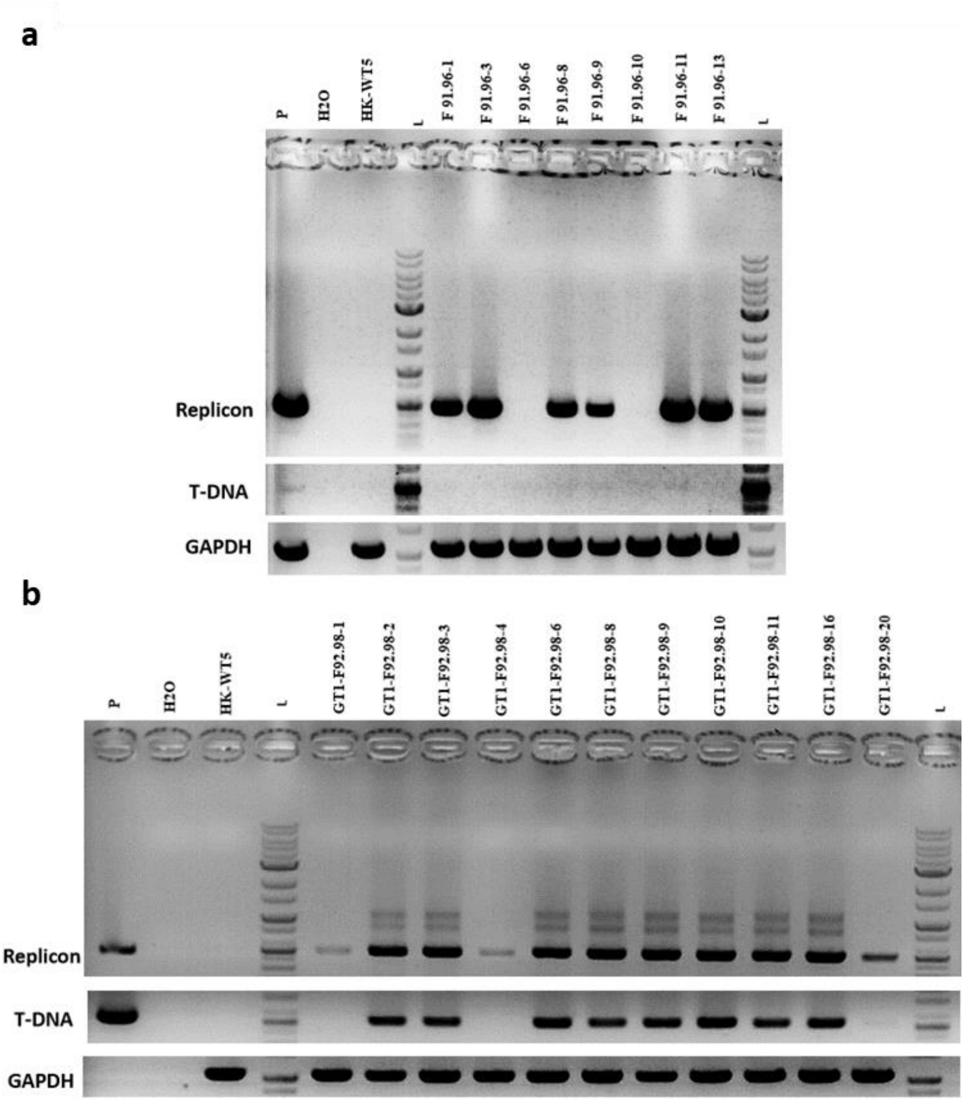


**Figure S18. Assessment of T-DNA and replicon in the *SlHKT1;2* GT1 plants.** **a** GT1 plants of the F91.96 *SlHKT1;2* GT0 event (without KUDN). **b** GT1 plants of the F92.98 *SlHKT1;2* GT0 event (generated by the NKUDN tool). P: The *SlHKT1;2* GT plasmid isolated from Agro clones used for generating the events. The water control was labeled H2O, while the WT control was labeled HK-WT5. The F92.98-GT1 homozygous plants were labeled F92.98-3, 4, 10, 16, and 20. The F92.98-GT heterozygous plants were labeled as F92.98-1-2-6-8-9-11. Similarly, the F91.96-GT1 homozygous plants of the F91.96 GT0 event were labeled as F91.96-6, 11, and 13, while the F92.98-GT1 homozygous plants of the F92.98 GT0 event were labeled as F92.98-1, 2, 3, 4, 6, 8, 9, 10, 11, 16, and 20.

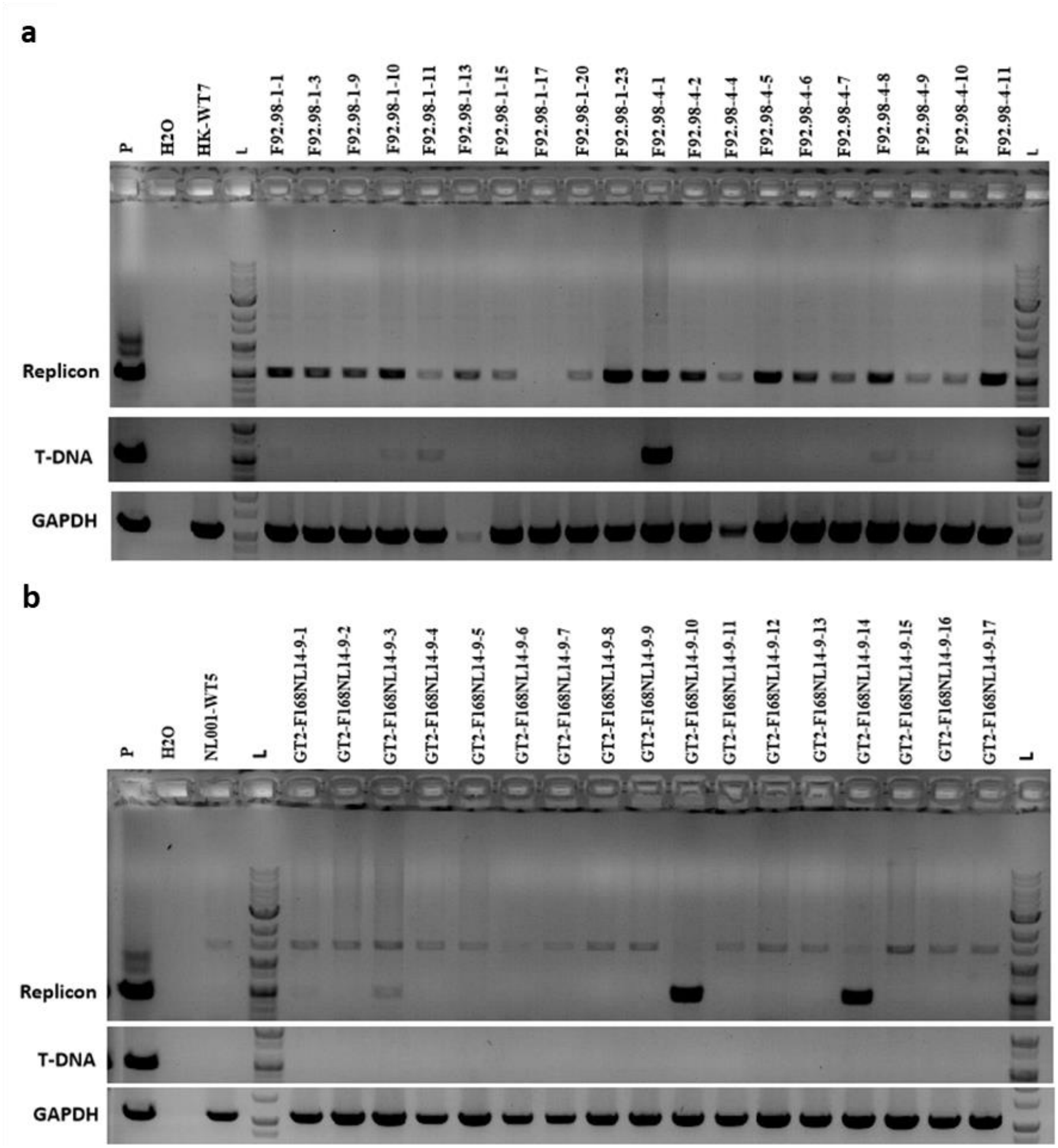


**Figure S19. PCR-based assessment of T-DNA and replicon in the *SlHKT1;2* and *SlCAB13* GT2 plants. a** GT2 homozygous plants of the *SlHKT1;2* (with NKUDN) GT1 line, F92.98-1. **b** GT2 homozygous plants of the *SlCAB13* GT1 line, F168NL14-9. The positive control (P) was *SlHKT1;2* GT plasmid isolated from the agrobacterium clone that generated the event F92.98; water control was labeled H2O; and Hongkwang wild-type control was labeled HK-WT5 while NL001 wild-type plants were marked as NL001-WT5. Samples of GT2 homozygous plants from the F92.98-1 GT1 line were marked as F92.98-1-1, 3, 9, 11, 13, 15, 17, 20, and 23, and samples of GT2 homozygous plants from the F92.98-4 GT1 line were marked as F92.98-4-1, 2, 4, 5, 6, 7, 8, 9, and 11. The GT2 homozygous plants from the F168NL14-9 GT1 line were marked as GT2-F168NL14-9-1, 2, 3, 4, 5, 6, 7, 8, 9, 10, 11, 12, 13, 14, 15, 16, and 17.

**a**

| Locus | gRNA | Target | Chromosome | Position | Direction | Mismatches |
|---|---|---|---|---|---|---|
| EPSPS1 | 1 | crRNA: TTTNCAAGGAATAGTTGGATTTCTTCC<br>DNA: TTTCCAAGGAAaAGTTGaATcTCTTCC | SL4.0ch05 | 60582371 | - | 3 |
| CAB13 | 1 | crRNA: TTTNATTAATTGGACCTCACTAAG<br>DNA: TTTAATTAATTGGcCCTCACTAca | SL4.0ch01 | 56418252 | + | 3 |
| | 2 | crRNA: TTTNGTCTGGCCCAAAAATGGAGT<br>DNA: TTTGGTtTGGCCCAAAAATGGtGT | SL4.0ch12 | 4306011 | - | 2 |

**b**

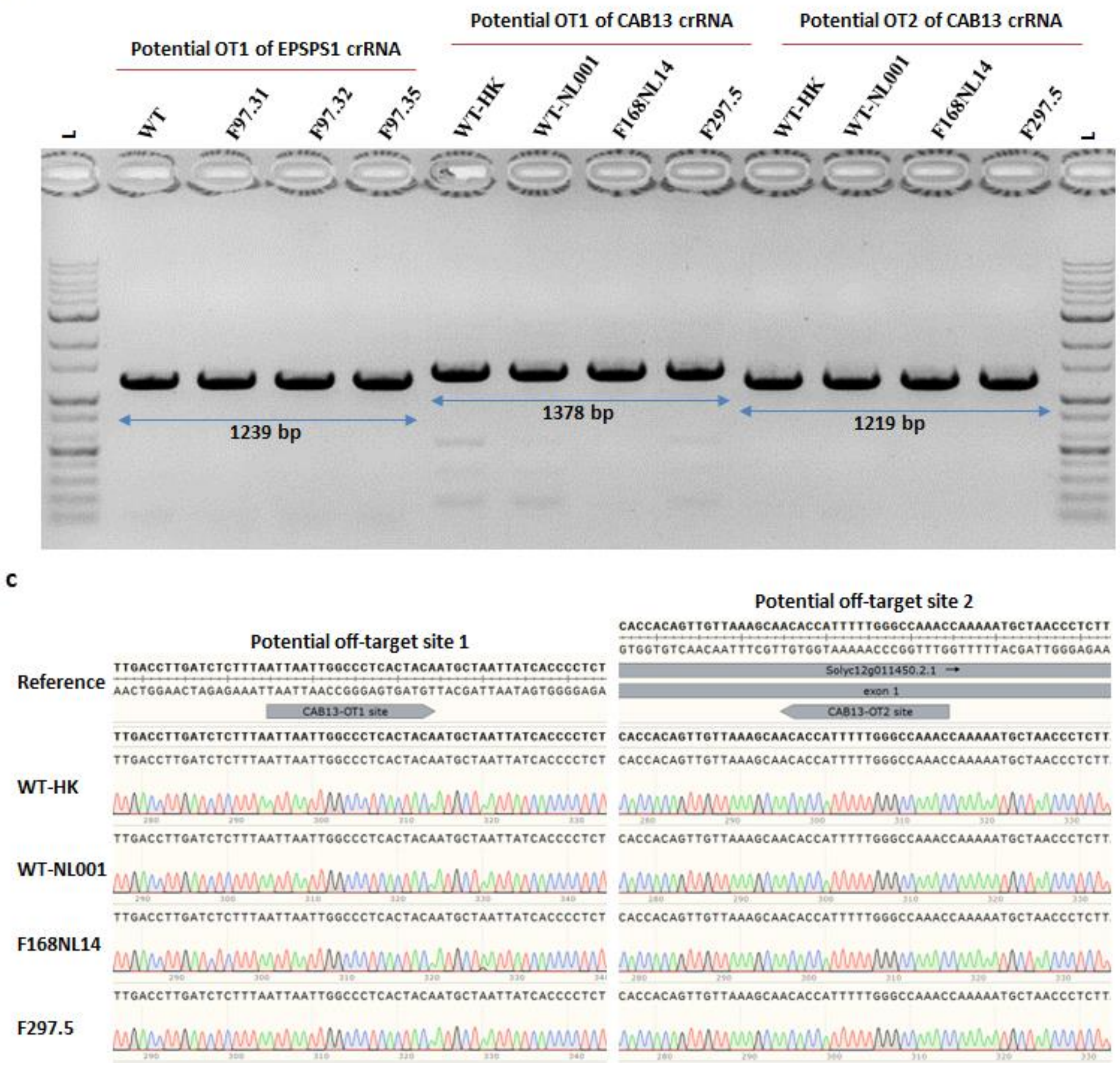


**Figure S20. Assessment of potential off-target activities of the KUDN-based GT tools in the GT events.** **a** Potential off-target sites of the gRNAs identified by Cas-offinder. **b** PCR products amplified from the selected potential off-target sites of *SlEPSPS1* and *SlCAB13* gRNAs in the GT events F97.31, F97.32, F97.35, and F168NL14 and other potential GT events (*SlEPSPS1*) and F297.5 (*SlCAB13*). **c** Chromatograms of the sequenced data show no editing traces at the potential off-target sites of *SlCAB13* gRNAs.

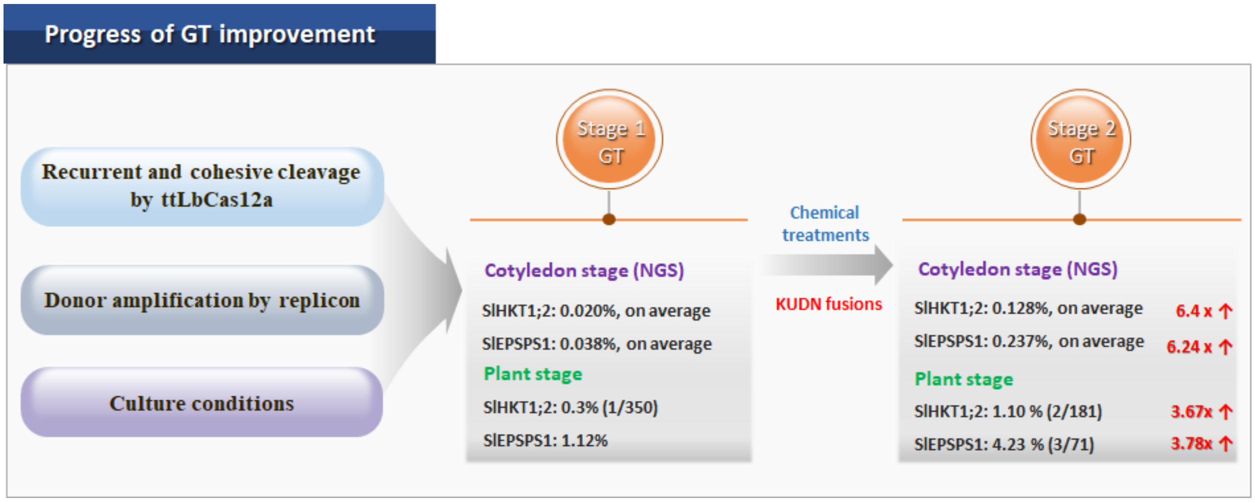


**Figure S21. The GT progress is updated with this study.** Previous stage GT study (Stage 1 GT) revealed the improvement of GT with the ttLbCas12a, donor amplification by the geminiviral replicon, favorable culture conditions, and chemical treatments. In this upgraded version (Stage 2 GT), the GT system has been updated with the addition of NKUDN that supported up to 6.4-fold and 3.78-fold enhancement of GT efficiency at the callus and plant stage, respectively.

### Supplemental Tables

**Table S1. The impacts of nuclease fusions on microhomology length and frequency**

| No. | Targeted locus | Construct | Total read | Indel frequency (%) | MMEJ read | MMEJ frequency (%) | MMEJ /indel ratio | Total reads | | | | | Frequency (%) | | | | | Normalized MMEJ/indel ratio |
|---|---|---|---|---|---|---|---|---|---|---|---|---|---|---|---|---|---|---|
| | | | | | | | | 2-bp MH | 3-bp MH | 4-bp MH | 5-bp MH | 6-bp MH | 2-bp MH | 3-bp MH | 4-bp MH | 5-bp MH | 6-bp MH | |
| 1 | *SlHKT1;2* | ttLbCas12a | 12990 | 33.53 | 500 | 3.85 | 0.115 | 165 | 57 | 167 | 44 | 29 | 1.27 | 0.44 | 1.29 | 0.34 | 0.22 | 1.00 |
| 2 | | hExo1a-ttLbCas12a | 19229 | 7.66 | 217 | 1.13 | 0.147 | 43 | 57 | 24 | 56 | 0 | 0.22 | 0.30 | 0.12 | 0.29 | 0.00 | 1.28 |
| 3 | | T5exo-ttLbCas12a | 16840 | 44.26 | 2221 | 13.19 | 0.298 | 1042 | 222 | 574 | 106 | 29 | 6.19 | 1.32 | 3.41 | 0.63 | 0.17 | 2.60 |
| 4 | | ttLbCas12a-hHE | 13070 | 44.30 | 749 | 5.73 | 0.129 | 223 | 12 | 296 | 126 | 84 | 1.71 | 0.09 | 2.26 | 0.96 | 0.64 | 1.13 |
| 5 | | ttLbCas12a-CtIP | 12348 | 17.82 | 1029 | 8.33 | 0.468 | 30 | 34 | 823 | 90 | 45 | 0.24 | 0.28 | 6.67 | 0.73 | 0.36 | 4.07 |
| 6 | *SlEPSPS1* | ttLbCas12a | 22097 | 20.46 | 741 | 3.35 | 0.164 | 710 | 31 | 0 | 0 | 0 | 3.21 | 0.14 | 0.00 | 0.00 | 0.00 | 1.00 |
| 7 | | hExo1a-ttLbCas12a | 21571 | 4.80 | 178 | 0.83 | 0.172 | 141 | 37 | 0 | 0 | 0 | 0.65 | 0.17 | 0.00 | 0.00 | 0.00 | 1.05 |
| 8 | | T5exo-ttLbCas12a | 26578 | 13.09 | 847 | 3.19 | 0.243 | 736 | 111 | 0 | 0 | 0 | 2.77 | 0.42 | 0.00 | 0.00 | 0.00 | 1.48 |
| 9 | | ttLbCas12a-hHE | 25111 | 10.98 | 505 | 2.01 | 0.183 | 462 | 24 | 0 | 0 | 9 | 1.84 | 0.10 | 0.00 | 0.00 | 0.04 | 1.12 |
| 10 | | ttLbCas12a-CtIP | 22365 | 2.84 | 162 | 0.72 | 0.255 | 105 | 57 | 0 | 10 | 0 | 0.47 | 0.25 | 0.00 | 0.04 | 0.00 | 1.55 |

**Table S2. The impacts of overexpression of free KU80DN on microhomology length and frequency**

| No. | Targeted locus | Construct | Total read | Indel frequency (%) | MMEJ read | MMEJ frequency (%) | MMEJ/indel ratio (%) | Total reads | | | | | Frequency (%) | | | | | Normalized MMEJ/indel ratio (%) |
|---|---|---|---|---|---|---|---|---|---|---|---|---|---|---|---|---|---|---|
| | | | | | | | | 2-bp MH | 3-bp MH | 4-bp MH | 5-bp MH | 6-bp MH | 2-bp MH | 3-bp MH | 4-bp MH | 5-bp MH | 6-bp MH | |
| 1 | *SlHKT1;2* | ttLbCas12a | 194596 | 26.34 | 7495 | 3.85 | 14.62 | 4007 | 953 | 1204 | 453 | 410 | 1.97 | 0.47 | 0.59 | 0.22 | 0.20 | 1.00 |
| 2 | | ttLbCas12a, -gRNA | 190331 | 0.06 | 12 | 0.01 | 10.04 | 7 | 1 | 1 | 3 | 0 | 0.00 | 0.00 | 0.00 | 0.00 | 0.00 | 0.69 |
| 3 | | ttLbCas12a, free KU80DN | 195458 | 35.63 | 16341 | 8.36 | 23.46 | 5294 | 2073 | 1915 | 6615 | 126 | 2.52 | 0.99 | 0.91 | 3.15 | 0.06 | 1.60 |
| 4 | *SlEPSPS1* | ttLbCas12a | 141624 | 25.75 | 5169 | 3.65 | 14.18 | 4716 | 328 | 0 | 0 | 0 | 2.79 | 0.19 | 0.00 | 0.00 | 0.00 | 1.00 |
| 5 | | ttLbCas12a, -gRNA | 188699 | 0.07 | 6 | 0.00 | 4.69 | 4 | 2 | 0 | 125 | 0 | 0.00 | 0.00 | 0.00 | 0.07 | 0.00 | 0.33 |
| 6 | | ttLbCas12a, free KU80DN | 168199 | 23.71 | 6672 | 3.97 | 16.73 | 5840 | 561 | 168 | 0 | 0 | 3.60 | 0.35 | 0.10 | 0.00 | 0.00 | 1.18 |

**Table S3. The impacts of the recruitment of KU80DN to the targeted sites by the Suntag system on GT and indel efficiency**

| No. | Targeted locus | Construct | Rep 1 | | | Rep 2 | | | Rep3 | | | Average | | | SEM | |
|---|---|---|---|---|---|---|---|---|---|---|---|---|---|---|---|---|
| | | | Total reads | GT efficiency (%) | Indel rate gRNA1 (%) | Total reads | GT efficiency (%) | Indel rate gRNA1 (%) | Total reads | GT efficiency (%) | Indel rate gRNA1 (%) | Total reads | GT efficiency (%) | Indel rate gRNA1 (%) | GT efficiency (%) | Indel rate gRNA1 (%) |
| 1 | *SlHKT1;2* | ttLbCas12a | 41385 | 0.031 | 24.678 | 97080 | 0.10 | 45.71 | 121976 | 0.058 | 50.353 | 260441 | 0.063 | 40.247 | 0.020 | 7.899 |
| 3 | | ttLbCas12a + 10xsuntag-KU80DN | 96018 | 0.024 | 3.799 | 87546 | 0.06 | 14.80 | 109884 | 0.015 | 11.565 | 293448 | 0.031 | 10.055 | 0.013 | 3.264 |
| 4 | *SlEPSPS1* | ttLbCas12a | 55518 | 0.022 | 6.976 | 42647 | 0.10 | 41.22 | 134686 | 0.014 | 20.392 | 232851 | 0.045 | 22.862 | 0.027 | 9.961 |
| 6 | | ttLbCas12a + 10xsuntag-KU80DN | 83297 | 0.012 | 2.295 | 60183 | 0.03 | 6.65 | 77297 | 0.021 | 6.197 | 220777 | 0.020 | 5.048 | 0.005 | 1.383 |

**Table S4. The impacts of synchronization toward S-G2 phases on microhomology length and frequency**

| Locus | Construct | Total read | Indel frequency (%) | MMEJ read | MMEJ frequency (%) | MMEJ/indel ratio | Total reads | | | | | Frequency (%) | | | | | Normalized MMEJ/indel ratio |
|---|---|---|---|---|---|---|---|---|---|---|---|---|---|---|---|---|---|
| | | | | | | | 2-bp MH | 3-bp MH | 4-bp MH | 5-bp MH | 6-bp MH | 2-bp MH | 3-bp MH | 4-bp MH | 5-bp MH | 6-bp MH | |
| *SlHKT1;2* | p35S-ttLbCas12a | 203401 | 16.99 | 4385 | 2.16 | 0.09 | 2477 | 196 | 1365 | 144 | 203 | 1.22 | 0.10 | 0.67 | 0.07 | 0.10 | 1.00 |
| | p35S-AtCDT1a(3C)-ttLbCas12a | 202361 | 33.54 | 8625 | 4.26 | 0.09 | 3071 | 1693 | 1216 | 2401 | 172 | 1.52 | 0.84 | 0.60 | 1.19 | 0.08 | 1.00 |
| | p35S-ttLbCas12a-Gem | 209717 | 18.90 | 5006 | 2.39 | 0.09 | 2350 | 592 | 1420 | 389 | 168 | 1.12 | 0.28 | 0.68 | 0.19 | 0.08 | 0.99 |
| | pHTR2-ttLbCas12a | 197688 | 32.96 | 12954 | 6.55 | 0.14 | 4829 | 1205 | 4758 | 891 | 644 | 2.44 | 0.61 | 2.41 | 0.45 | 0.33 | 1.44 |
| *SlEPSPS1* | p35S-ttLbCas12a | 169151 | 24.54 | 5074 | 3.00 | 0.12 | 4092 | 178 | 108 | 561 | 135 | 2.42 | 0.11 | 0.06 | 0.33 | 0.08 | 1.00 |
| | p35S-AtCDT1a(3C)-ttLbCas12a | 168472 | 23.26 | 6060 | 3.60 | 0.15 | 5151 | 225 | 684 | 0 | 0 | 3.06 | 0.13 | 0.40 | 0.00 | 0.00 | 1.27 |
| | p35S-ttLbCas12a-Gem | 162169 | 19.32 | 3270 | 2.02 | 0.10 | 3001 | 516 | 28 | 0 | 0 | 1.85 | 0.32 | 0.02 | 0.00 | 0.00 | 0.85 |
| | pHTR2-ttLbCas12a | 162723 | 43.09 | 8440 | 5.19 | 0.12 | 7712 | 571 | 56 | 155 | 0 | 4.74 | 0.35 | 0.03 | 0.10 | 0.00 | 0.98 |

**Table S5. The targeted loci and gRNAs used in the study**

| No. | Locus | Accession | Primer name | Sequence (5' - 3') |
|---|---|---|---|---|
| 1 | *SlHKT1;2* | Solyc07g014680 | LbCpf1_gR1.HKT12 | ACTATTCACCACAGTATCAACTT |
| 2 | | | LbCpf1_ gR2.HKT12 | CCTACAAATGAAAACATGATGAT |
| 3 | *SlEPSPS1* | Solyc01g091190 | LbCpf1_ gR1.EPSPS1 | CAAGGAATAGTTGGATTTCTTCC |
| 4 | | | LbCpf1_ gR2.EPSPS1 | AATCGTTCCTTCTTCGTGCCATT |
| 5 | *SlCAB13* | Solyc07g063600 | LbCas12a_gR1.CAB13 | ATTAATTGGACCTCACTAAG |
| 6 | | | LbCas12a_gR2.CAB13 | GTCTGGCCCAAAAATGGAGT |

**Table S6. Primer sequence used in this study.**

| No. | Loci | Primer name | Sequence (5' - 3') | Note |
|---|---|---|---|---|
| 1 | *SlHKT1;2* | UPHKT12-F1 | TTCACATGCTTTGACCCATAAA | 1st PCR |
| 2 | | DNHKT12-R1 | CTCTTCCTATAAACGTGCACTCA | |
| 3 | | nHKT12-F2 | ACACTCTTTCCCTACACGACGCTCTTCCGATCTCCCTAGCGCCAAACAAATC | 2nd PCR |
| 4 | | nHKT12-R2 | GTGACTGGAGTTCAGACGTGTGCTCTTCCGATCTGGGATAAGAATGAGAAGAAGACCTGAATT | |
| 5 | | HKT12-sF1 | CAAAGATTATGAGCTAGGGAATGT | Sequencing primer |
| 6 | *SlEPSPS1* | UPEPSPS1-F2 | ACATGTAAGTTAGACAAGAGCTAGG | 1st PCR |
| 7 | | DNEPSPS1-R1 | GGGAGTGAGTGCATACTTGTT | |
| 8 | | EPPE-F2 | ACACTCTTTCCCTACACGACGCTCTTCCGATCTGGCAGTTTCCTGTCGGTAA | 2nd PCR |
| 9 | | nEPSPS1-R2 | GTGACTGGAGTTCAGACGTGTGCTCTTCCGATCTCAACACTGGAAAAAAAGAAGAAAAAA | |
| 10 | | EPSPS1-sF1.2 | TGGTGTCCTGAACAACTCATAC | Sanger sequencing |
| 11 | *SlCAB13* | UPCAB13-F1 | ATCGAGAGTCCTTACTCTTCACG | 1st PCR |
| 12 | | DNCAB13-R1 | CTCCAGGAAGCCCATTAATTCT | |
| 13 | | CAB13-F2 | ACACTCTTTCCCTACACGACGCTCTTCCGATCTCAGTTATTCTTTATATATAAGTATATAA | 2nd PCR |
| 14 | | CAB13-R2 | GTGACTGGAGTTCAGACGTGTGCTCTTCCGATCCTATATCCCTTAGGGGGTTAGC | |
| 15 | | SlCAB13-sF2 | GTGTGAACAACATATACAAGATTGG | Sanger sequencing |
| 16 | GT tool | RB-qF2 | CTCTTAGGTTTACCCGCCAATA | T-DNA |
| 17 | | GR-F1 | TTGAGATGAGCACTTGGGATAG | |
| 18 | | GR-F1 | TTGAGATGAGCACTTGGGATAG | Replicon |
| 19 | | pNOS-cR1 | AACGTGACTCCCTTAATTCTCC | |
| 20 | *SlGAPDH* | GAPDH-F1 | CCATAACCTAATTTCTCTCTC | Internal control |
| 21 | | GAPDH-R1 | GTCATGAGACCCTCAACAAT | |

**Table S7. Primers used for off-target analysis**

| No. | Potential off-target site | Chromosome location | Primer | Sequence (5’-3’) | Length (nt) | Note |
|---|---|---|---|---|---|---|
| 1 | EPSPS1-OT1 | SL4.0ch05-60580370..60584370 | EPSPS1-OT1-F1 | CCTACTCAGTGGGAGCTTAGT | 21 | PCR |
| 2 | | | EPSPS1-OT1-R1 | AATGCCCGCCCATAAAGAA | 19 | |
| 3 | | | EPSPS1-OT1-sF1 | TTCTTATGGTTGCACTTCTCA | 21 | Sequencing |
| 4 | CAB13-OT1 | SL4.0ch01-56416253..56420253 | CAB13-OT1-F1 | ATTTGGTGCCCTGGTCTTAG | 20 | PCR |
| 5 | | | CAB13-OT1-R1 | ATCCCGACCATCTTCGTATTG | 21 | |
| 6 | | | CAB13-OT1-sF1 | ACCTGCAGGCCTTTGATTT | 19 | Sequencing |
| 7 | CAB13-OT2 | SL4.0ch12-4304011..4308011 | CAB13-OT2-F1 | GATAGCCATCTCATTAATTTCAAGACC | 27 | PCR |
| 8 | | | CAB13-OT2-R1 | TCCACCGTCACTGAAGATTTG | 21 | |
| 9 | | | CAB13-OT2-sF1 | CTCCACTTTAGGCCTCGATTAT | 22 | Sequencing |

### Sequences used in the study

- **ttLbCas12a sequence**

ATGCCCAAGAAGAAGCGCAAGGTGGGACGCGTCTGCAGGATATCAAGCTTGCGGTACCGCGGCCCG
GGATCGCCACCATGAGCAAGCTGGAGAAGTTTACAAACTGCTACTCCCTGTCTAAGACCCTGAGGTTCA
AGGTAAAGCCTCGATTTTTGGGTTTAGGTGTCTGCTTATTAGAGTAAAAACACATCCTTTGAAATTGTTT
GTGGTCATTTGATTGTGCTCTTGATCCATTGAATTGCTGCAGGCCATCCCTGTGGGCAAGACCCAGGAGA
ACATCGACAATAAGCGGCTGCTGGTGGAGGACGAGAAGAGAGCCGAGGATTATAAGGGCGTGAAGAA
GCTGCTGGATCGCTACTATCTGTCTTTTATCAACGACGTGCTGCACAGCATCAAGCTGAAGAATCTGAAC
AATTACATCAGCCTGTTCCGGAAGAAAACCAGAACCGAGAAGGAGAATAAGGAGCTGGAGAACCTGGA
GATCAATCTGCGGAAGGAGATCGCCAAGGCCTTCAAGGGCAACGAGGGCTACAAGTCCCTGTTTAAGA
AGGATATCATCGAGACAATCCTGCCAGAGTTCCTGGACGATAAGGACGAGATCGCCCTGGTGAACAGCT
TCAATGGCTTTACCACAGCCTTCACCGGCTTCTTTAGAAACAGAGAGAATATGTTTTCCGAGGAGGCCAA
GAGCACATCCATCGCCTTCAGGTGTATCAACGAGAATCTGACCCGCTACATCTCTAATATGGACATCTTC
GAGAAGGTGGACGCCATCTTTGATAAGCACGAGGTGCAGGAGATCAAGGAGAAGATCCTGAACAGCG
ACTATGATGTGGAGGATTTCTTTGAGGGCGAGTTCTTTAACTTTGTGCTGACACAGGAGGGCATCGACG
TGTATAACGCCATCATCGGCGGCTTCGTGACCGAGAGCGGCGAGAAGATCAAGGGCCTGAACGAGTAC
ATCAACCTGTATAATCAGAAAACCAAGCAGAAGCTGCCTAAGTTTAAGCCACTGTATAAGCAGGTGCTG
AGCGATCGGGAGTCTCTGAGCTTCTACGGCGAGGGCTATACATCCGATGAGGAGGTGCTGGAGGTGTT
TAGAAACACCCTGAACAAGAACAGCGAGATCTTCAGCTCCATCAAGAAGCTGGAGAAGCTGTTCAAGAA
TTTTGACGAGTACTCTAGCGCCGGCATCTTTGTGAAGAACGGCCCCGCCATCAGCACAATCTCCAAGGAT
ATCTTCGGCGAGTGGAACGTGATCCGGGACAAGTGGAATGCCGAGTATGACGATATCCACCTGAAGAA
GAAGGCCGTGGTGACCGAGAAGTACGAGGACGATCGGAGAAAGTCCTTCAAGAAGATCGGCTCCTTTT
CTCTGGAGCAGCTGCAGGAGTACGCCGACGCCGATCTGTCTGTGGTGGAGAAGCTGAAGGAGATCATC
ATCCAGAAGGTGGATGAGATCTACAAGGTGTATGGCTCCTCTGAGAAGCTGTTCGACGCCGATTTTGTG
CTGGAGAAGAGCCTGAAGAAGAACGACGCCGTGGTGGCCATCATGAAGGACCTGCTGGATTCTGTGAA
GAGCTTCGAGAATTACATCAAGGCCTTCTTTGGCGAGGGCAAGGAGACAAACAGGGACGAGTCCTTCT
ATGGCGATTTTGTGCTGGCCTACGACATCCTGCTGAAGGTGGACCACATCTACGATGCCATCCGCAATTA
TGTGACCCAGAAGCCCTACTCTAAGGATAAGTTCAAGCTGTATTTTCAGAACCCTCAGTTCATGGGCGGC
TGGGACAAGGATAAGGAGACAGACTATCGGGCCACCATCCTGAGATACGGCTCCAAGTACTATCTGGC
CATCATGGATAAGAAGTACGCCAAGTGCCTGCAGAAGATCGACAAGGACGATGTGAACGGCAATTACG
AGAAGATCAACTATAAGCTGCTGCCCGGCCCTAATAAGATGCTGCCAAAGGTGTTCTTTTCTAAGAAGT
GGATGGCCTACTATAACCCCAGCGAGGACATCCAGAAGATCTACAAGAATGGCACATTCAAGAAGGGC
GATATGTTTAACCTGAATGACTGTCACAAGCTGATCGACTTCTTTAAGGATAGCATCTCCCGGTATCCAA
AGTGGTCCAATGCCTACGATTTCAACTTTTCTGAGACAGAGAAGTATAAGGACATCGCCGGCTTTTACAG
AGAGGTGGAGGAGCAGGGCTATAAGGTGAGCTTCGAGTCTGCCAGCAAGAAGGAGGTGGATAAGCTG
GTGGAGGAGGGCAAGCTGTATATGTTCCAGATCTATAACAAGGACTTTTCCGATAAGTCTCACGGCACA
CCCAATCTGCACACCATGTACTTCAAGCTGCTGTTTGACGAGAACAATCACGGACAGATCAGGCTGAGC

GGAGGAGCAGAGCTGTTCATGAGGCGCGCCTCCCTGAAGAAGGAGGAGCTGGTGGTGCACCCAGCCA
ACTCCCCTATCGCCAACAAGAATCCAGATAATCCCAAGAAAACCACAACCCTGTCCTACGACGTGTATAA
GGATAAGAGGTTTTCTGAGGACCAGTACGAGCTGCACATCCCAATCGCCATCAATAAGTGCCCCAAGAA
CATCTTCAAGATCAATACAGAGGTGCGCGTGCTGCTGAAGCACGACGATAACCCCTATGTGATCGGCAT
CGATAGGGGCGAGCGCAATCTGCTGTATATCGTGGTGGTGGACGGCAAGGGCAACATCGTGGAGCAGT
ATTCCCTGAACGAGATCATCAACAACTTCAACGGCATCAGGATCAAGACAGATTACCACTCTCTGCTGGA
CAAGAAGGAGAAGGAGAGGTTCGAGGCCCGCCAGAACTGGACCTCCATCGAGAATATCAAGGAGCTG
AAGGCCGGCTATATCTCTCAGGTGGTGCACAAGATCTGCGAGCTGGTGGAGAAGTACGATGCCGTGAT
CGCCCTGGAGGACCTGAACTCTGGCTTTAAGAATAGCCGCGTGAAGGTGGAGAAGCAGGTGTATCAGA
AGTTCGAGAAGATGCTGATCGATAAGCTGAACTACATGGTGGACAAGAAGTCTAATCCTTGTGCAACAG
GCGGCGCCCTGAAGGGCTATCAGATCACCAATAAGTTCGAGAGCTTTAAGTCCATGTCTACCCAGAACG
GCTTCATCTTTTACATCCCTGCCTGGCTGACATCCAAGATCGATCCATCTACCGGCTTTGTGAACCTGCTG
AAAACCAAGTATACCAGCATCGCCGATTCCAAGAAGTTCATCAGCTCCTTTGACAGGATCATGTACGTGC
CCGAGGAGGATCTGTTCGAGTTTGCCCTGGACTATAAGAACTTCTCTCGCACAGACGCCGATTACATCAA
GAAGTGGAAGCTGTACTCCTACGGCAACCGGATCAGAATCTTCCGGAATCCTAAGAAGAACAACGTGTT
CGACTGGGAGGAGGTGTGCCTGACCAGCGCCTATAAGGAGCTGTTCAACAAGTACGGCATCAATTATCA
GCAGGGCGATATCAGAGCCCTGCTGTGCGAGCAGTCCGACAAGGCCTTCTACTCTAGCTTTATGGCCCT
GATGAGCCTGATGCTGCAGATGCGGAACAGCATCACAGGCCGCACCGACGTGGATTTTCTGATCAGCCC
TGTGAAGAACTCCGACGGCATCTTCTACGATAGCCGGAACTATGAGGCCCAGGAGAATGCCATCCTGCC
AAAGAACGCCGACGCCAATGGCGCCTATAACATCGCCAGAAAGGTGCTGTGGGCCATCGGCCAGTTCA
AGAAGGCCGAGGACGAGAAGCTGGATAAGGTGAAGATCGCCATCTCTAACAAGGAGTGGCTGGAGTA
CGCCCAGACCAGCGTGAAGCACGCCTATCCCTATGACGTGCCCGATTATGCCAGCCTGGGCAGCGGCTC
CCCCAAGAAAAAACGCAAGGTGGAAGATCCTAAGAAAAAGCGGAAAGTGGACGGCATTGGTAGTGGG
AGCTAAGCTT

Purple font: SV40 NLS; blue font: LbCas12a; 1xHA tag; tan font: Trp1 intron; yellow highlighted font: D156R modification of the temperature tolerant LbCas12a; black font: linkers.

- **ttLbCas12a-10xSuntag sequence**

ATGCCCAAGAAGAAGCGCAAGGTGGGACGCGTCTGCAGGATATCAAGCTTGCGGTACCGCGGGCCCG
GGATCGCCACCATGAGCAAGCTGGAGAAGTTTACAAACTGCTACTCCCTGTCTAAGACCCTGAGGTTCA
AGGTAAAGCCTCGATTTTTGGGTTTAGGTGTCTGCTTATTAGAGTAAAAACACATCCTTTGAAATTGTTT
GTGGTCATTTGATTGTGCTCTTGATCCATTGAATTGCTGCAGGCCATCCCTGTGGGCAAGACCCAGGAGA
ACATCGACAATAAGCGGCTGCTGGTGGAGGACGAGAAGAGAGCCGAGGATTATAAGGGCGTGAAGAA
GCTGCTGGATCGCTACTATCTGTCTTTTATCAACGACGTGCTGCACAGCATCAAGCTGAAGAATCTGAAC
AATTACATCAGCCTGTTCCGGAAGAAAACCAGAACCGAGAAGGAGAATAAGGAGCTGGAGAACCTGGA
GATCAATCTGCGGAAGGAGATCGCCAAGGCCTTCAAGGGCAACGAGGGCTACAAGTCCCTGTTTAAGA
AGGATATCATCGAGACAATCCTGCCAGAGTTCCTGGACGATAAGGACGAGATCGCCCTGGTGAACAGCT
TCAATGGCTTTACCACAGCCTTCACCGGCTTCTTTAGAAACAGAGAGAATATGTTTTCCGAGGAGGCCAA
GAGCACATCCATCGCCTTCAGGTGTATCAACGAGAATCTGACCCGCTACATCTCTAATATGGACATCTTC

GAGAAGGTGGACGCCATCTTTGATAAGCACGAGGTGCAGGAGATCAAGGAGAAGATCCTGAACAGCG
ACTATGATGTGGAGGATTTCTTTGAGGGCGAGTTCTTTAACTTTGTGCTGACACAGGAGGGCATCGACG
TGTATAACGCCATCATCGGCGGCTTCGTGACCGAGAGCGGCGAGAAGATCAAGGGCCTGAACGAGTAC
ATCAACCTGTATAATCAGAAAACCAAGCAGAAGCTGCCTAAGTTTAAGCCACTGTATAAGCAGGTGCTG
AGCGATCGGGAGTCTCTGAGCTTCTACGGCGAGGGCTATACATCCGATGAGGAGGTGCTGGAGGTGTT
TAGAAACACCCTGAACAAGAACAGCGAGATCTTCAGCTCCATCAAGAAGCTGGAGAAGCTGTTCAAGAA
TTTTGACGAGTACTCTAGCGCCGGCATCTTTGTGAAGAACGGCCCCGCCATCAGCACAATCTCCAAGGAT
ATCTTCGGCGAGTGGAACGTGATCCGGGACAAGTGGAATGCCGAGTATGACGATATCCACCTGAAGAA
GAAGGCCGTGGTGACCGAGAAGTACGAGGACGATCGGAGAAAGTCCTTCAAGAAGATCGGCTCCTTTT
CTCTGGAGCAGCTGCAGGAGTACGCCGACGCCGATCTGTCTGTGGTGGAGAAGCTGAAGGAGATCATC
ATCCAGAAGGTGGATGAGATCTACAAGGTGTATGGCTCCTCTGAGAAGCTGTTCGACGCCGATTTTGTG
CTGGAGAAGAGCCTGAAGAAGAACGACGCCGTGGTGGCCATCATGAAGGACCTGCTGGATTCTGTGAA
GAGCTTCGAGAATTACATCAAGGCCTTCTTTGGCGAGGGCAAGGAGACAAACAGGGACGAGTCCTTCT
ATGGCGATTTTGTGCTGGCCTACGACATCCTGCTGAAGGTGGACCACATCTACGATGCCATCCGCAATTA
TGTGACCCAGAAGCCCTACTCTAAGGATAAGTTCAAGCTGTATTTTCAGAACCCTCAGTTCATGGGCGGC
TGGGACAAGGATAAGGAGACAGACTATCGGGCCACCATCCTGAGATACGGCTCCAAGTACTATCTGGC
CATCATGGATAAGAAGTACGCCAAGTGCCTGCAGAAGATCGACAAGGACGATGTGAACGGCAATTACG
AGAAGATCAACTATAAGCTGCTGCCCGGCCCTAATAAGATGCTGCCAAAGGTGTTCTTTTCTAAGAAGT
GGATGGCCTACTATAACCCCAGCGAGGACATCCAGAAGATCTACAAGAATGGCACATTCAAGAAGGGC
GATATGTTTAACCTGAATGACTGTCACAAGCTGATCGACTTCTTTAAGGATAGCATCTCCCGGTATCCAA
AGTGGTCCAATGCCTACGATTTCAACTTTTCTGAGACAGAGAAGTATAAGGACATCGCCGGCTTTTACAG
AGAGGTGGAGGAGCAGGGCTATAAGGTGAGCTTCGAGTCTGCCAGCAAGAAGGAGGTGGATAAGCTG
GTGGAGGAGGGCAAGCTGTATATGTTCCAGATCTATAACAAGGACTTTTCCGATAAGTCTCACGGCACA
CCCAATCTGCACACCATGTACTTCAAGCTGCTGTTTGACGAGAACAATCACGGACAGATCAGGCTGAGC
GGAGGAGCAGAGCTGTTCATGAGGCGCGCCTCCCTGAAGAAGGAGGAGCTGGTGGTGCACCCAGCCA
ACTCCCCTATCGCCAACAAGAATCCAGATAATCCCAAGAAAACCACAACCCTGTCCTACGACGTGTATAA
GGATAAGAGGTTTTCTGAGGACCAGTACGAGCTGCACATCCCAATCGCCATCAATAAGTGCCCCAAGAA
CATCTTCAAGATCAATACAGAGGTGCGCGTGCTGCTGAAGCACGACGATAACCCCTATGTGATCGGCAT
CGATAGGGGCGAGCGCAATCTGCTGTATATCGTGGTGGTGGACGGCAAGGGCAACATCGTGGAGCAGT
ATTCCCTGAACGAGATCATCAACAACTTCAACGGCATCAGGATCAAGACAGATTACCACTCTCTGCTGGA
CAAGAAGGAGAAGGAGAGGTTCGAGGCCCGCCAGAACTGGACCTCCATCGAGAATATCAAGGAGCTG
AAGGCCGGCTATATCTCTCAGGTGGTGCACAAGATCTGCGAGCTGGTGGAGAAGTACGATGCCGTGAT
CGCCCTGGAGGACCTGAACTCTGGCTTTAAGAATAGCCGCGTGAAGGTGGAGAAGCAGGTGTATCAGA
AGTTCGAGAAGATGCTGATCGATAAGCTGAACTACATGGTGGACAAGAAGTCTAATCCTTGTGCAACAG
GCGGCGCCCTGAAGGGCTATCAGATCACCAATAAGTTCGAGAGCTTTAAGTCCATGTCTACCCAGAACG
GCTTCATCTTTTACATCCCTGCCTGGCTGACATCCAAGATCGATCCATCTACCGGCTTTGTGAACCTGCTG
AAAACCAAGTATACCAGCATCGCCGATTCCAAGAAGTTCATCAGCTCCTTTGACAGGATCATGTACGTGC
CCGAGGAGGATCTGTTCGAGTTTGCCCTGGACTATAAGAACTTCTCTCGCACAGACGCCGATTACATCAA
GAAGTGGAAGCTGTACTCCTACGGCAACCGGATCAGAATCTTCCGGAATCCTAAGAAGAACAACGTGTT
CGACTGGGAGGAGGTGTGCCTGACCAGCGCCTATAAGGAGCTGTTCAACAAGTACGGCATCAATTATCA

GCAGGGCGATATCAGAGCCCTGCTGTGCGAGCAGTCCGACAAGGCCTTCTACTCTAGCTTTATGGCCCT
GATGAGCCTGATGCTGCAGATGCGGAACAGCATCACAGGCCGCACCGACGTGGATTTTCTGATCAGCCC
TGTGAAGAACTCCGACGGCATCTTCTACGATAGCCGGAACTATGAGGCCCAGGAGAATGCCATCCTGCC
AAAGAACGCCGACGCCAATGGCGCCTATAACATCGCCAGAAAGGTGCTGTGGGCCATCGGCCAGTTCA
AGAAGGCCGAGGACGAGAAGCTGGATAAGGTGAAGATCGCCATCTCTAACAAGGAGTGGCTGGAGTA
CGCCCAGACCAGCGTGAAGCACGCCTATCCCTATGACGTGCCCGATTATGCCAGCCTGGGCAGCGGCTC
CCCCAAGAAAAAACGCAAGGTGGAAGATCCTAAGAAAAAGCGGAAAGTGGACGGCATTGGTAGTGGG
AGCGgaggtgggggatctggttcgatgGAAGAACTTTTGAGCAAGAATTATCATCTTGAGAACGAAGTGGCTCG
TCTTAAGAAAGGTTCTGGCAGTGGAGAAGAACTGCTTTCAAAGAATTACCACCTGGAAAATGAGGTAGC
TAGACTGAAAAAGGGGAGCGGAAGTGGGGAGGAGTTGCTGAGCAAAAATTATCATTTGGAGAACGAA
GTAGCACGACTAAAGAAAGGGTCCGGATCGGGTGAGGAGTTACTCTCGAAAAATTATCATCTCGAAAAC
GAAGTGGCTCGGCTAAAAAAGGGCAGTGGTTCTGGAGAAGAGCTATTATCTAAAAACTACCACCTCGAA
AATGAGGTGGCACGCTTAAAAAAGGGAAGTGGCAGTGGTGAAGAGCTACTATCCAAGAATTATCATCT
TGAGAACGAGGTAGCGCGTTTGAAGAAGGGTTCCGGCTCAGGAGAGGAACTGCTCTCGAAGAACTATC
ATCTTGAAAATGAGGTCGCTCGATTAAAAAAGGGATCGGGCAGTGGTGAGGAACTACTTTCAAAGAATT
ACCACCTCGAAAACGAAGTAGCTCGATTAAAGAAAGGTTCAGGGTCGGGTGAAGAATTACTGAGTAAA
AATTATCATCTGGAAAATGAGGTAGCGAGACTAAAAAAGGGAGTGGTTCTGGCGAGGAATTGCTATC
GAAAAATTATCATCTTGAGAACGAAGTTGCTAGGCTCAAAAAGGGCTCAGGCTCAGGCACCGCGtaa

Purple font: SV40 NLS; blue font: LbCas12a; 1xHA tag; tan font: Trp1 intron; red font: GCN4_v4 (Suntag); yellow highlighted font: D156R modification of the temperature tolerant LbCas12a; black font: linkers.

##### ❖ hExo1a sequence (cloned from pmCherry-EXO1a plasmid (Addgene plasmid #111629))

ATGGGGATACAGGGATTGCTACAATTTATCAAAGAAGCTTCAGAACCCATCCATGTGAGGAAGTATAAA
GGGCAGGTAGTAGCTGTGGATACATATTGCTGGCTTCACAAAGGAGCTATTGCTTGTGCTGAAAAACTA
GCCAAAGGTGAACCTACTGATAGGTATGTAGGATTTTGTATGAAATTTGTAAATATGTTACTATCTCACG
GGATCAAGCCTATTCTCGTATTTGATGGATGTACTTTACCTTCTAAAAAGGAAGTAGAGAGATCTAGAAG
AGAAAGACGACAAGCCAATCTTCTTAAGGGAAAGCAACTTCTTCGTGAGGGGAAAGTCTCGGAAGCTC
GAGAGTGTTTCACCCGGTCTATCAATATCACACATGCCATGGCCCACAAAGTAATTAAAGCTGCCCGTTC
TCAGGGGGTAGATTGCCTCGTGGCTCCCTATGAAGCTGATGCGCAGTTGGCCTATCTTAACAAAGCGGG
AATTGTGCAAGCCATAATTACAGAGGACTCGGATCTCCTAGCTTTTGGCTGTAAAAAGGTAATTTTAAAG
ATGGACCAGTTTGGAAATGGACTTGAAATTGATCAAGCTCGGCTAGGAATGTGCAGACAGCTTGGGGA
TGTATTCACGGAAGAGAAGTTTCGTTACATGTGTATTCTTTCAGGTTGTGACTACCTGTCATCACTGCGT
GGGATTGGATTAGCAAAGGCATGCAAAGTCCTAAGACTAGCCAATAATCCAGATATAGTAAAGGTTATC
AAGAAAATTGGACATTATCTCAAGATGAATATCACGGTACCAGAGGATTACATCAACGGGTTTATTCGG
GCCAACAATACCTTCCTCTATCAGCTAGTTTTTGATCCCATCAAAAGGAAACTTATTCCTCTGAACGCCTA
TGAAGATGATGTTGATCCTGAAACACTAAGCTACGCTGGGCAATATGTTGATGATTCCATAGCTCTTCAA
ATAGCACTTGGAAATAAAGATATAAATACTTTTGAACAGATCGATGACTACAATCCAGACACTGCTATGC
CTGCCCATTCAAGA

❖ **T5 exonuclease sequence, synthesized**

ATGTCAAAGTCTTGGGGCAAGTTCATCGAGGAGGAGGAGGCCGAGATGGCGTCAAGGCGCAACCTCAT
GATTGTCGACGGCACCAATCTGGGCTTCCGGTTCAAGCACAACAATTCTAAGAAGCCTTTCGCCTCCAGC
TACGTGTCCACAATCCAGAGCCTCGCCAAGTCCTACAGCGCGCGCACCACAATTGTGCTGGGCGACAAG
GGCAAGTCAGTtTTCCGGCTGGAGCATCTGCCGGAGTACAAGGGCAACAGGGATGAGAAGTACGCACA
GAGGACCGAGGAGGAGAAGGCACTCGATGAGCAGTTCTTCGAGTACCTCAAGGACGCCTTCGAGCTGT
GCAAGACCACATTCCCAACCTTCACAATCAGGGGAGTGGAGGCAGACGATATGGCAGCGTACATCGTtA
AGCTCATTGGCCACCTGTACGATCATGTGTGGCTCATTTCCACAGACGGCGATTGGGACACCCTCCTGAC
AGACAAGGTtTCACGGTTCTCTTTCACCACACGGAGGGAGTACCACCTGAGGGATATGTACGAGCACCA
TAACGTGGACGATGTCGAGCAGTTCATCAGCCTCAAGGCCATTATGGGCGATCTGGGCGACAATATCAG
GGGAGTCGAGGGAATTGGAGCAAAGAGGGGCTACAACATCATTCGGGAGTTCGGCAATGTGCTCGATA
TCATTGACCAGCTCCCGCTGCCAGGCAAGCAGAAGTACATCCAGAACCTCAATGCGTCCGAGGAGCTCC
TGTTCCGCAATCTCATCCTGGTGGATCTGCCGACCTACTGCGTCGACGCAATTGCAGCAGTGGGACAGG
ATGTCCTCGACAAGTTCACAAAGGATATCCTGGAGATTGCGGAGCAG

❖ **hHE sequence (cloned from pCas9-HE-Geminin plasmid (Addgene plasmid #109402))**

ATGAACATCTCGGGAAGCAGCTGTGGAAGCCCTAACTCTGCAGATACATCTAGTGACTTTAAAGATCTG
TGGACCAAACTAAAAGAATGTCATGATAGAGAAGTACAAGGTTTACAAGTAAAAGTAACCAAGCTTAAG
CAAGAGCGAATCTTAGATGCACAAAGACTAGAAGAATTCTTCACCAAAAATCAACAGCTGAGGGAACA
GCAGAAAGTCCTTCATGAAGCCATTAAAGTTTTAGAAGATCGGTTAAGAGCAGGCTTATGTGATCGCTG
TGCAGTAACTGAAGAACATATGCGGAAAAAACAGCAAGAGTTTGAAAATATCCGGCAGCAGAATCTTA
AACTTATTACAGAACTTATGAATGAAAGGAATACTCTACAGGAAGAAAATAAAAAGCTTTCTGAACAAC
TCCAGCAGAAAATTGAGAATGATCAACAGCATCAAGCAGCTGAGCTTGAATGTGAGGAGGACGTTATTC
CAGATTCACCGATAACAGCCTTCTCATTTTCTGGCGTTAACCGGCTACGAAGAAAGGAGAACCCCCATGT
CCGATACATAGAACAAACACATACTAAATTGGAGCACTCTGTGTGTGCAAATGAAATGAGAAAAGTTTC
CAAGTCATCAACTCATCCACAACATAATCCTAATGAAAATGAAATTCTAGTAGCTGACACTTATGACCAA
AGTCAATCTCCAATGGCCAAAGCACATGGAACAAGCAGCTATACCCCTGATAAGTCATCTTTTAATTTAG
CTACAGTTGTTGCTGAAACACTTGGACTTGGTGTTCAAGAAGAATCTGAAACTCAAGGTCCCATGAGCCC
CCTTGGTGATGAGCTCTACCACTGTCTGGAAGGAAATCACAAGAAACAGCCTTTTGAG

❖ ***SlCtIP* sequence, cloned using tomato total cDNA**

ATGTATTCTGAGGCAAGAAGTGCTGCTGAAACTTCATGGAAAGAAAAGGAAAAAGATTTGCTTTTACAG
ATGGAAAAGATGCAATTTGAAAAGGAGAAGGTTCTTCAAGAAAACCAATACCTGAAAATGGAGAATGC
AAAGCTGTTGGATTCTGAACTGTCCTCAGCTAATCATGTTAAGGAGCTTCAAAATGAGTTGAAACAGAG
AACTAGTGAAATTAACGATTTAAGGGAAACAATCCGGAGATCATGTATTCTTCTTGAAACAAGTGCCCCA
CTTGTTTGCAAATATGAGAACCCACAGAGAGAACTTGAAGATAGAGTTCTCCTGCTAGTGAAAAAACAA
CAGATTTCAGAGCTTGAAGTTGGCAGGTTGCAGCTAGAACTCAAGAATAACTCGATAGAAGTTGACGGT
GGACTAGAGTTGCAAAATAAGCTTGTCCAATTGGCTCATTCAAAAACTTCTTCAGCAGCATATAAAGAAA
AGCAACTAAAAGAATATGAACAGAAaACAGCCAAACTTCTGGCTGAGCTTGAAACTACACGAAGAAGA

GTAGATGAACTCAACGAGGAGCTTAGAGAaAAGACAGCTGCAGTAGAAAAAGGACAGAAAATGCAAG
AAAATTTACTAAGTAAAGTTCGGTTGCTAGATGCAGAAATCATGAAGAATGATGATTTGTTGAATCAATA
CAAGAATGAGAAAGAACCACTTATGACCAAAGTAAAGAATCTGGAAACTGATGTTTATGATCTGCAGAA
AGAACTCCTGAATAAAAAGTCTGAAGTAGAGGAGGGAAGGAAGTTGCATGATCAATTACGTCAACAGA
TAGATTTGTATTCTTTAGAAAGGTCAAAGACAGGGCAGGAGTTGGAAGAGCTTGAGAAGGAGAACAAA
AAGGTACTAGCTAAATTGAGAGAATCAGAGGAAAAGATCGATAAGCTTCAGACAAATCTGAGAGAGAG
AGGCAAGGATTCCTCTGAGGGAATTAAATTACATGGGAAGTTACTTCATCTGATTCAAGCAAAAGAATC
TGAGTTGTTGGCCGAGAAGAAGAAAAGGAAGGATATGATTGCTTCTTATAAAAGTCTGAAATCACAGTA
CAACTTTCTATGTGCAAGATATGGTCTTACTTCAGAGAACATGCACCTACAAAGCAAATTATTAGAACAC
AGTGCATTACAGACTGACCAGAGTCCTCTAACTTCACGTGAGGTTGAAAATAAAGTTCCTCAGGCTTCTG
GTTTTGCCTGCAAAGTAATTAAGCAAGAAGAtAAACAGGAAGTCCAGGACGATGACCAAAGAGCTAGCC
TGATCCCAAGATCCAACTCCATCTCACCTCCAACTTCAAGTGCTTTTGTTGCTCCTAACATTCCAGCCAAT
GTCAGATCCTGCCCAACAGCTGGTACAAAGCGTTCTGTTTCTTATTGGAGGGATACCAGATCGCATCAGA
GTCGAGTTGGACCTGATCCTCATGATGATTTTCTCGATACTCCCCTGGAGAATATCAGAGGAAACTTAGG
AAAGGTGATGAAGGATGAGGTTGAGAATCACACAAAACCAAATTCGAAGGACAAAAAAATTGAgGACT
CAGATGATGAAACTCAGGACATGAATATTGATAGTGACCCTAAAAAGCAGGAAATGCTGCCTCCGGCCA
GGTCAGGTGCGACAGGCTTCAAATACATTGAACCTGTGAGAAAGAAAGCTGAACGAGAAAATTTAAAA
GGGGTCGAATGCCTGCAATGTAAGAAGTTTTATGATGCTGTTCATCCTGGTGAAGATAAGGAGTCCAAT
GGTAATAGGCAAAACCTACGCTGTGAGCATCATGATGGAGTTTCAAGGCATCGATACAGATATGCACCT
CCTTTAACTCCTGAAGGATTTTGGAATATTGGGTTTGAATCTGAAAAG

❖ ***SlKU80DN* sequence for fusions, cloned using tomato genomic DNA**

AAGCAACAAGACGCAGCAGATAAATTGGTTCAGATGTTGGATCTTGCACCACCTGGAAAACAGGAAGT
GTTATCACCTGACTTCACACCTAATCCTGTTCTAGAGCGTTACTACCGCTATCTTAACCTGAAGTCAAAGC
ACCCAGATGCAGCTGTTCCTCCACTTGATGAAACCCTCAGAAAGATAACAGAACCTGATGTTGAACTTCT
TTCTCAAAACAAGTCCATCATAGAGGAACTCCGTAGGTCTTTTGAACTAAAAGATAATCCAAAGCTGAAA
AAATCAGCAAGAAGAATAAAAGAaAGACCTTCAGGATCAGATGAGGAGATAGAAGAATTCAACAAAGA
TGCTGATGTCAAAGCTATAGACTCCATGGAATACTCAGCCAAAACAGAAGTTGAGAAAGTTGGAGATGT
TAATCCTGTCAAAGACTTTGAGGATATGCTGTCTCGAAGAGATAATCCAAAATGGATTAGTAAGGCCATT
CAGGATATGAAAAATAGGATCTTTGATCTCGTCGAAAATTCTTGTGACGGAGATACATTTCATAAAGCAT
TGCAATGTTTGGTGGCTCTACGCAAAGGTTGCATCCTTGAGCAGGAACCAAAGCAGTTCAATGATTTCCT
GTGCCACCTATCTAAATTTTGCCAAGAAAAAGACCTGAGAAGTTTCTGTCTATATCTCACATCTCATGAAA
TCACTTTGATAACCAAGGCAGAAGCTCCAGACAGTGAAATTTCAGAACATGAGGCTAGAAGCTTTATGG
TCAAGCCTGAACTTGACTCGCAAAATATGAAATCAGAGGCTAAAGCAGAGGATGATATTATGAGTATAT
ACCTGGGAGGGAAG

❖ ***SlKU80DN* sequence for overexpression in free form, cloned using tomato genomic DNA**

ATGAAGCAACAAGACGCAGCAGATAAATTGGTTCAGATGTTGGATCTTGCACCACCTGGAAAACAGGA
AGTGTTATCACCTGACTTCACACCTAATCCTGTTCTAGAGCGTTACTACCGCTATCTTAACCTGAAGTCAA
AGCACCCAGATGCAGCTGTTCCTCCACTTGATGAAACCCTCAGAAAGATAACAGAACCTGATGTTGAACT

TCTTTCTCAAAACAAGTCCATCATAGAGGAACTCCGTAGGTCTTTTGAACTAAAAGATAATCCAAAGCTG
AAAAAATCAGCAAGAAGAATAAAAGAaAGACCTTCAGGATCAGATGAGGAGATAGAAGAATTCAACAA
AGATGCTGATGTCAAAGCTATAGACTCCATGGAATACTCAGCCAAAACAGAAGTTGAGAAAGTTGGAG
ATGTTAATCCTGTCAAAGACTTTGAGGATATGCTGTCTCGAAGAGATAATCCAAAATGGATTAGTAAGG
CCATTCAGGATATGAAAAATAGGATCTTTGATCTCGTCGAAAATTCTTGTGACGGAGATACATTTCATAA
AGCATTGCAATGTTTGGTGGCTCTACGCAAAGGTTGCATCCTTGAGCAGGAACCAAAGCAGTTCAATGA
TTTCCTGTGCCACCTATCTAAATTTTGCCAAGAAAAAGACCTGAGAAGTTTCTGTCTATATCTCACATCTC
ATGAAATCACTTTGATAACCAAGGCAGAAGCTCCAGACAGTGAAATTTCAGAACATGAGGCTAGAAGCT
TTATGGTCAAGCCTGAACTTGACTCGCAAAATATGAAATCAGAGGCTAAAGCAGAGGATGATATTATGA
GTATATACCTGGGAGGGAAGggttcgccaaagaagaagcggaaggtctaa

Black color font: ku80DN; Purple font: SV40 NLS.

- **scfv-Ku80DN**

atgggccccgacatcgtgatgacccagagccccagcagcctgagcgccagcgtgggcgaccgcgtgaccatcacctgccgcagcagca
ccggcgccgtgaccaccagcaactacgccagctgggtgcaggagaagcccggcaagctgttcaagggcctgatcggcggcaccaacaa
ccgcgcccccggcgtgcccagccgcttcagcggcagcctgatcggcgacaaggccaccctgaccatcagcagcctgcagcccgaggact
tcgccacctacttctgcgccctgtggtacagcaaccactgggtgttcggccagggcaccaaggtggagctgaagcgcggcggcggcggc
agcggcggcggcggcagcggcggcggcggcagcagcggcggcggcagcgaggtgaagctgctggagagcggcggcggcctggtgca
gcccggcggcagcctgaagctgagctgcgccgtgagcggcttcagcctgaccgactacggcgtgaactgggtgcgccaggcccccggcc
gcggcctggagtggatcggcgtgatctggggcgacggcatcaccgactacaacagcgccctgaaggaccgcttcatcatcagcaagga
caacggcaagaacaccgtgtacctgcagatgagcaaggtgcgcagcgacgacaccgccctgtactactgcgtgaccggcctgttcgact
actggggccagggcaccctggtgaccgtgagcagctacccatacgatgttccagattacgctggtggaggcggaggttctgggggagga
ggtagtggcggtggtggttcaggaggcggcggaagcGGTTCGAAGCAACAAGACGCAGCAGATAAATTGGTTCAGA
TGTTGGATCTTGCACCACCTGGAAAACAGGAAGTGTTATCACCTGACTTCACACCTAATCCTGTTCTAGA
GCGTTACTACCGCTATCTTAACCTGAAGTCAAAGCACCCAGATGCAGCTGTTCCTCCACTTGATGAAACC
CTCAGAAAGATAACAGAACCTGATGTTGAACTTCTTTCTCAAAACAAGTCCATCATAGAGGAACTCCGTA
GGTCTTTTGAACTAAAAGATAATCCAAAGCTGAAAAAATCAGCAAGAAGAATAAAAGAaAGACCTTCAG
GATCAGATGAGGAGATAGAAGAATTCAACAAAGATGCTGATGTCAAAGCTATAGACTCCATGGAATACT
CAGCCAAAACAGAAGTTGAGAAAGTTGGAGATGTTAATCCTGTCAAAGACTTTGAGGATATGCTGTCTC
GAAGAGATAATCCAAAATGGATTAGTAAGGCCATTCAGGATATGAAAAATAGGATCTTTGATCTCGTCG
AAAATTCTTGTGACGGAGATACATTTCATAAAGCATTGCAATGTTTGGTGGCTCTACGCAAAGGTTGCAT
CCTTGAGCAGGAACCAAAGCAGTTCAATGATTTCCTGTGCCACCTATCTAAATTTTGCCAAGAAAAAGAC
CTGAGAAGTTTCTGTCTATATCTCACATCTCATGAAATCACTTTGATAACCAAGGCAGAAGCTCCAGACA
GTGAAATTTCAGAACATGAGGCTAGAAGCTTTATGGTCAAGCCTGAACTTGACTCGCAAAATATGAAAT
CAGAGGCTAAAGCAGAGGATGATATTATGAGTATATACCTGGGAGGGAAGggttcgccaaagaagaagcgga
aggtctaa

Orange color font: scfv sequence; red color font: HA tag; green color font: linker; black color font: ku80DN; Purple font: SV40 NLS.

##### ❖ *SlBRCA1* sequence, cloned using tomato total cDNA

ATGGCAGATATTTCGCACCTTGAAAGAATGGGAAGAGAGCTCAAATGCCCCATTTGCTTGAGTCTGTTC
AATTCTGCTGTTTCACTTACATGTAATCATGTATTTTGCAATTTATGTATTCAAACTGGTATGAAATCTGG
ATCCAATTGTCCGGTGTGCAAAGTTCCATTTCATCGCAGAGAAATCCGACCTGCTCTCCACATGGATAAC
TTGGTGAGCATCTATAAGAACATGGAAATTGCTTCAGGAGTCAATATGTTTCTCACTCAATCCAATCCTTC
CACAAAATTACCAGGGGAAAACACTCGATCTAATGGCGAAAAAGTTTGTGGATTCCAAGAGACACCTAA
AACTGTAACAGAAGCTCCAGCGACAGACAATCAGAAAAGAAAAAGAGGGAAAGGATCAAAAAGATCTT
CAGGGTGCAACAAAAAAATTTCCGGATCAAATCTTATTAGACCTTCTTTTCCAACAAAGAAGAGAGTACA
GGTGCCACAATATCCACCTTCAGAGACTCCACCACCTACAAAGTTAGTTGATGGAAATGGCAAATCCATC
ACCGATGAAGTTCAGAAACCATTGGTAATTGAGAGAGATAGGTCTATGCTAAATGAAAAAGGAGAACC
TGTGCTATCTCCATTCTTCTGGCTGAGAGAAGAGGATGTAGATAAATCAAGTCAGCAAACAGATGGGGA
CGTTATCATGGATACTCCTCCAGCTTTTCCATCCTTCAGCGATATGAAAGACTTGGATGATGAGGTTCACT
GCGAAATGACTCCAAAAAGTGGACCCTATGATGCAGCAAATGGAGCAGATCTTTTTGACAGTGAGATGT
TTGACTGGACACAAAGAGCTTGCTCCCCTGAACTTTGTTCTAGTCCCTTTAAGATGAAGCTTAAGGATAC
TATTGATTCTGCTGAAGCTCAGGAAAAGACTCAAGCTCACTCAGTTGAGGAAAGTGACATTAATGCATC
AGCAACTGAAAACAGAACAGCTGTGGAAAATGAAAAGGGTACTGATAAAGGACAGCTGAGTTCACCCG
CGATATTTTCTCCTGTAAATAAAACCACTAGTAGAAAAGACGTTGTTTGCAAGTCTAGCAGGAGCAAGTC
ATTGAGAAGTAGCCAGAAGAAACAAGGAAAAAACATAATTGGAGAATTATCAGAAGTTCATGATGCTTC
ACTAAAAGCAGCTGAAAACACTATGAAGAACAATCAGGATAATGCCAATGCATTTATCTCGAATAAGAA
GGATTCGAAAAACAAGAAAAAGGGTAGATCCTCTAGAAATGTCACTGAGTCAGTTGTAGAAGACATTTC
CACTTCATGTGGTGCTAAAAGACTTCGCAAGGGCAATAACTCAAAGTCCTTCAACTTGTCTACTATTGTG
AATCAGGAAAAACATAGTGAAGGAAGTGTCGAGACACTTGACTTGAAGACGCATAATATATTCCGCAAG
GGATCATTACGTGAACAAGCTAAAAACTGTTTTGGACCAAAAGGCGGAAAGAAAACTGCTGTGTGTAAC
ATTCCACAAATTCAAGATGAAGCTTTGCCTTTCGAATCAGCAAATAGATTGATTCCCATGGATAACAAGA
AACCTACTCGTAGTACAAAACTGAAGAAATGTGAGCTTGGTAGTGACAACAAGCTTCATGGGAAGAAAA
AAGTGAAGTTTTCTGATGATGGGCAGCTTGCTGACAAGGATAATATTACGCTTCAAAAGATTCAGAAGA
GAGTGCTCAATTCATTGGAAACAGATAAATCTGTTTTGAACTCGAATGATTCAGTTTTGCAGAAGTGTGA
GGCAAGCCAAAGCAAAATACAATGTGCTTTCTGCCGTTCAGCAGAGATATCTGAGGTTTCAGGAATCAT
GGTAAGCTACCTTAATGGAAACCCTGTCAAAGAAGATGTCAATGGAGCACCAGGTGTTATACATGTTCA
CAAATATTGTGCAGAATGGGCCCCTAATGTATATTTTGGAGACGATGATGTGGTCAACCTTGAATCTGAA
CTGAAGAGGAGTCGAAGGATCACTTGTTTTTTCTGTGGAGTAAAGGGGGCAGCTCTTGGGTGTTATGAG
ATGAGTTGTCGCAAAAGCTTTCATGTTCCTTGTGCCAAATTGACACCAGAGTGTAGATGGGATTCTGATA
ACTTTGTTATGTTATGCCCTTTGCATGCTAACTCTAAGTTGCCATCCGAGATCCCAGGAAAACAGACAAA
GATTGGAGATAGCATTAAAAGAAATTCTCGTATCCACCAACCCAATGTCTCAGCAACACCTGATAATGGT
GCTACTTTGCAGTGGAAGTCTCAGAAGAAGAATAAGAACTTAGTGCTCTGTTGTTCAGCTCTAACCGCA
GATGAAAAAGAGCTTGTTTCTAAATTAAAGAGGTTGTCTGGCGTGACAGTAGTCAAGAACTGGGACCTA
AGTGTCACCCATGTCATTGCTTCTACTGATGAGAAAGGAGCATGCCGAAGAACTCTCAAGTATTTGATG
GGTGTCTTGGCTGGGAAATGGATAATGAGCATCAACTGGATTATTGCTAGCTTGGAAGCCACAGAATAT
GTCGATGAACAACAATACGAGATTAAAATAGATACTCATGGCATTGTGGATGGGCCTAAGCTTGGAAGA

TTGAGGATTTTAAACAAGCAACCAAAGCTTTTCAATGGATACAAGTTCTTTTTTATGGGCGACTTTTTATC
TTCATACAAGAGCTACCTACATGACCTTGTTATTGCTGCTGGAGGAATTGTCCTTAACAGGAAGCCTATT
GCACTGGATCAGGAAATTCTTTCACCCGGATGCCCTCTACTGTTTGTAATTTATAGTCATGAACAACTTGA
TCAGTGCGAAGGAAGTGAAAAAATTTCAATTATAGCGCGCAGAAGATCTAATGCAGAAGTTTTGGCTAG
TTCAACTGGAGCTGTAGCTGCCAGCCACTCCTGGATTCTAAACTGCATTGCTGGCTCTAGGTTGTTGGAG
CTGGAATAG

- **hGeminin(1-110) sequence (cloned from pCas9-HE-Geminin plasmid (Addgene plasmid #109402))**

ATGAATCCCAGTATGAAGCAGAAACAAGAAGAAATCAAAGAGAATATAAAGAATAGTTCTGTCCCAAG
AAGAACTCTGAAGATGATTCAGCCTTCTGCATCTGGATCTCTTGTTGGAAGAGAAAATGAGCTGTCCGC
AGGCTTGTCCAAAAGGAAACATCGGAATGACCACTTAACATCTACAACTTCCAGCCCTGGGGTTATTGTC
CCAGAATCTAGTGAAAATAAAAATCTTGGAGGAGTCACCCAGGAGTCATTTGATCTTATGATTAAAGAA
AATCCATCCTCTCAGTATTGGAAGGAAGTGGCAGAAAAACGGAGAAAGGCGCTG

- **pAtHTR2 sequence , cloned using *Arabidopsis* genomic DNA**

GCAGGATGAGCTATAGGTGAAAGGTTGAAGATTTGAtGACCAAAGTTTTCAGTCTCTTTATTGTTTTATC
GTTTTCTTTTGAAGTATTTTAGCATCTAGTTTGTCGGAGATAGAAACAAGTTTTAGCAAAACTAAAAGGG
GGATACAATGTCTGTCAGATGCTCATTTTAGACTTTAAAAAACCATGAAGAAGTTTTGCAGTTGCTTCTCT
TTCTCTACTCTTGTTTCTCTGCTAAGCCAATGTCCGCCAAAATCGGATTCTGGTGTATTCGTTCACTGCTCC
CCACCCTTTGAGAATTTGTTGTTAGTCTCTTGTCAGTCAATACAGACCACCAAGTCCATAATCCAACAGTC
CAACGTCCCTGCAAGTTGATTGATACTAGAGTAGTAGTAATGTAGAATGTCTTTCACTTTCCAAACAATTT
CCACATTGTCTATACACTTATACTGGGTTCACTTCCATTGTGAcGACAGATGTTGAGATAACAAAAATTAT
ATAGAGATAATTGTACCCCAACCACAGATGCTTAGAACGCTAGAGAACGAATGCAGTGCAATCTTTGTTT
TACCTTAACCTTAGACAGCAATATTTTAATTCATGATGTCATTCTCAAAAGATGAAGTCACAATCTTTATA
CAAAACAGCGTCCTAGCATATTGAGATGGTATTTAAGCCCAGAAAAAGCTGATTGAGAACCAAAAAAAG
AAAAACAATTACGCAGGCCCTATTAAGTAATGGGCTTGGTTTTTTGGTAAAGATAATATATTTAGATTAT
TTCTTTTATTTCTATTGGTgTTCGTACCTTCACGCGGATCAGTGTTAGAAAAGAATGAAATCTGTCCGTAG
AATTGTATTAATCTACGGTTGCTGTGATTTCACGCGGATCGTGATAGAAACCTCTTATATCCGTTGATTAA
AAACCAATGAACGATCCAGATCTAATCCCGCTGAAAAAGAAAAGCGAAATATTTTCCCTCCCACTTCTTG
TAATTTCCAGACGATAAATATCCCCTTCCTCAATCGAAACAAACTATCCAGATCTCAACTTTCTCTCATCTT
CAAATTAAAATCAAACAGTTTCTTAATAACATTTTACTTC

- ***AtCDT1a (3C)* sequence, cloned using *Arabidopsis* total cDNAs**

CGGCTGATGAGTACTTCTCTGGCAGCTCGTCCGCTGAAAAGAAGCAACGGTCATACAAATCCAGATGAT
ATTTCTGCTGACCCACCGACTAAGCTCGTGAGGCGTTCCTTGTCTCTGAACTTTGATTCTTATCCTGAGGA
TGAGAGAACTATGGATTTCACAGATGATATACCCATTGATCAAGTACCAGAAGAAGATGTATCCAGCGA
TGATGAAATACTCAGCATACTTCCTGATAAACTTCGACATGCGATAAAAGAACAAGAGAGGAAAGCGAT
TGAGGATCAAAATCCAGCAATCTCTCTGGCAAAAAGAAGGCGAAAGATGATCGCTTGTCTACCTAAACT
TTTCAATGTCATCCATTACTTAATTCAATCGATAAGGCGTTGGGTTATCACAAAAGAAGAGCTTGTGCAC

AAGATCATTGCCGGTCACTCTGATATTACTGACAGAAAGGAAGTTGAAGAACAACTAATCTTACTGCAA
GAAATTGTCCCTGAATGGATGTCTGAGAAAAAATCATCAAGTGGAGATGTTCTAGTATGCATAAACAAG
TTGGCATCTCCTCTAACAATCCGATCACGTCTTGAAGAAGAAAACAAGCAAGAGATGGCTCCACTGCTTT
CT

❖ **Linker sequence for N-terminal fusion**

GGAATGATGGCCCCCAAGAAGAAGCGCAAGGTGGGACGCGTCTGCAGGATATCAAGCTTGCGGTACCG
CGGGCCCGGGATCGCCACC

Red font: SV40 NLS sequence

❖ **Linker sequence for C-terminal fusion**

AGCCTGGGCAGCGGCTCCCCCAAGAAAAAACGCAAGGTGGAAGATCCTAAGAAAAAGCGGAAAGTGG
ACGGCATTGGTAGTGGGAGCGGAGGTGGGGGATCTGGTTCG

Red font: SV40 NLS sequence

❖ **crRNA expression cassettes and donor sequences for *SlHKT1;2* locus**

- U6-crR1-2.20[HKT1;2] expression cassette:

TGATCAAAAGTCCCACATCGATCAGGTGATATATAGCAGCTTAGTTTATATAATGATAGAGTCGACA
TAGCGATTGtAATTTCTACTAAGTGTAGATACTATTCACCACAGTATCAATAATTTCTACTAAGTGTAGA
TCCTACAAATGAAAACATGATTTTTTT
Red font: AtU6 promoter; green font: LbCas12a scaffolds; light blue font: LbCas12a_gRNA1, 20nt; orange font: LbCas12a_gRNA2, 20nt; black G: transcription start; black TTTTTT: termination sequence.

❖ **crRNA expression cassettes and donor sequences for *SlEPSPS1* locus**

- Dual U6-crR1-2.23[EPSPS1] expression cassette:

TGATCAAAAGTCCCACATCGATCAGGTGATATATAGCAGCTTAGTTTATATAATGATAGAGTCGACA
TAGCGATTGtAATTTCTACTAAGTGTAGATGCGAAGTCCTCTACTGTCTCTAATTTCTACTAAGTG
TAGATCATATGGAGGAGGTCTGTGTTTTTTT
Red font: AtU6 promoter; green font: LbCas12a scaffolds; blue font: LbCas12a_gRNA1, 23nt; purple font: LbCas12a_gRNA2, 23nt; black G: transcription start; black TTTTTT: termination sequence.

❖ ***SlCAB13* donor sequence**

TGGAAAATGACTTTGATCGTATCAAGCCAATGAATAAAAAAAACTTTATAAATTCGATAATCTCCGAATATATC
CCGTGAATAACATATATTACTTGCGTTTCTCTTCATTATCTTTTTTTAAAAATATCTGTATATAGATATCTTCAGC
TAATTCTTTTAGTGAAATATGTTTATAGAGCAATGCACTTGTAATTATATATGGTCGTAAAAGTATGGTTTAAT
TCAACTCTTGGCTTGATTAAGCACAGAGGCATGAAGGGGACTCCTTGAGCACATAATTAATTAGTTGCATGGG
GTGCCATGAATTTGTTTCTTCAATTCAATTACTTTTTCAATTTCAATAACTCGTGGTTATAGAATAGAGTGTGAA

CAACATATACAAGACACGAGTTATAGAATAGAGTGTGAACAACATATACAAGATTGGAAATTGACAATTAAA
ATTCATTTAGTATAGCCACTAGAAAAAGTTGGCTAATCATACGCAGTAGACTCGAATTACTTTGGTGCGCATG
AGTCTAAACATTTTGGCAAGTTGGGAAACATGAAACAACGCTTACAATAAAGTGCATCGATATAAAAGTATGT
TTAAATACCATAAAATTTCATATTTAAATTTTAGCCTGACAAAATAGTAATGAGCGATTTATTCTCATCTATCAA
AATCTTAATAAATAAAGTTATTTAATTTTTATATTAATTACAAATTGATTCGTACGCCACAATAATAAAATAAAA
TCCAAAAAGCTTGACCAATGACCATATATTACATTGTGTACTACTATATGTTAGCAAGATAAAACTCTGGATAG
AAGGATATGACATGAAGATACTCAAAATCTACTCAACACTTCTCATTGGTTGATACAAAAATTTCAAAACTATT
TCACACCACAATTTTATAAAACACAAGAACCAGAATTGTTAAATGGGGCCCCTCCAGATATTTTTCCAACAGTT
ATTCTTTATATATAAGTATATAAATTGTGCAAGAAACTTAGTGAAAATTGTGAGGTCCAATTAAcTAgAAGAGA
AATTAAGCAATAGAGAAAATGGCAGCAACAGGTAGCTCAGCCACAGTTGTTAGAGCAACTCCATTTTTGGGC
CAaACCAAgTATGCTAACCCCCTAAGGGATATAGTTCCTATGGGCTCTGCCAGATTCACCATGGTAACTTTTTA
TTTTTCTTTTAAGTGTTTTTATAATTGAGTTCGATATTTAATATTATTGAACTTCAAATAAATTTATGATTTGTGT
AAAAAGATACTGAACCTGTTATGTTTTATCAGAGTAATGATTTGTGGTATGGACCTGACCGTGTCAAGTACTT
GGGACCATTTTCTGCTCAAACTCCTTCATACTTGACTGGAGAATTCCCTGGTGATTACGGATGGGATACTGCTG
GTTTATCTGCTGATCCCGAGGCCTTTGCTAAGAACAGAGCTCTTGAGGTACTTTCACTTTTTCTTTTACTAGTAG
TAAGTTTGTTTAAAATAAGTAACAGACATTTGCGTGACTAAAATTCATTTTTAGCTAAAAAAGTTTGAAAAATA
AAAAAATTATCTTTTTCGTTTTATTTTATGTGACACTATTTGACTTGACACAGTGTTTTAAGAAGAAGAAAAAAT
GGTCTAAACCATTTCTTAGATATTGTGTAGTTGAAAATCATCTCATTAAGGATAAAAGAGGTATTTTAAAGCTA
ACTTGTTTCTAATTATAGTGATATTTTTTTTTAACGGATAAAAATGAAAGAATATCACATAAAATAGGACAGAG
GAATATCATTTTTGTTGGGACAAAACAAAAAAGTGAAGCACGCCATAGAAAATGTACAGATTTGAGACTCGT
AACGATTAATTTAGTATTAAGACATAGTTGATGGATTGATATGAATTGAAATTGTGTTAATTATTCTAGGTTAT
CCATGGGAGATGGGCCATGCTTGGAGCTTTTGGTTGCATTACACCAGAAGTTCTTGAAAAATGGGTAAAAGT
GGACTTCAAAGAACCAGTATGGTTCAAAGCTGGAGCCCAGATCTTCAGTGAAGGTGGGCTGGACTATTTGGG
CAACCCAAACCTTGTCCATGCTCAGAGC

Blue font: LbCas12a_gR1.CAB13 binding site; Purple font: LbCas12a_gR2.CAB13 binding site; yellow highlighted font: modified PAM and LbCpf1 gRNA core region to prevent recut after HDR editing; red font: AAATTGTGA inserted sequence.

- **crRNA expression cassettes and donor sequences for *SlCAB13* locus**
    - Dual U6-crR1-2.20$^{CAB13}$ expression cassette:

TGATCAAAAGTCCCACATCGATCAGGTGATATATAGCAGCTTAGTTTATATAATGATAGAGTCGACA
TAGCGATTGtAATTTCTACTAAGTGTAGATATTAATTGGACCTCACTAAGTAATTTCTACTAAGTG
TAGATGTCTGGCCCAAAAATGGAGTTTTTTT

Red font: AtU6 promoter; green font: LbCas12a scaffolds; blue font: LbCas12a_gR1.CAB13, 20nt; purple font: LbCas12a_gR2.CAB13, 20nt; black G: transcription start; black TTTTTT: termination sequence.
